# Unraveling the Transcriptional Landscape within a Minimized Bacterium via Comparative Analysis

**DOI:** 10.1101/2025.10.23.681674

**Authors:** Troy A. Brier, Jay E. Cournoyer, Benjamin R. Gilbert, Sage A. Glass, Yang-le Gao, Yanbao Yu, Zane R. Thornburg, Karrie Goglin, Gavin John, Tiana Mamaghani, Sujata Shivakumar, Christopher J. Fields, John I. Glass, Angad P. Mehta, Zaida Luthey-Schulten

**Affiliations:** Department of Chemistry, University of Illinois Urbana-Champaign, Urbana, IL 61801, USA; J. Craig Venter Institute, La Jolla, CA 92037, USA; Department of Chemistry and Biochemistry, Univeristy of Delaware, Newark, DE 19716, USA; Cancer Center at Illinois, University of Illinois at Urbana–Champaign, Urbana, IL 61801, USA; Beckman Institute for Advanced Science and Technology, University of Illinois at Urbana–Champaign, Urbana, IL 61801, USA; Genomatica, San Diego, CA 92121, USA; Roy J. Carver Biotechnology Center, Urbana, IL 61801, USA; Carl R. Woese Institute for Genomic Biology, Urbana, IL 61801, USA; Department of Physics, University of Illinois Urbana-Champaign, IL 61801, USA; NSF Science and Technology Center for Quantitative Cell Biology, Beckman Institute, Urbana, IL 61801, USA

**Keywords:** RNA sequencing, RNA isoforms, JCVI-syn3A, minimized bacteria, transcriptomics, genome sequence architecture

## Abstract

Stochastic nature of gene expression leads to the complex formation of the bacterial transcriptome and proteome. In contrast to typical transcriptome studies, we employ a near wild-type, Syn1.0, of the naturally genome-reduced *Mycoplasmas*, and the dramatically further genome-reduced JCVI-syn3A thus avoiding additional contributions from many non-essential cellular functions. To aid in profiling the transcriptional landscape within these bacteria, we present a bioinformatic analysis of the genetic sequence motifs implicated in modulating the stochastic gene expression events, coupled with genome-wide short-read (Illumina) and long-read (Oxford Nanopore Technologies and Pacfic Biosciences) RNA sequencing. The bioinformatic analysis coupled with information from structural studies assigns strengths of the Shine-Dalgarno signatures and identifies both transcription initiation and termination sites, leading to predictions of RNA isoforms in Syn1.0 (and related organisms). The long-read and short-read RNA sequencing characterized the predicted transcriptional activity, and the long-read methods provide direct insight into the RNA isoform complexity within Syn1.0. Comparison of the RNA sequencing results with that of the bioinformatic analysis highlights the inability of bioinformatics alone to capture the results of bacterial transcription without including effects of RNA degradation. This study emphasizes the need for comparative analysis and potential dangers of genome reduction, exemplified through the discovery of altered gene expression patterns of JCVI-syn1.0 and JCVI-syn3A, achieved via the union of our transcriptome study with their proteomics data. Analysis of the transcriptomics data sets through a Jupyter notebook allows any genomic region to be easily examined.

**Table of Content Image:** 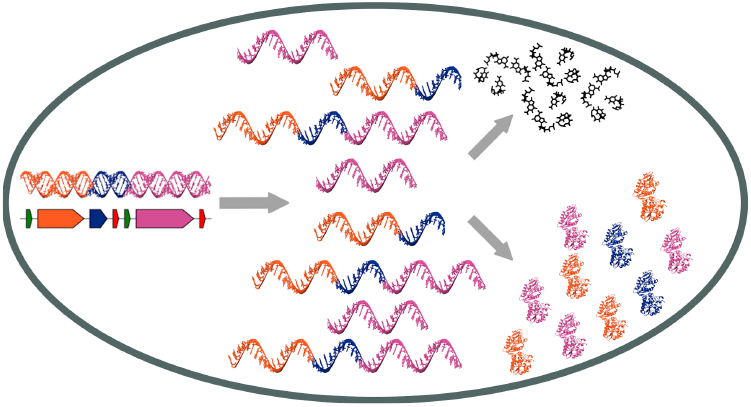

## 1. Introduction

Genetically minimized bacteria provide a unique opportunity to elucidate the inherent phenomena governing bacterial life (Morowitz, 1984; Lachance et al., 2019). JCVI-syn1.0 (Syn1.0) is a strain of the caprine pathogen Mycoplasma mycoides subspecies capri that has a synthetic genome. In synthesizing the Syn1.0 genome which has roughly 910 genes, 14 wild-type *M. mycoides* genes were either deleted or disrupted and five non-native genes required for the cloning procedure as well as four watermarks were incorporated (Gibson et al., 2010). Based on annotation of those genes, their absence or alteration is unlikely to affect transcription. *Mycoplasmas* evolved from low G+C% Gram-positive bacteria similar to *Bacillus subtilis* or *Staphylococcus aureus* through a process of massive gene loss beginning about 600 million years ago (Maniloff et al., 1992). This was possible because as obligate parasites, *Mycoplasmas* lived in extremely stable environments such as mammalian respiratory and urogenital tracts. Wild-type *M. mycoides capri* lives only in goat upper respiratory tracts, which are ∼30 °C, and lower respiratory tracts, which are ∼39 °C. An analysis of adenine methylation using Pacific Biosciences (PacBio) genome sequencing of a very similar goat pathogen, *Mycoplasma capricolum* showed 15 GATGA sites with methylation of the first adenine when the bacteria were grown at ∼37 °C, and a different 308 GATGA sites with that methylation in DNA from bacteria grown at ∼30 °C (data not shown). So these *Mycoplasmas* likely use DNA methylation to control expression of some genes. Even after millions of years of genome minimization, *Mycoplasmas* still actively regulate gene transcription. It is noteworthy that Syn1.0 encodes only 24 methyltransferase proteins (**Table** S10), while *B. subtilis*, a phylogenetically similar species that can live in many different environments, encodes 145 (Elfmann et al., 2024). The minimization of *Mycoplasma* ancestors similar to *B. subtilis* likely resulted in the loss of large numbers of genes that modified bacterial DNA and RNA that were not needed for life in the very stable environmental niches that *Mycoplasmas* occupy.

While the primary focus of this study is Syn1.0, we analyzed a more minimized version of *M. mycoides* called JCVI-syn3A (Syn3A) (Breuer et al., 2019). Its additional genome reduction to 493 genes was not an evolutionary process. The genome of Syn3A was designed to comprise only the minimum set of genes needed for rapid growth in laboratory media at ∼37 °C (∼ 80 of the 493 genes are non-essential but were retained to keep operons intact and for other reasons not related to viability). Most phenotypic characteristics of Syn3A and Syn1.0 are similar. Syn3A divides more slowly than Syn1.0 (2 hours between cell divisions as opposed to 1 hour) (Breuer et al., 2019). In recent years extensive experimental characterization of Syn1.0 and Syn3A has been reported (Hutchison et al., 2016; Breuer et al., 2019; Mariscal et al., 2018; Hutchison et al., 2019; Gilbert et al., 2021; Pelletier et al., 2021; Haas et al., 2022; Thornburg et al., 2022; Moger-Reischer et al., 2023; Sandberg et al., 2023; Bittencourt et al., 2024; Justice et al., 2024; Safronova et al., 2024). To briefly summarize, the initial genome synthesis and transplantation resulted in JCVI-syn1.0, a near wild-type version of *Mycoplasma mycoides* with a chemically synthesized genome of 911 genes capable of growth and division (Gibson et al., 2010). Genome reduction resulted in JCVI-syn3.0, which to date has the least number of genes observed in any bacteria (Hutchison et al., 2016). However, its inconsistent morphology motivated the re-introduction of approximately 19 genes resulting in the more robust, JCVI-syn3A with 493 genes (Breuer et al., 2019; Pelletier et al., 2021). Syn3A was constructed to only have primarily essential genes leading to few remaining identified regulatory elements (see **Table** S1-**Table** S2). Like other *Mycoplasmas*, Syn1.0 and Syn3A encode only a single sigma factor. Most bacteria encode multiple sigma factors to regulate their gene expression. Importantly, Syn3A, whose genome was designed to live in a precise stress free laboratory environment, lacks 6 of the 18 annotated transcriptional regulatory protein encoding genes present in Syn1.0. The transcriptional regulators remaining may be essential for control of the regulation of genes during a bacterial cell cycle and may be required in most bacteria. Contrary to their reduced nature, the minimized bacteria still have the genes needed to perform the fundamental bacterial cellular functions common to all life: having a cellular envelope, acquiring molecular building blocks, genome replication, transcription of genes to make RNA, and translation of messenger RNA (mRNA) into proteins.

Consequently, we hypothesize that *Mycoplasmas* like Syn1.0, which are the products of millions of years of evolution that left them with highly minimized genomes, offer model systems to investigate how gene expression is controlled in the primordial core sets of genes present in essentially all bacteria. Furthermore, Syn3A, which did not evolve to live a a goat parasite, but was designed to grow rapidly and with consistent morphology in an unchanging laboratory environment, is an even better chassis to for both experimental and bioinformatic investigation of transcription and translation in essential bacterial genes.

A detailed understanding of how bacterial cells control the transcription of RNA is critical to understanding how cells work. Most bacteria are able to function in environments that change as a result of being able to adapt gene expression to turn genes on an off to suit different circumstances. This adaptability is possible because most bacteria have large enough sets of genes that allows survival in unstable or multiple niches. However there are some bacteria that evolved to live in extremely stable environments and as a result of that stability were able to lose large fractions of their genomes. Through analyzing transcription and proteomics data from minimal bacteria as opposed to a more typical model organism, such as *Escherichia coli* or *Bacillus subtilis*, we can better understand the fundamental principles of gene expression in the core elements common to bacterial cells. We use bioinformatics analysis, RNA sequencing, and proteomics to analyze transcription in two bacterial strains whose genomes have undergone massive gene loss.

The bio-informatic approach identifies the genetic motifs capable of modulating gene expression within JCVI-syn1.0 (and related organisms like Syn3A). Genome sequence architecture or the arrangement of the genetic motifs along the chromosome is highly conserved across bacterial life (Tamames, 2001; Fondi et al., 2022). From a genetic motif perspective, the circular chromosome of Syn1.0 (and Syn3A) is highly organized like many other bacteria. The local arrangement of sequence motifs important for controlling transcription and translation such as promoters, gene open reading frames (ORFs), Shine-Dalgarno sequences, transcription start sites (TSSs), and intrinsic termination loops exhibit some variation as shown in **Fig. 1A**, giving rise to different patterns of gene expression. The transcription units (TUs), defined to be the regions of the DNA that are transcribed during a single transcriptional event, may contain multiple genes, and a single gene may belong to multiple TUs as has been observed in *Mesoplasma florum* (Matteau et al., 2020). TU production begins with the initial binding of RNA polymerase (RNAP) at the promoter followed by transcription initiation at the TSS and finally ends with the halting of the RNAP due to the formation of an intrinsic termination loop encoded within the DNA sequence (de Hoon et al., 2005; Matteau et al., 2020). The usage of an intrinsic termination loop as a mechanism to halt transcription is typical for Gram-positive bacteria instead of the Gram-negative mechanism of Rho-dependent transcription termination (de Hoon et al., 2005). Transcription of multiple bacterial genes gives rise to a single polycistronic mRNA and may result in multiple mRNA isoforms upon subsequent processing such as cleavage into smaller transcripts by ribonucleases, primarily occurring at degradosomes. The possible RNA isoform outcomes for bacteria have been recently reviewed in Taggart et al. (2021) (See **Fig. 1B**). Recent stochastic simulations of a smal bacteriophage genome (10 genes) indicate that the local TU architecture plays a role in the observed gene expression pattern, and manipulation of the genome sequence architecture can produce pre-determined gene expression patterns (Shah et al., 2022).

**Figure 1.**
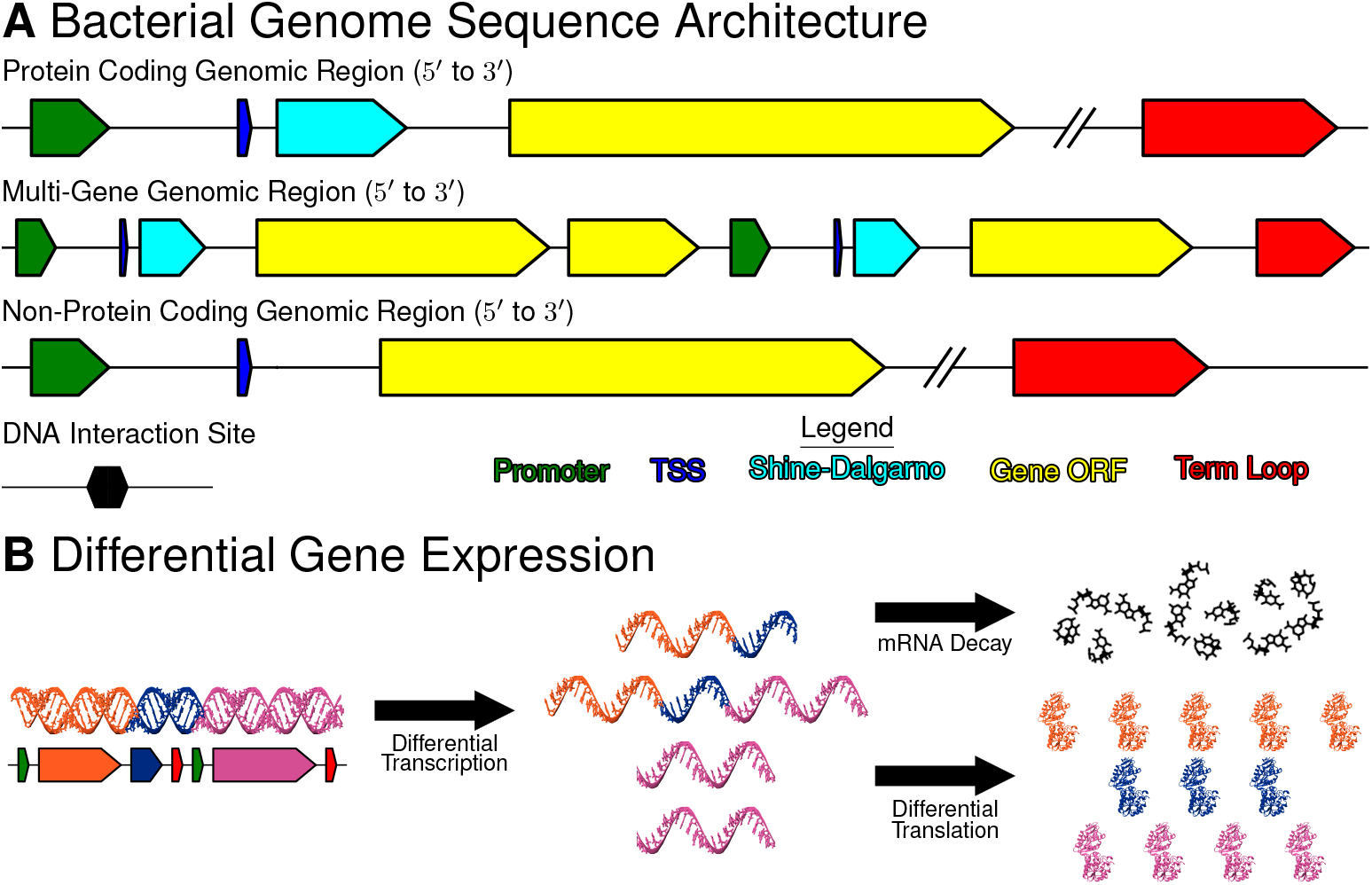
Summary of Bacterial Transcriptome Formation or Known effectors of Bacterial Transcriptome Formation: **(A)** Local genome sequence architecture varies depending whether the genomic region codes for a protein or not. Gene encoding regions contain the genetic information for transcription and eventual translation of a gene. Possible genetic motifs are gene open-reading frame (yellow), promoters (green), Shine-Dalgarno (cyan), transcription start sites (TSS, blue), and intrinsic termination loops (red). DNA interaction sites (black) are regions of the DNA where a molecule can bind causing some functional change in the DNA such as riboswitches, DnaA binding signature, transcription factor binding sites, nucleoid associated proteins binding sites, etc. Each motif has a role in modulating the efficacy of a step (translation or transcription) within the central dogma. **(B)** Differential gene expression impacts the transcriptome and proteome abundances. The genome sequence architecture of the DNA (left) varies initiation and termination events in transcription generating RNA isoforms (middle). Each RNA isoform has a unique affinity towards post-transcriptional processing such as differential translation or RNA decay. Protein expression patterns arise from the differential translation of the RNA isoforms (right). In the naive example of three genes (orange, blue, pink), differential gene expression produces a 1:1:1 ratio at the genome level, a 2:2:3 ratio at the transcriptome level, and a 5:3:4 ratio at the proteome level.

Differential gene expression at the translational level results from variation in the interactions between RNA transcripts and the ribosomes. This can occur because of folding states in a transcript altering ribosome accessibility and regulatory motifs at the start of the transcript, known as Shine-Dalgarno (SD) sequences, modifying translation initiation. The SDs detectable within the genome are located at the 5*′* end of a mRNA and interact via RNA hybridization with the anti-Shine-Dalgarno sequences (aSD), the 3^*′*^ end of the 16S rRNA (Shine and Dalgarno, 1974). The SD-aSD interaction has been observed structurally in *Thermus thermophilus* (Kaminishi et al., 2007) and has been characterized in many other organisms (Hockenberry et al., 2017, Hockenberry et al., 2018; Matteau et al., 2020; Saito et al., 2020; Webster et al., 2024). Other post-transcriptional processes, including, but not limited to translational regulation (Walker et al., 1984; Zhao et al., 2019) and interactions with small RNAs Bianchi et al. (2020) will also impact the eventual expression of the gene products encoded by these transcripts at the translational level.

The bioinformatics analysis enabled predictions about the transcriptome landscape. To validate these predictions we used three RNA sequencing methods. Advances in RNA sequencing (RNAseq) has greatly improved the ability to understand the transcriptome complexity (Soneson et al., 2019). Illumina-based sequencing is currently the most common technique used for capturing gene expression information due to robustness, cost, and throughput. Illumina instruments are capable of generating billions of reads per flow cell, and the technology for generation of RNA-based sequencing data has undergone considerable refinement since initial publications using the technique (Mortazavi et al., 2008; Nagalakshmi et al., 2008; Wilhelm et al., 2008). Sample preparation of gene expression libraries requires several intermediary steps prior to sequencing: (i) fragmentation of RNA, conversion of isolated mRNA fragments to double-stranded complementary DNA (cDNA) through reverse transcription, (iii) adapter ligation, (iv) PCR amplification, and finally, (v) size selection (Stark et al., 2019). Fragments may be sequenced either from one end (single-end) or both ends (paired-end), generally with read lengths of 150-250 nt depending on the instrument used. Though the reads are short, they are also well over 99% base accuracy, particularly using the latest chemistries and instrumentation (Illumina, 2011). Pacific Bioscience (PacBio) sequencing, in comparison, requires similar initial library preparation steps from mRNA to cDNA but is capable of sequencing full-length cDNA through a standard protocol, termed Iso-Seq™, which incorporates SMRTBell adapters allowing for circular consensus sequencing (CCS, otherwise known as HiFi sequencing) (Wenger et al., 2019). Improved instrumentation such as the PacBio Revio and newer techniques like Kinnex concatamerization (https://www.pacb.com/technology/kinnex/) ensure high yield and highly accurate sequence data of full length isoforms of up to 20 kb, though longer transcripts may be truncated or missing if size selection is performed. PacBio sequencing requires significant RNA concentrations for sequencing, thus PCR amplification is commonly applied during library preparation, and some size selection may also be needed. In addition, PacBio has a known bias towards sequencing smaller fragments which can make quantification of expression signal more challenging (Rhoads and Au, 2015); protocols are available to help limit this but require care with library preparation and sample loading. Kinnex concatamerization may effectively address this, but as the technique is quite new at this no independent studies are available that compare these to paired Illumina RNA-Seq libraries. Illumina sequencing will only measure short fragments of RNA, failing to capture the full complexity of transcripts present within a bacterial cell. Both Illumina and PacBio Iso-Seq additionally require several additional steps for library preparation including conversion to cDNA, PCR amplification, and size selection. However, Oxford Nanopore Technologies (ONT) have developed a method to capture long-read transcripts directly using a nanopore within a membrane. RNA transcripts are passed through the nanopore (3^*′*^ to 5^*′*^), and sequenced based on the perturbations of an applied electric current (Garalde et al., 2018). We elected to also sequence our samples using ONT because of its ability to read full length native transcripts without the need to convert the RNA to cDNA, which can perturb the native state causing the loss of information (Soneson et al., 2019) The technique provides the ability quantify transcripts as well as characterize co-expression of genes, highlight anti-sense / intergenic transcription, and possibly detect some common modifications to bases in the transcripts such as 6-methyl-adenosine (Liu et al., 2019). The RNAseq methods are further described and reviewed in Stark et al. (2019); Satam et al. (2023).

Comparison of motifs uncovered from the long-read RNAseq data with the bio-informatics predictions suggested a relatively high degree of co-expression among certain genes, potentially providing a mechanism used by these minimal cells to create proteins at functional stoichiometries. Discrepancies between the two methods highlighted previously unidentified regulatory motifs within the Syn1.0 genome. Additional analysis and comparison of the three RNAseq methods unveiled previously unknown transcriptional activity within the bacteria, and motivates the need for further investigation. Finally, using Syn1.0 and Syn3A results in conjunction, we highlighted some unintended consequences of the synthetic genome reduction between the two organisms. Our work presents an effort to better understand the composition and factors influencing the transcriptome within bacteria. This study provides a foundation for the transcriptome profiling needed to enhance kinetic models of stochastic gene expression such as Thornburg et al. (2019); Macklin et al. (2020); Fischer et al. (2021); Thornburg et al. (2022). It also emphasizes the need for comparative analysis (short- and long-read sequencing) and the potential dangers of genome reduction, exemplified through the discovery of altered gene expression patterns between JCVI-syn1.0 and JCVI-syn3Adetected via the union of our transcriptome study with proteomics data. To allow the analysis of transcriptomics data associated with any genomic region of interest, we provide a Jupyter notebook as we can only present a few examples in this article. This work represents a first of its kind on the transcriptome within the minimized bacteria.

## 2. Results and Discussion

### 2.1. *De Novo* Genetic Motifs and Transcription Unit Identification

Given the critical role genetic motifs (gene ORF, Shine-Dalgarno, promoter, and intrinsic termination loop) have within gene expression, we set out to identify them within the genome of JCVI-syn1.0 and related organisms: JCVI-syn3A, *Mycoplasma mycoides* subsp. *capri, Mesoplasma florum*, and *Mycoplasma pneumoniae*. The general approach taken to identify the motifs was to search through the genome using template sequence according to a well-defined consensus sequence to locate and characterize each motif (more details provided in **Section 4.1**). Reference genome sequences were obtained from the NCBI GenBank database (Sayers et al., 2021) for each organism of interest in the study. Each NCBI GenBank file lists all gene ORFs within the organism including information about the strand, genomic coordinates (position), coding type (protein coding or non-coding), and gene product. JCVI-syn1.0 and the other organisms have multiple forms of RNA including small RNA (sRNA) and transfer-messenger RNA (tmRNA), which have regulatory functionality within the cell (Burgos et al., 2021; Boutet et al., 2022). The presence of these genes within Syn1.0 and more so the minimal cell Syn3A indicate their regulatory roles are an essential function for bacterial cellular life (Hutchison et al., 2016; Breuer et al., 2019). Additional regulatory regions (referred to as *misc_features* or *misc_binding* within the NCBI GenBank entries) involved in transcription were also reported for each of the organisms examined in this study except for *M. pneumoniae*. These regulatory regions belong to the general category of DNA interaction sites (**Fig. 1A**). The lack of such regions in *M. pneumoniae* is suspected to be due to a lack of reporting of these motifs within its GenBank entry, but could be a result of differences in transcriptional activity between the organisms.

Additional motifs, which were promoter, Shine-Dalgarno, and intrinsic termination loops, were identified using the genomic sequence and position. Using a gene ORF as a reference point, SD were identified upstream within a ∼30 nucleotide region. Promoter were identified using the SD position and searching a region, this ends up being between 13 and 56 nucleotides upstream of the gene ORF. Termination loops were identified within the region downstream of each gene ORF until the next gene ORF or 500 nucleotides, whichever is reached first. Complete protocol details used for each motif are described in Materials and Methods, **Section 4.1. Table 1** reports a summary of the findings. We detected many instances of the motifs throughout the genome, however it was infrequent that a gene ORF would have each motif within immediate proximity. Shine-Dalgarno results (**Figs**. S3-S7) show the expected relationship between the SD and the 3^*′*^ 16S rRNA (aSD). Functionality was assigned using binding energy assuming a spontaneous process (ΔG ≤0) and the SD genomic position according to experimentally observed trends in *Escherichia coli* (Saito et al., 2020). Promoter and intrinsic termination loops (**Figs**. S9-S13) fall within the known characteristics (genomic positions and sequence) observed in the literature (de Hoon et al., 2005; Lloréns-Rico et al., 2015; Matteau et al., 2020). For further details regarding promoter, Shine-Dalgarno, and intrinsic termination loops motif detection see **Section** S2. A complete record of the motif information is compiled in **SI File 3**.

**Table 1.** Motif Identification Summary: Data for Syn1.0, Syn3A, *M. mycoides* subsp. *capri* (Mmy), *M. florum* (Mfl), and *M. pneumoniae* (Mpn). Numbers for Shine-Dalgarno and promoters are for motifs predicted to be functionally active. Protocol to identify the features is explained in **Section 4.1.** Sequences describing the motifs are reported in **Table** S4. Transcription units are identified using a protocol outlined in **Section 4.2** and **Fig. 10**. Results (with red column headers) from previous studies are also presented. Qualitative agreement between organisms and methods is observed.

|  | Syn1.0<sup>1,*</sup> | Syn3A<sup>1</sup> | Mmy<sup>1,◇</sup> | Mfl<sup>1</sup> | Mfl<sup>2</sup> | Mpn<sup>1</sup> | Mpn<sup>3</sup> |
| --- | --- | --- | --- | --- | --- | --- | --- |
| Gene-ORF | 911 (42) | 496 (3) | 911 (40) | 724 (7) | 720 (-) | 774 (44) | 733 |
| Shine-Dalgarno | 539 | 308 | 542 | 421 | 548 | 223 | - |
| Promoter | 659 | 353 | 668 | 537 | 433 | 264 | 709 |
| Intrinsic Terminator | 371 | 185 | 372 | 296 | 298 | 159 | 225 |
| Bioinformatic TUs | 753 | 400 | 757 | 589 | 387 | 389 | 447 |
| Genes per TU | 2.5 | 2.9 | 2.5 | 2.4 | 2.2 | 4.5 | - |
| Kbp per TU | 2.63 | 2.67 | 2.64 | 2.36 | 2.40 | 4.30 | - |

<sup>1</sup> Data from this study
<sup>2</sup> Data from Matteau et al. \(2020\)
<sup>3</sup> Promoter data from Lloréns-Rico et al. \(2015\), Gene-ORF, Intrinsic terminator, and TUs data from Güell et al. \(2009\)
\* There is a discrepancy in the number of genes reported in the GenBank file (911) and literature (908) (Gibson et al., 2010). Mmy gene count should be larger than Syn1.0.
◇ There is a discrepancy in the number of genes reported in the GenBank file (911) and literature (917) (Gibson et al., 2010). Mmy gene count should be larger than Syn1.0.

A transcription unit (TU) is the region of the DNA transcribed during a single transcription cycle of initiation, elongation, and termination. To define the transcription unit, only the transcription start (initiation) sites (TSSs) and the stop (termination) sites (TTSs) are needed. The mechanism of promoter recognition and subsequent transcription initiation has been well-studied and structurally observed in Chen et al. (2020). Therefore, we assume initiation occurs at the promoter motifs and identify them as the TSSs. The TSSs could not be resolved with any certainty by our bio-informatic methods as the TSS is a single nucleotide (A/G), see **Fig. 1A**. Termination within Gram-positive bacteria is observed to commonly proceed through intrinsic termination loops, so then the TTSs are identifiable via the intrinsic termination loops. Possible TUs are predicted by considering all regions bordered by a matching pair of the previously identified start and stop sites. We assume that every promoter motif is acting as a site for transcription to start and every predicted terminator causes a stoppage in transcription to define TUs without evidence for the activity of the sites. A more complete explanation of the TU identification protocol is presented Materials and Methods–**Section 4.2** and **Fig. 10**. The results of the TU discovery are reported in this study are provided in **Table 1**.

Of the organisms studied, all but *M. pneumoniae* have generally close agreement among the metrics describing the TU results (**Table 1**). The average length in kbp and the average number of genes per TU are almost identical. The ratio of gene ORFs to transcription units (∼0.80) is consistent for all organisms except *M. pneumoniae*. The differences between *M. pneumoniae* and the other four organisms is not surprising given that *M. pneumoniae* is a member of a different phylogenetic branch of *Mycoplasmas* than the other species. Evidence of this change is that the genus name of *M. pneumoniae* has recently been changed to *Mycoplasmoides*. The similarity in results is anticipated as genome architecture is conserved between related organisms especially in the case of the *M. mycoides* subsp. *capri*, Syn1.0, and Syn3A due to genome minimization. The outlier of *M. pneumoniae* can be explained by the promoter and intrinsic termination loop results. For example, promoter results for *M. pneumoniae* (**Fig**. S13B), show more promoters being assigned non-functional than functional, a trend not seen in the organisms. The lower number of identified motifs means a lower number of of paired TSSs and TTSs, thereby leading to a lower number of predicted TUs. Our resulting TUs for *M. florum* and *M. pneumoniae* agree with TUs previously identified using other methods for *M. florum* (Matteau et al., 2020) and *M. pneumoniae* (Güell et al., 2009; Lloréns-Rico et al., 2015), respectively. Our results agree with these other predictions derived from different methods than those used here. The SD is not related to transcription and does not play a role in TU identification. However, a reasonable fraction (∼0.35) of the leading gene (first to be transcribed) within a TU are identified with a functional SD. A majority of the detected SD are within transcription units.

### 2.2. Transcriptional Activity Observed from RNA Sequencing

Short- and long-read RNA sequencing were performed on Syn1.0 to systematically investigate the transcriptional activity within the cell. **Fig. 2A** provides a simple outline of the technique, highlighting the four major steps: (i) RNA sample preparation, (ii) RNA sequencing via flowcell loading, (iii) base-calling, which is the signal conversion to nucleotide sequence, and (iv) mapping transcripts back to regions along a reference genome. The specifics of each step vary depending on the protocol used, and exact details have been provided in Materials and Methods. The resulting characteristics of the experimentally observed transcriptome are inherent to the methodology used, short-read vs long-read (see **Fig. 2B**) and stranded vs non-stranded. Long-read methods (ONT and PacBio) enabled the analysis of full length RNA transcripts at a single-molecule level, with ONT providing an additional advantage as the sequenced data is direct and stranded as compared to PacBio. Due to inherit bias in long-read techniques, a short-read approach (Illumina) was used to more accurate quantify the RNA within Syn1.0. We elected to use stranded Illumina RNAseq to have a more direct comparison with the ONT results. Initial results provided a broad characterization of the transcriptome (minus rRNA) within Syn1.0. **Fig. 2C** characterizes the experimental reads length distributions of each technique. Long-read methods showed significantly shifted distributions as compared to the short-read data with PacBio resulting in the longest observed transcripts. A summary of the RNAseq data is given in **Table** S5. PacBio and Illumina showed the highest percentage of mapped reads (∼99%) and genomic coverage, ∼790x and ∼125x respectively. While ONT was outperformed by the others, its mapping percentage (∼80%) and genomic coverage (∼60x) were significantly greater than those observed for a variety of organisms in (Grünberger et al., 2019a) (Grünberger *et al*. mapping percentage are approximately 56% and the average genomic coverage observed was 11.5x.). Our relatively high mapping percentages are indicative of high quality data. The reduction observed in the ONT data can potentially be explained by the lack of cDNA conversion in the *direct* method (Garalde et al., 2018). With no cDNA conversion, post-transcriptional processing of RNA such as RNA modifications is observable in direct RNAseq (Liu et al., 2019). A transcript altered via a post-transcriptional process may have a sequence when sequenced and base-called that appears as if it did not originate from the target reference genome resulting in a lower than expected mapping percentage. More importantly the coverage values are relatively high in all three methods. Coverage represents a measure of the average sequencing depth across the entire genome, therefore a greater value directly correlates to greater observations of transcriptional activity. Full descriptions of the Syn1.0 RNAseq coverage observed for each method and replicate are reported in **SI Files 6–8**.

**Figure 2.**
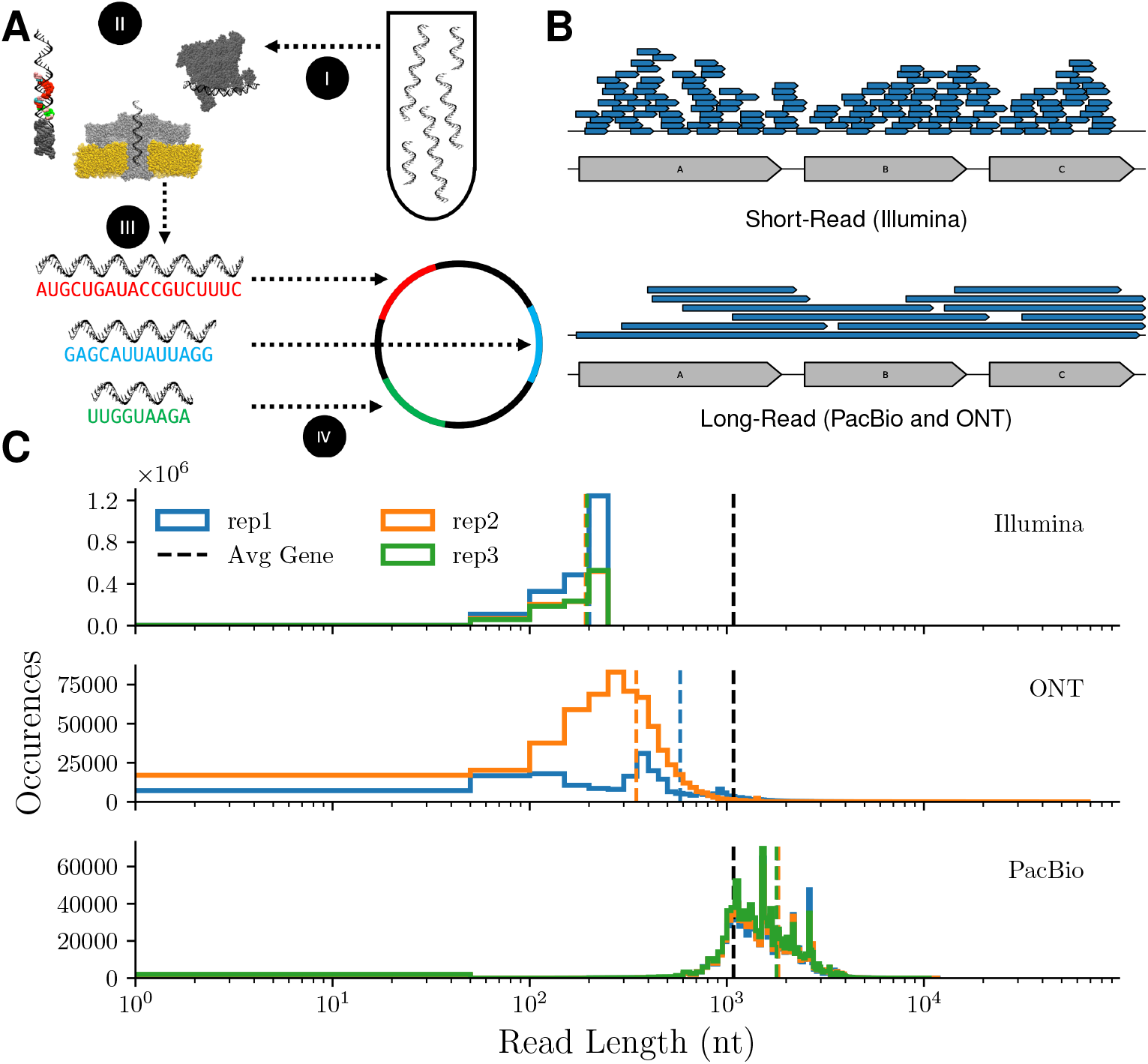
Summary of the RNA Sequencing: **(A)** Schematic of the RNA sequencing experimental methodology with the following steps: *(i)* RNA sample preparation, *(ii)* RNA sequencing via flowcell loading, *(iii)* base-calling, which is the signal conversion to nucleotide sequence, *(iv)* mapping transcripts back to regions along reference genome. **(B)** Diagram highlighting the difference between short-read (Illumina) and long-read (PacBio and ONT) RNA sequencing methodologies. Typically short-read methods provide significantly more reads, but fail to capture transcripts with length near average gene length. **(C)** Length distribution of Syn1.0 RNA transcripts sequenced for each RNAseq method. Average length of gene in Syn1.0 is shown with black dotted line.

Sequencing depth is a measure at nucleotide resolution of the frequency a transcript has been detected and mapped to a reference genome within an RNAseq experiment. Therefore it enables a semi-quantitative description of the transcriptional activity observed for a sample. Given that RNA is only formed via transcription any non-zero sequencing depth is indicative of some transcription event. It is important to note that sequencing depth will not disclose any post-transcription events simply relative characterization of transcription. However, transcriptional activity alone is quite informative in being able to understand expression patterns among a genomic region. For example, steady sequencing depth along a region indicates that expression is consistent within a region; whereas variable sequencing depth presents inconsistent expression. Furthermore, the change in sequencing depth with respect to the direction of transcription is a direct indication of a transcription initiation (increasing) and termination (decreasing) especially in long-read RNAseq (Tarnowski and Gorochowski, 2022). For example, single and clusters of genes within well-characterized operons show varied transcriptional patterns corroborating a revised view of operons reviewed in Taggart et al. (2021). Specific operon examples are shown in **Figs**. S17-S19.

Sequencing depth also enabled the discovery of new genetic features previously unidentified in Syn1.0. **Fig. 3** and **Fig**. S21 depict transcriptional activity measured in regions of the genome where there are no known motifs. These two examples represent two different type of newly observed genetic features. **Fig. 3** is an example of a completely new gene ORF (likely encoding a non-portein coding gene). Illumina and ONT RNAseq methods indicate significant expression within the forward strand region downstream of MMSYN1_0884. The results show sharp increases and decreases in the sequencing depth not seen within the neighboring genes indicating a unique transcription event (initiation and termination). PacBio corroborates the observation of an expressed gene, however due to a lack of strand specificity within the experiments it cannot be discerned the extent of expression originating from the forward strand specifically. This is confirmed by the observed sequencing depth across the reverse strand gene, *gidA*/0885. Additional evidence was confirmed by examining a ONT run from Syn3A where this region has been maintained during the genome reduction and shows a similar expression trend (see **Fig**. S20). Computational efforts to identify the gene downstream of MMSYN1_0884 were unsuccessful.

**Figure 3.**
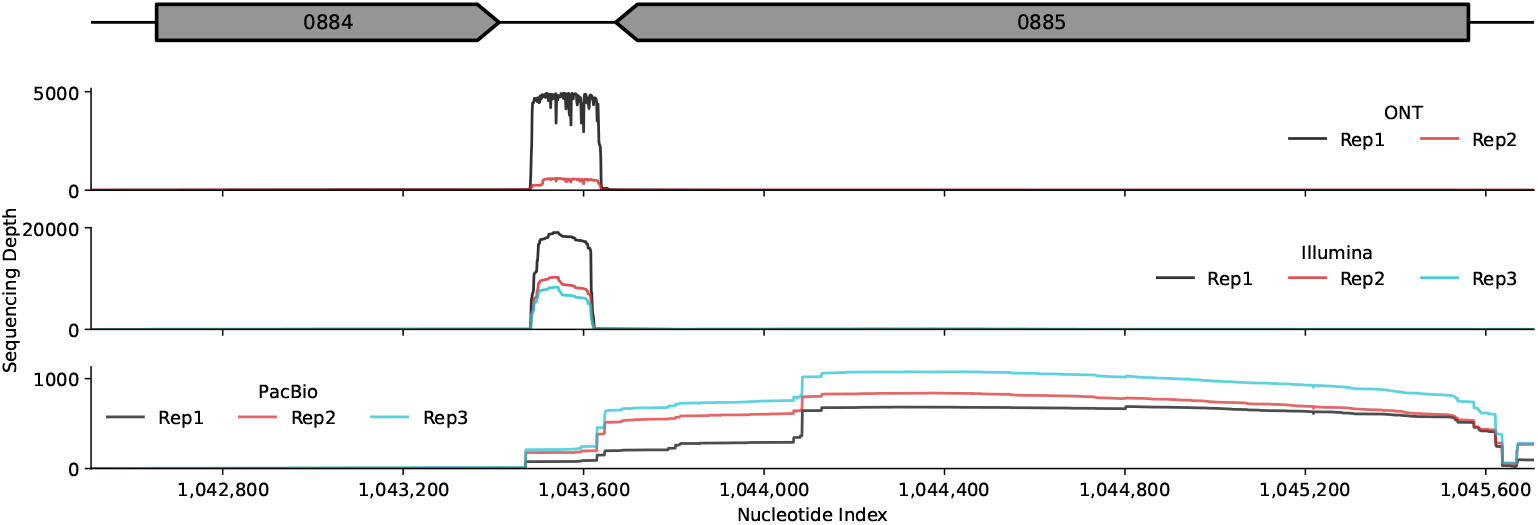
Intergenic Transcription Activity in JCVI-syn1.0: Sequencing depth measured in ONT (top), Illumina (middle), and PacBio (bottom) RNA sequencing experiments in Syn1.0. Illumina and ONT show significant sequencing depth (forward strand) within a region of the genome where no gene ORF has been identified. PacBio results show this as well but not as pronounced due to the lack of strand specificity within the PacBio methods (both strands shown in data). Transcriptional activity is observed by measuring a non-zero sequencing depth indicating a transcript was sequenced and mapped back to genomic region. Sequencing depth within MMSYN1_0884 is observed, however it appears to be undetected in the plot because the sequencing depth of the intergenic region is significantly greater. The intergenic expression is indicative of a previously unannotated gene, and is also observed to be transcriptionally active in Syn3A (see **Fig**. S20).

Using similar logic to the intergenic expression, possible anti-sense RNAs (asRNAs) were identified. asRNAs are regulatory motifs proposed to have regulatory functions either through RNA duplex formation or transcriptional interference (Lloréns-Rico et al., 2016). They are found on the opposite strand to a gene ORF, therefore significant anti-sense transcriptional activity suggests their presence. **Fig**. S21 presents a reverse strand gene, MMSYN1_0293, and its respective anti-sense (forward strand) transcriptional activity highlighting possible asRNA detection. Each of the RNAseq methods show corresponding anti-sense expression. In comparison the sense expression (see **Fig**. S22) shows a notable difference. (PacBio lacks of strand specificity results in reverse and forward data being nearly identical.) This asRNA was previously identified by Lloréns-Rico et al. (2016) with Illumina RNAseq in Syn1.0; however short-read methods cannot distinguish if the anti-sense expression is a result of transcriptional read-through. Due to the long-read data, we can confirm this is an asRNA generated from a distinct transcription event. However we cannot ascertain whether is has a distinct function or is a result of pervasive transcription, which is known to be more common in A-T rich genomes such as that of Syn1.0 (Lloréns-Rico et al., 2016). Complementary investigations outside the scope of this study into these genes (and similar genes) are highly important to better understand the transcriptional landscape as these genes belong to the sRNA family, which are well-characterized regulatory elements typically inhibiting gene expression via some post-transcriptional process (Storz et al., 2011; Balasubramanian and Vanderpool, 2013; Mars et al., 2016; Boutet et al., 2022).

To quantify the expression patterns, we examined the intracellular abundances determined by the RNAseq methods. Transcript per million (TPM) was used to normalize counts for each gene to compare replicates and sequencing methods. Additionally, absolute abundances of RNA species were calculated using the relative abundances (TPM) to scale the estimated number of each RNA species. For a single cell, RNA numbers were estimated using the molecular weights and the an assumed total RNA dry mass taken from related organisms. To account for specific RNA types, we enforced rRNA, tRNA, and mRNA totalize 80%, 15%, and 5% of the total RNA mass of a cell (Milo et al., 2009; Westermann et al., 2012; Matteau et al., 2020). Determination of TPM and absolute abundances values for genes is explained in **Section 4.7**. To assess the absolute copy numbers, the values were converted to RNA cellular concentrations compared to a related organism (see **Table** S7). Calculations were done assuming a Syn1.0 diameter of 439 nm reported in Moger-Reischer et al. (2023). The concentrations among the RNAseq methods agree quite well with only some slight difference among the rRNA for the short- and long-read methods. Averaging across all experimental data, RNA concentrations were calculated to be 6.6 × 10^3^, 4.0 × 10^3^, 4.8 × 10^4^, 1.7 × 10^5^ RNA/*µm*^3^ for mRNA, ncRNA (sRNA, tmRNA. etc), rRNA, and tRNA, respectively. Matteau *et al*. obtained very comparable data for *M. florum*’s RNA, 4.7 × 10^3^, 5.4 × 10^4^, and 2.0 × 10^5^ RNA/*µm*^3^ for mRNA, rRNA, and tRNA, respectively (Matteau et al., 2020), Table EV1. The PacBio abundances are likely to be inflated due to the lack of strandedness in the experiments, enabling transcripts assigned to their opposite strand. All abundances data has been provided in **SI File 5**.

Short-read (Illumina) RNAseq is traditionally used for RNA quantification so these values were assumed as the baseline to compare the other methods. Generally, the gene copy numbers obtained from the RNAseq methods are correlated assessed by Pearson’s correlation coefficient (linear) and Spearman’s (non-linear, monotonic) correlation coefficient (see **Fig**. S16). There is significant overlap among the relatively highly expressed genes detected across the three RNAseq methods. In the 95^th^ percentile of expressed genes, representing 45 genes, a total of 29 genes are consistent among the three methods. A significant majority of these highly expressed genes are tRNA genes (23), however of great interest are two genes, *ssrA*/0158 encoding transfer-messenger RNA (tmRNA), *rnpB*/0356 encoding baterial RNase P class B, and MMSYN1_0916 encoding an RNA interference gene. Each of these genes are implicated in processes responsible for regulating bacterial homeostasis: tmRNA – rescuing stalled ribosomes (Rae et al., 2019), baterial RNase P class B – processing of pre-cursor tRNA (Frank and Pace, 1998), and RNA interference – a mechanism used by the cell to regulate gene expression (Wilson and Doudna, 2013). Assuming the relatively high expression holds in Syn3A, a reduced-genome version of Syn1.0, the functions encoded by the retained genes, *ssrA*/0158 and *rnpB*/0356, are exceedingly critical to bacterial life beyond the essentiality results from Breuer et al. (2019). Deviations between Illumina and the other methods cannot be ignored, and are likely attributed to the read assignment protocol of HTseq, a high-throughput RNAseq python analysis module that allows for ambiguous assignment of transcripts (Anders et al., 2014). The long-read methods have roughly a factor of 10^2^ (ONT) and 10^3^ (PacBio) more ambiguous assignments than the Illumina data (Data not shown). This trend is reasonable as the long-read methods have transcripts spanning multiple genes, which results in ambiguous (multiple) gene assignment.

### 2.3. Long-Read RNA-Seq Exclusive Analyses: RNA Isoform Profiling and Transcription Unit Prediction

Within the long-read RNAseq data is a unique opportunity to examine the transcriptome in its more native state accounting for RNA isoforms, the set of mono- or polycistronic transcripts evolved from a region of the genome. The complexity within a set of isoforms is an effect of differential transcription due to the stochastic nature of gene expression (Taggart et al., 2021) and post-transcription processing such as RNA cleavage/degradation (Taggart et al., 2021, Taggart et al., 2025) or RNA modifications (Liu et al., 2019). Although this study’s results are more amenable to characterizing the impacts of differential transcription, the long-read RNAseq supplies a general view of the post-transcriptional processing of Syn1.0. Each RNAseq read (RNA isoform) assigned to a single or multiple gene ORF was examined for completeness with respect to genetic information of its assigned gene ORFs. An RNA isoform was considered full-length if its mapping coordinates on the reference genome completely overlapped with all its assigned genes start and end coordinates. Within all replicates from two different techniques, a significant majority of RNA isoforms are incomplete. The results are plotted in **Fig. 4**. Two possible reasons for this trend are the strict definition of a full-length transcript (complete overlap) and/or perturbations during sequencing causing errors in the read. However, a similar trend was observed in other bacteria *E. coli* (Herzel et al., 2022) and *Mycobacterium tuberculosis* (Ju et al., 2024). Coupled with the fact that Syn1.0 has genes capable of RNA processing, it is reasonable to claim these results (**Fig. 4**) are genuine evidence of frequent post-transcriptional RNA processing. In an attempt to further characterize the potential post-transcriptional processes of JCVI-syn1.0, we further analyzed the ONT direct RNAseq. Direct RNA sequencing gives the possibility of identifying methylated adenine bases (m^6^A) (Liu et al., 2019), because its sequencing template is the RNA isolated from bacterial cells rather than cDNA reverse transcripts. We adopted the strategy of (Liu et al., 2019, 2021), but no precises inferences can be made from the data (see Supplementary Information—**Section** S3).

**Figure 4.**
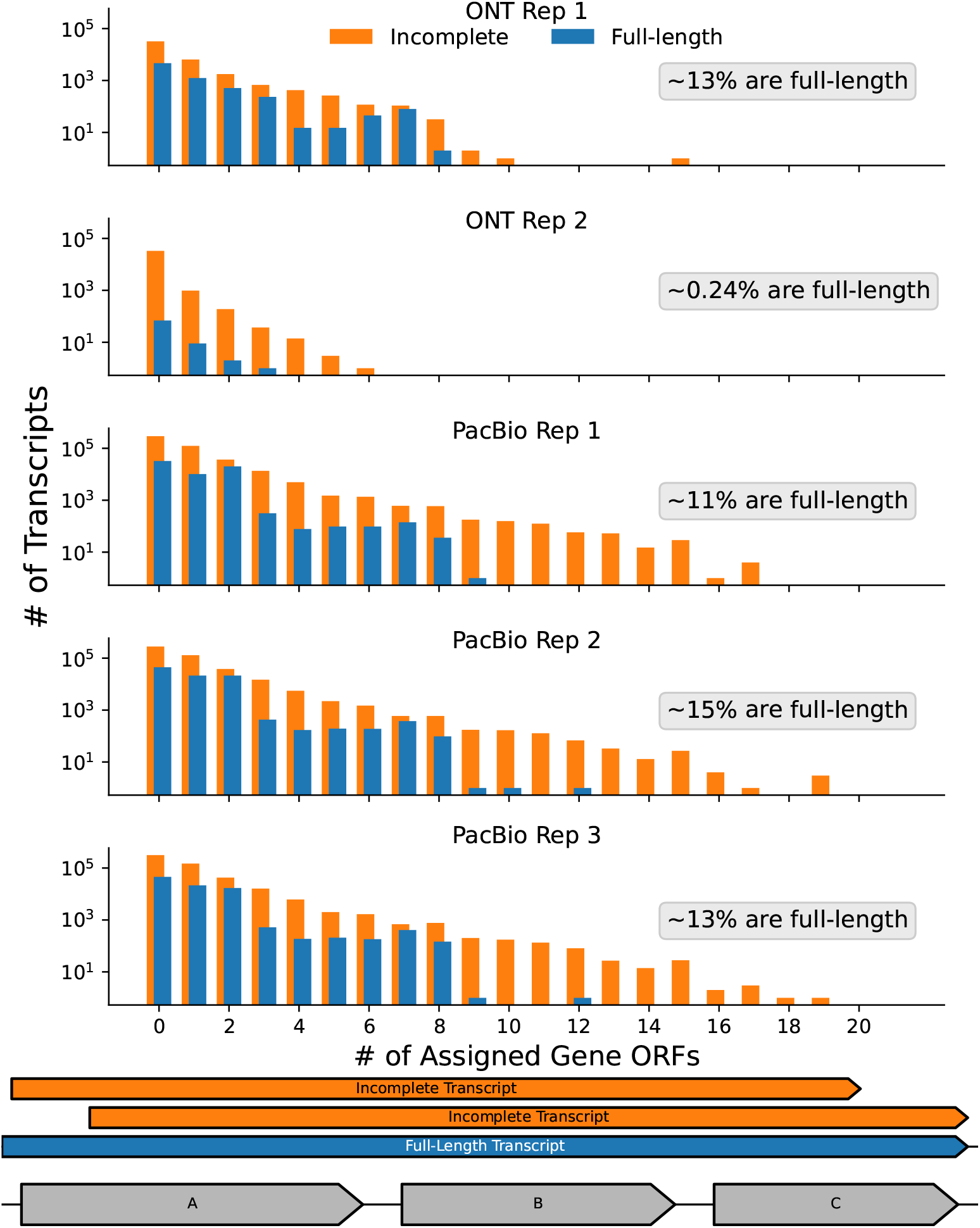
Transcript Completeness: Distribution of the transcript “completeness” for the long-read Syn1.0 RNAseq data for full and incomplete transcripts. A transcript is complete if all genetic information for all genes assigned to a read is found within the gene (blue). (Each assigned to a read is checked for overlap, see **Section 4.7.2** for read assignment protocol). Example transcripts for reads assigned to geneA, geneB, and geneC are shown at the bottom. Approximately 11% of transcripts are full-length/complete. Similar trends are observed in *E. coli* (Herzel et al., 2022) and *Mycobacterium tuberculosis* (Ju et al., 2024).

Given the high likelihood and magnitude of RNA processing, we needed to develop a way to discern if an RNA isoform was a result of transcription or additional processes. This was solved by using the *de novo* transcription unit predictions coupled with transcription units derived from the ONT RNAseq results. The TUs outline the initial formation of the RNA isoforms, and thus an RNA isoform that doesn’t have a corresponding transcription event must have evolved from additional processing. The RNAseq TUs are calculated only from the ONT data because it is strand specific, which is mandatory when assigning TSSs or TTSs to genes according to proximity. Within the *de novo* TUs, TSSs and TTSs are related to the identified promoters and termination motifs (see **Section 4.2**). For the RNAseq, the explicit TSS and TTS can be extrapolated from either the sequencing depth (Grünberger et al., 2019a; Tarnowski and Gorochowski, 2022) or the start and stop positions of the individual reads (Matteau et al., 2020). In this study, the explicit reads were used to hopefully provide more context of the RNA isoform characteristics. A read start per million of mapped reads (RSPM) and read ends per million of mapped reads (REPM) value was calculated at every nucleotide position for each strand. To determine significance of the results, only RSPM and REPM values greater than the sum of their average and standard deviation were accepted and nucleotide positions below the threshold were set to zero. Roughly <0.1% of all genomic positions had non-zero RSPM/REPM values indicating those genomic positions were transcription start/termination sites. This strategy was adapted from (Matteau et al., 2020). RNAseq predicted TSS and TTS were assigned to a specific gene based on proximity, either upstream (TSS) or downstream (TTS). In attempt to probe RNA cleavage, which would alter an RNA isoform, we compared TSS and TTS pairs between adjacent genes on the same strand from the two methods. Using the assumption that a TSS and TTS pair observed in the RNAseq data, but not among the *de novo* motifs suggests cleavage. Of 158 possible sites, 9 (∼ 5%) and 44 (∼ 28%) in the two ONT replicates were found to show active transcription termination and initiation without any predicted motifs. Further investigation into the identified sites did not highlight any consensus sequence, which was likely a result of a weak consensus for RNase Y targets recently found for *B. subtilis* (Taggart et al., 2025). A full comparison of *de novo* and RNAseq TSS and TTS is presented in **Fig**. S23 and shows good agreement between the different sets of TSS and TTS. TSS deviate slightly, which is likely due to the *de novo* over predicting promoters that may not be functional given experimental conditions. Deviation in the TTS values could be a result of a limitation in the *de novo* search, which did not search within genes. Final TUs predictions were generated using the scheme of **Fig. 10**. All predicted TUs are provided in **SI File 4**.

Combining the RNA isoforms with the TUs enables a relatively complete investigation into genomic regions. **Fig. 5** and **Fig. 6** showcase two unique regions of Syn1.0 and the ability of long-read RNAseq to characterize the impacts of differential transcription on the RNA isoform landscape. The first region, the Division and Cell Wall (DCW) operon, was previously discussed in **Section 2.2** with respect to inconsistent transcriptional activity over the region, in conflict with the ideal operon representation. The region appears to have two transcriptionally linked sub-regions: the first three genes (0520-0522) and the last three genes (0523-0525). The RNA isoforms depicted in **Fig. 5** (and **Fig**. S24) strongly reinforce this notion. Simultaneously the TUs confirm this as well (**SI File 4**), while also indicating the other isoforms are likely a result of transcription than cleavage of RNA isoforms spanning the entire sub-regions.This expression difference is relevant for the cell as *ftsZ* /0522 and *ftsA*/0523 need to be in at least a 5:1 ratio in *B. subtilis* and *E. coli* for proper function (Feucht et al., 2001; Rueda et al., 2003). Previous work by Benders *et al*., corroborate this behavior in other *Mycoplasmas*, where internal transcription initiation within the DCW operon was observed (Benders et al., 2005). The *de novo* motif identification and RNAseq, which define the TUs, confirm this observation with the predictions of transcription initiation sites (promoters) upstream of *yqkD*/0520 and *ftsZ* /0522 (**Fig**. S25). The low expression observed within the second sub-region is a result of a transcription factor, MraZ, binding site known to repress transcription in *Mollicutes* (Fisunov et al., 2016). This expression difference is relevant for the cell as *ftsZ* /0522 and *ftsA*/0523 need to be in atleast a 5:1 ratio in *B. subtilis* and *E. coli* for proper function (Feucht et al., 2001; Rueda et al., 2003). We hypothesis the RNA isoform landscape enables this proper stoichiometry to primarily be achieved at the transcriptional level via an increased probability of transcription initiation events in the first sub-region coupled with regulation of expression in the second sub-region; however, a more targeted study would be needed to confirm this hypothesis. **Fig**. S26

**Figure 5.**
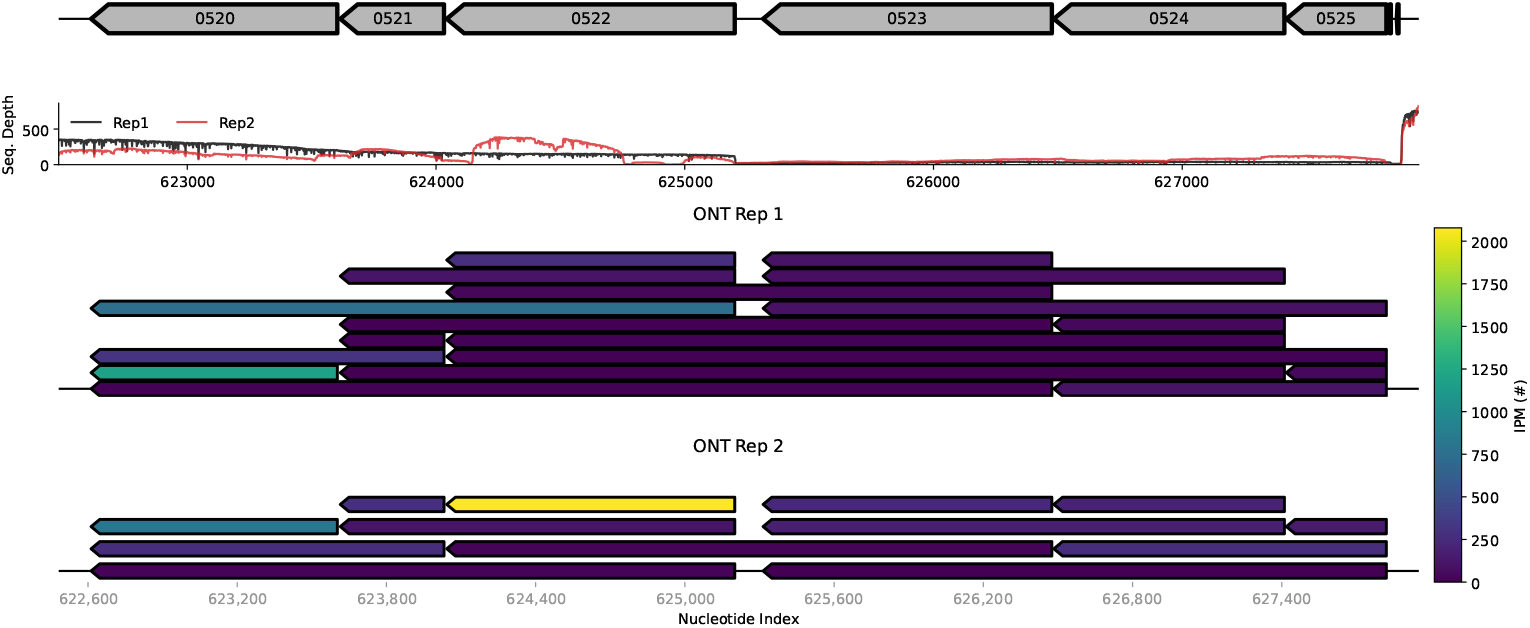
DCW “Operon” RNA Isoforms: (top) ONT RNAseq derived isoforms from the Division and Cell Wall (DCW) genomic region. (mid) Traces of the ONT sequencing depth across the region. Large changes in sequencing depth reflect differences in transcription activity between the two sub-regions: (i) genes *yqkD*/0520, *sepF* /0521, and *ftsZ* /0522 and (ii) genes *ftsA*/0523, *mraW* /0524, and *mraZ* /0525. The first sub-region show relatively higher transcriptional activity. (bot) RNA isoforms are colored by absolute abundances according to observations in the ONT RNAseq data. RNA isoforms are drawn with lengths overlapping their associated genes.

**Figure 6.**
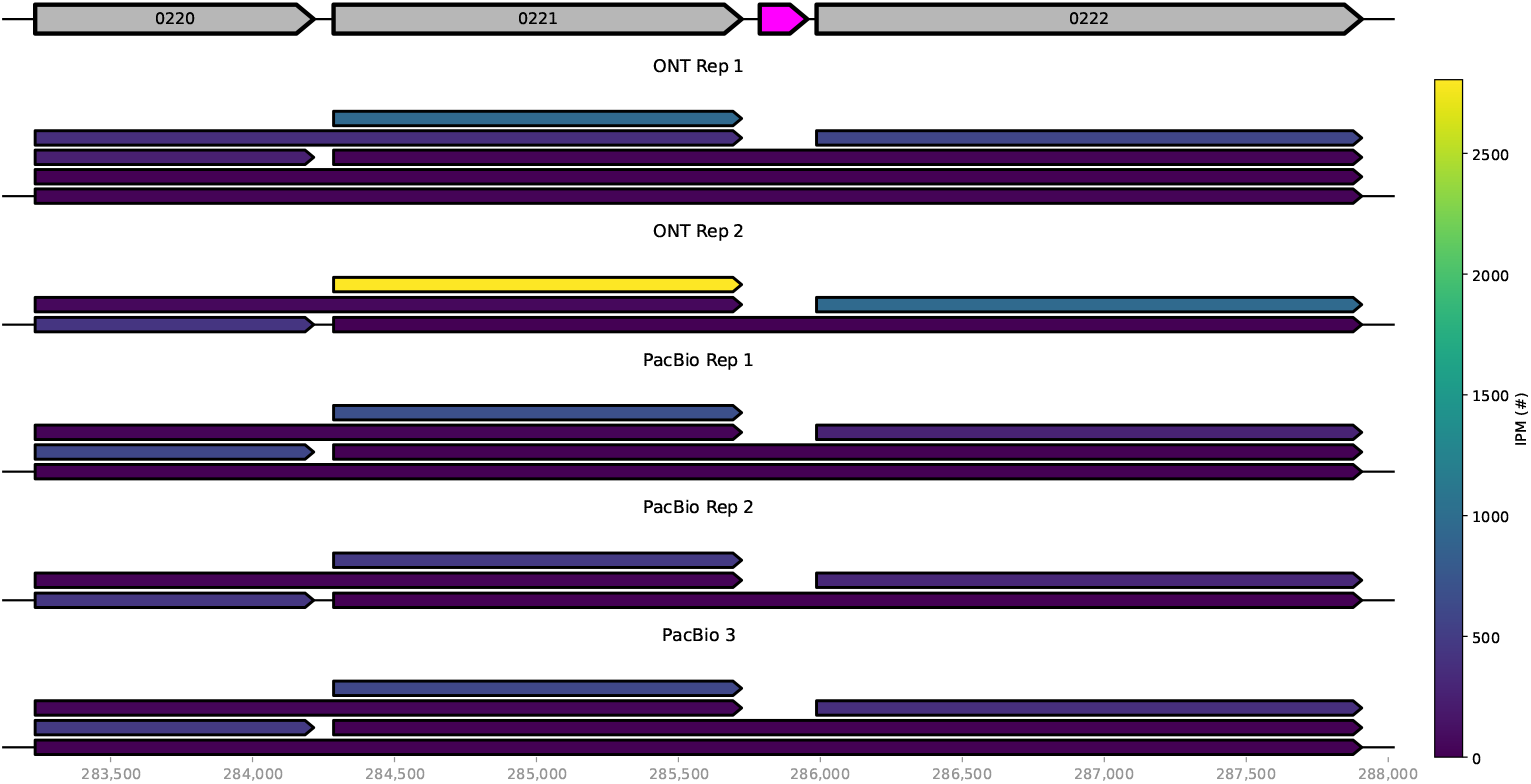
T Box Riboswitch I Region RNA Isoforms: (top) Region of Syn1.0 where the T Box Riboswitch I was identified (pink) using the RiboDB scanner (Mukherjee et al., 2019). The riboswitch is observed upstream of *thrRS*/0222, the threonine tRNA ligase. (bot) RNA isoforms are colored by absolute abundances according to observations in the RNAseq data. RNA isoforms are drawn with lengths overlapping their associated genes. Data shows instances of full transcription of *pyk* /0221 and *thrRS*/0222 (active riboswitch state, no transcription termination), however a majority of the isoforms are monocistronic suggesting an inactive riboswitch state (transcription termination). The monocistronic isoforms likely indicate unique transcription events due to their relative difference in abundances this is corroborated in **Fig**. S27.

The DCW undergoes inhibition of transcription while the second region, shown in **Fig. 6**, engages in a mechanism that inhibits transcription termination via a riboswitch. Riboswitches are non-coding RNA, which bind small molecules, to modulate gene expression typically via conformational changes in the riboswitch as a result of metabolite or ion binding within the aptamer domain (Mandal and Breaker, 2004). Two were previously identified in Syn3A in Breuer et al. (2019), and due to the relationship between Syn3A and Syn1.0, they are expected in Syn1.0. This study confirms their presence in Syn1.0 with sequence alignment. (Any) Additionally, two more riboswitches were found in both Syn1.0 and Syn3A using RiboDB Scanner (http://service.iiserkol.ac.in/~riboscan/) (Mukherjee et al., 2019). They are both T-Box riboswitches, which act via an anti-termination mechanism (Kreuzer and Henkin, 2018). **Figs**. 6-S27 showcase a region of one of the T-Box riboswitches found upstream of *thrRS*/0222. These riboswitches sense high uncharged tRNA abundances and prevent the intrinsic termination loop from forming, enabling transcription of the nearby amino acid tRNA ligase (Mandal and Breaker, 2004; Kreuzer and Henkin, 2018). Inactive T-Box riboswitches resulting from low uncharged tRNA abundances, do not block the intrinsic termination loop from forming, thus down-regulating expression of its respective ligase (Mandal and Breaker, 2004; Kreuzer and Henkin, 2018). Investigation into the RNA isoforms for the T-Box 1 genomic region suggest the regular inactivation of the T-Box 1 riboswitch (active intrinsic termination loop) because a majority of the isoforms do not encompass *thrRS*/0222 and the immediate upstream gene, *pyk* /0221 (see **Fig. 6**). The TSS and TTS (**Fig**. S27) and transcription units (**SI File 4**) are consistent with an inactive state of the riboswitch given the distinct TSS for *thrRS*/0222 and TTS for *pyk* /0221. It is critical to note the RNA isoforms do show full expression among all genes meaning that there must be some instances of an active riboswitch (inactive intrinsic termination). More targeted experiments are need to fully understand the extent the riboswitch switches between states.

### 2.4. Synthetic Evolution Reshapes the Proteome via Altering the Transcriptomic Landscape

The previous section focused on insights of the transcriptome landscape according to observations of the RNA isoforms and the transcription units generated from the bioinformatics and RNAseq data, primarily focusing on a single organism. However, the synthetic origin of the minimized bacteria, JCVI-syn1.0 and JCVI-syn3A, provides a unique platform to investigate the impact genetic additions or deletions have directly on the transcriptomic and thus proteomic landscape. To probe the changes between the two organism, we collected proteomic data from both Syn1.0 and Syn3Ain addition to the Syn1.0 RNAseq data. We leveraged recently collected short-read RNA sequencing data obtained from Sandberg et al. (2023), and old proteomic data sets, Syn1.0 (Glass, 2024) and Syn3A (Breuer et al., 2019). Our new proteomic removes bias observed as well as improved upon the detection of the older data sets. The primary purpose of the transcriptome is to generate the proteome, a finely tuned and robust process, which consistently generates proteins at near-ideal stoichiometric ratios (Taggart et al., 2020). We first sought to examine the relationship between the transcriptome in the within each organism individually. **Fig. 7** shows the abundances of the transcriptome vs those of the proteome. For comparison we have shown a similar plot for related organisms, *E. coli* and *M. florum* **Fig**. S26. According to these plots none of the organism have a strong relationship between RNA count and protein count. Excluding the Syn3A protein vs transcriptome data from this study, the synthetic organism shows a weaker relationship (determined by R^2^ values) than the natural bacteria. This could be due to less regulatory elements being present in the synthetic organisms, however the difference between the synthetic and natural organisms is quite small. Of more significance is the comparison between the transcriptome and proteome of Syn1.0 and Syn3A. Various proteins are observed to have distinct changes in their relative expression within the two minimized bacteria (see **SI File 1**). We postulate that some of these difference are a direct result of the genome minimization leading to perturbations in the genome sequence architecture and thus the produced transcriptome landscape.

**Figure 7.**
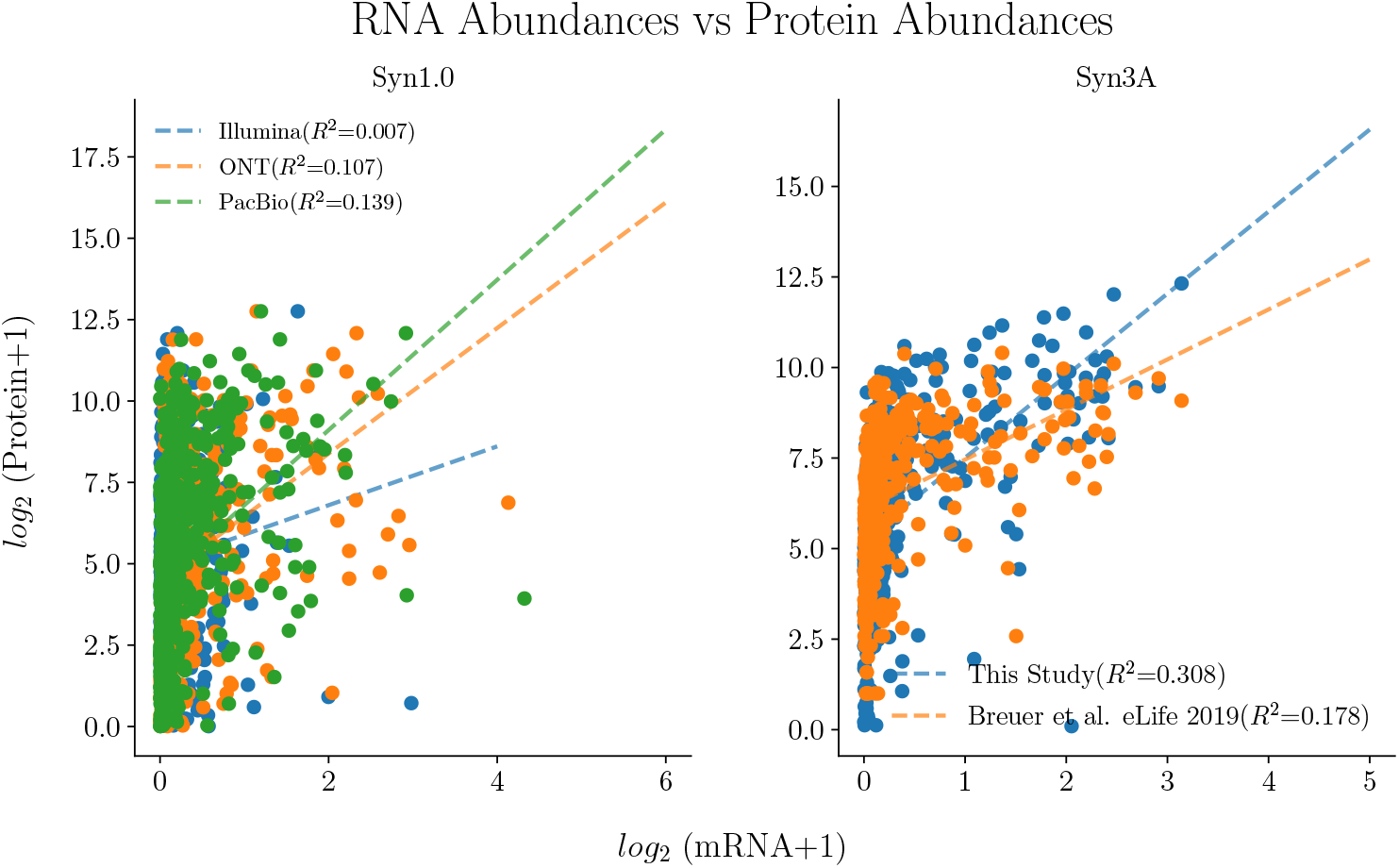
mRNA vs Protein Abundances: Relationship between standard-scaled abundances measurements of mRNA and its encoded protein for JCVI-syn1.0 (top) and JCVI-syn3A (bot). Dashed lines reflect a linear fit to the data. Coefficient of determination, R^2^, are provided of each fit. mRNA and protein abundances for Syn1.0 are measured in this study. Additional data is shown from a previously collected dataset (Glass, 2024). Protein abundances (all methods shown) while. Additional Mid data is RNAseq measurements from the stationary(left) and exponential (right) growth phases of Syn3A collected by Sandberg et al. (2023). Syn3A protein measurements were acquired from Breuer et al. (2019) and the data reported herein.

**Fig. 8** attempts to provide context on the impacts of altering the genome sequence architecture and the resulting change imposed on the RNA isoform landscape. In the synthetic minimization of Syn1.0 to Syn3A, genes were removed without accounting for the surrounding genome sequence architecture (Hutchison et al., 2016; Breuer et al., 2019). The removal of one gene was observed to have unintended consequences on another neighboring gene. *gpsA*/0349 was determined to be non-essential in Syn1.0 (Gibson et al., 2010), and thus was removed in Syn3A (Breuer et al., 2019). While Syn3A cells are able propagate and survive, chromosome conformation capture maps lacked chromosome interaction domains delineated by convergent/divergent gene boundaries, which are indicative of persistent super-coiling resulting from transcriptional activity in other bacteria (Le et al., 2013; Trussart et al., 2017). This atypical result in Syn3A is proposed to be caused by relative low expression of the protein encoded by *hupA*/0350 (Gilbert et al., 2021). In contrast, Syn1.0 does not have this relatively low expression of *hupA*/0350 protein, the protein actually corresponds to one of the highly expressed proteins within the cell (see **SI File 1**). The absolute abundances span two orders of magnitude within the two cells, ∼30 in Syn3A (24^th^ percentile in expression), and ∼3000 in Syn1.0 (99^th^ percentile in expression). **Fig. 8A-B** shows the genome sequence architecture and sequencing depth changes between the two organisms. The relative transcriptional activity for *hupA*/0350 in Syn1.0 is significantly higher than the neighboring genes while in Syn3A the expression is relatively consistent across all genes in the region. The Syn1.0 RNA isoforms, **Fig. 8C** and **Fig**. S28, confirm the co-expression because a majority of *hupA*/0350 encoding transcripts also encode for *gpsA*/0349. The TSS upstream of *hupA*/0350 is contained within the 3^*′*^ end of *gpsA*/0349 (see **Figs**. S29-S30), so the removal of *gpsA*/0349 led to disruption in the co-expression of the two genes. Another example of a similar reduction of protein expression from Syn1.0 to Syn3A is *dut* /0447, which had a five-fold decrease (139 to 24, see **SI File 1**). We attribute this decrease to a the removal of MMSYN1_0446 upstream of *dut* /0447. We compiled a list of all genes with shifts greater than 50% in their percentile, and hypothesized if the change in genome sequence architecture resulting from genome reduction is the primary contributor to the change in expression (see **SI File 1**). However, to conclude whether or not the perturbation of the genome sequence architecture is the main contributing factor for the decreased expression between Syn1.0 to Syn3A, the native genomic region must be restored followed with proteomic analysis. These observation motivates additional care during genome minimization to ensure unintended phenotypic variations do not arise. Other work has come to a similar conclusion when approaching genome minimization, concluding it could potentially allow for even more minimized genomes (Lachance et al., 2021).

**Figure 8.**
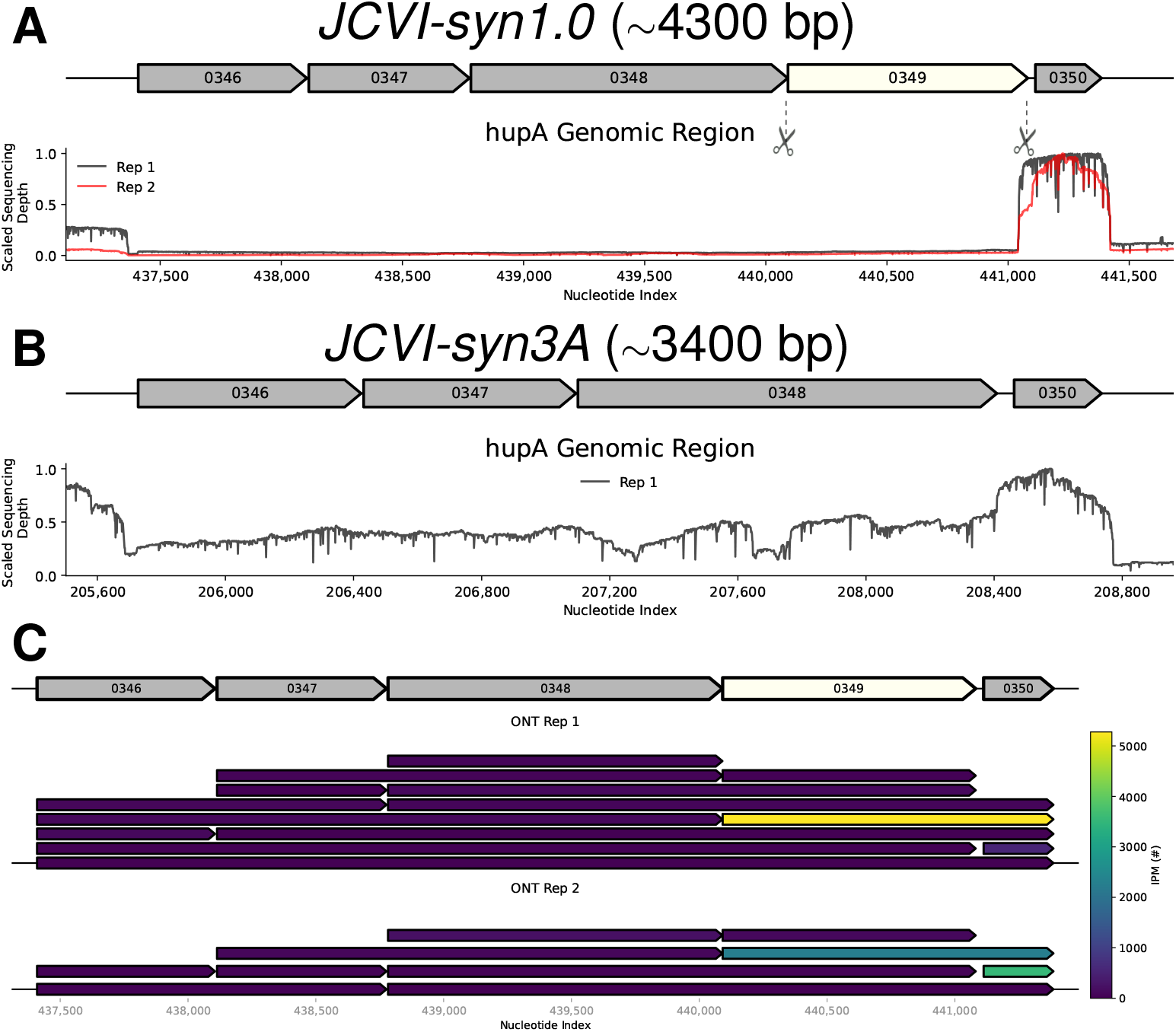
Impact of Genome Reduction: Comparison of the genome architecture between Syn1.0 **(A)** and the genome reduced Syn3A **(B)**. The Syn1.0 region corresponds to ∼4300 nucleotides, while the Syn3A region is approximately 3400 nucleotides. *gpsA*/0349 and some downstream regions were removed during genome minimization (white colored ORF). Scaled sequencing depth traces plotted show the relative transcriptional activity within the regions in the two organisms. Colors of traces correspond to replicate index with no association between Syn1.0 and Syn3A. **(C)** ONT RNA isoforms from Syn1.0 are colored by absolute abundances according to observations in the ONT RNAseq data. (See also **Fig**. S28.) RNA isoforms are drawn with lengths overlapping the genes they are associated with. *hupA*/0350 and *gpsA*/0349 are commonly found encoded within a single RNA isoform, thus the removal of *gpsA*/0349 has resulted in significant (∼10^2^) drop in the protein expression of *gpsA*/0350 from Syn1.0 to Syn3A.

## 3. Conclusion

Gene expression complexity within bacteria produces distinct physiological states and the ability to respond to stimuli, and so understanding the emergence of this complexity is an important goal of biological investigations. We took a multi-faceted approach using bio-informatic analysis coupled with multiple RNA sequencing methods to uncover the factors shaping the transcriptome within a minimized bacterium. Beginning with the initial step in gene expression, we made a simple assumption that the genome sequence architecture outlines all transcription events and then used the literature known sequence motifs to identify them within bacterial genomes. Our bio-informatic approach was validated by the reasonable agreement with results from other studies (**Table 1**). We further validated our bioinformatics method using short- and long-read RNA sequencing methods, which highlighted the similarities via experimental and theoretical results for the transcription units/RNA isoforms (**Fig**. S23, **SI File 4**). However due the simple assumption of the bio-informatic approach, it fails to account for the dynamic activity of the genetic motifs and thus is limited in capturing the true landscape of the transcriptome.

We are able to overcome this limitation because of the RNAseq experiments, which provide a more thorough view of the transcriptome. This is evident in the our observation of transcriptional activity in regions completely un-identified by the bio-informatic method (**Fig. 3**). This result emphasizes the additional details RNAseq provides and highlights the importance for validation of computation with experiment. Additionally, our choice to use short- and long-read methods enables us to directly highlight the difference in information obtainable from each method. Short-read methods are most reliable for quantification (Stark et al., 2019), however to understand any aspect of co-expression long-read RNAseq methods are necessary. The long-read RNAseq results provide co-expression events via predicted transcription units/RNA isforms (**Fig. 5, Fig**. S25, **Fig. 6**) spotlighting the strength to characterize a more complete perspective of the bacterial transcriptome.

Our protocol is applicable to other organisms, however, JCVI-syn1.0 was an ideal case because of the reduced genome size, lack of complex processes such as transcription-translation coupling or Rho-dependent transcription termination, its synthetic nature. The genome reduction of Syn1.0 to Syn3A provided a unique opportunity to investigate how modifications to the genome sequence architecture could manifest within an organism. Using comparative proteomics (**SI File 1**) from Syn1.0 and Syn3Awe were able to identify a set of genes with large proteomic shifts, and then determine with the long-read RNAseq if those genes shared RNA isoforms with neighboring genes that had been removed during the synthetic genome reduction. We were then able to explain a relatively drastic change in physiological state observed in Gilbert et al. (2021) as an un-intended consequence of the genome reduction (synthetic evolution) altering of the transcriptome (**Fig. 8**).

### 3.1. Limitation of the Study

This study provides a foundation to better understand the events shaping the bacterial transcriptome, but further investigation is need to comprehensively characterize and eventually predict a cell’s transcriptome. For example, this work attempted to describe the post-transcriptional processes of Syn1.0 such as RNA modifications (**Section** S3) and differential translation (**Fig**. S31), however no conclusive statements can be inferred. Accurate predictions of RNA modifications with ONT requires more targeted rather costly experiments requiring samples with and without modification (methyltransferase) activity such as those performed in Tavakoli et al. (2023). Differential translation contributes to the transcriptome landscape as messengers are prevented from degradation by ribosomes during translation (Taggart et al., 2021). The potential importance of differential translation can be seen in some of the Shine-Dalgarno motif results (**Fig**. S31) and RNAseq data within operons, unfortunately however the Syn1.0 data is inconclusive. While limited in this study, a more detailed investigation could elucidate any relationship with Shine-Dalgarno sequence influenced differential translation and polycistronic messengers.Additionally, environmental factors were not taken into account in this study and can play in the gene expression pattern in a bacterial cell. Nevertheless we provide an outline for the characterization of the transcriptional landscape using experiment and computation. Our results motivate a revising the synthetic genome reduction strategy to preserve TUs. We hypothesize that the re-introduction of the native genome sequence architecture around *hupA*/0350 will restore the Syn1.0 in Syn3A. Additionally, our compiled results profile the transcriptional activity within Syn1.0, and provide some insight to Syn3A’s transcriptional behavior. A complete description of all the possible outcomes is needed to fully describe how bacteria tune their transcriptome. **Fig. 9** conceptualizes the processes modifying the transcriptome as it was examined in this study. The full potential of the results in this study is to improve the description of the processes of kinetic models used in whole-cell simulations such as Thornburg et al. (2019); Macklin et al. (2020); Thornburg et al. (2022). Differential transcription via the handling of operonal expression has already been shown to be advantageous to improve the capabilities of cell simulations (Sun et al., 2024). Incorporation of the data generated in this study and others like it with cell simulations will allow for more complex cellular phenomenon such as the stoichiometric protein expression (Zhao et al., 2019; Taggart et al., 2021) to be understood at fundamental levels.

**Figure 9.**
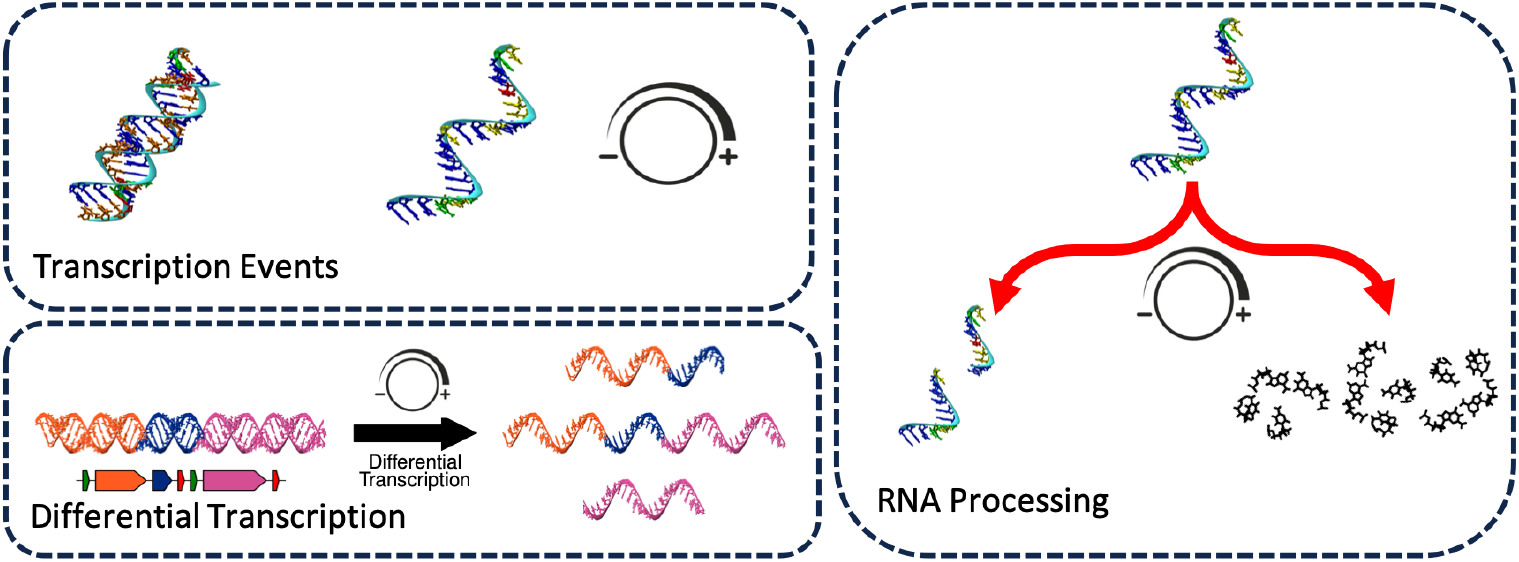
Categorization of Factors Impacting Transcriptome: A simplistic view of the categories of processes which lead to the formation of the transcriptome observed in bacteria. Transcription events include those that form RNA transcripts such as the newly identified expression of an intergenic region shown in **Fig. 3**. Differential transcription specifically accounts for the possible alteration of transcription events varying the encoded genetic information within RNA transcripts. RNA processing primarily accounts for the decay of RNA transcripts, but also includes post-transcriptional modifications to RNA transcripts, which would impact the encoded genetic information. Each category acts like a knob to “tune” the landscape of the shaping of the transcriptome. Complete depictions of each category (attempted with data presented in this study) will improve kinetic models of the genetic information processing such as those presented in Thornburg et al. (2019, Thornburg et al. 2022).

**Figure 10.**
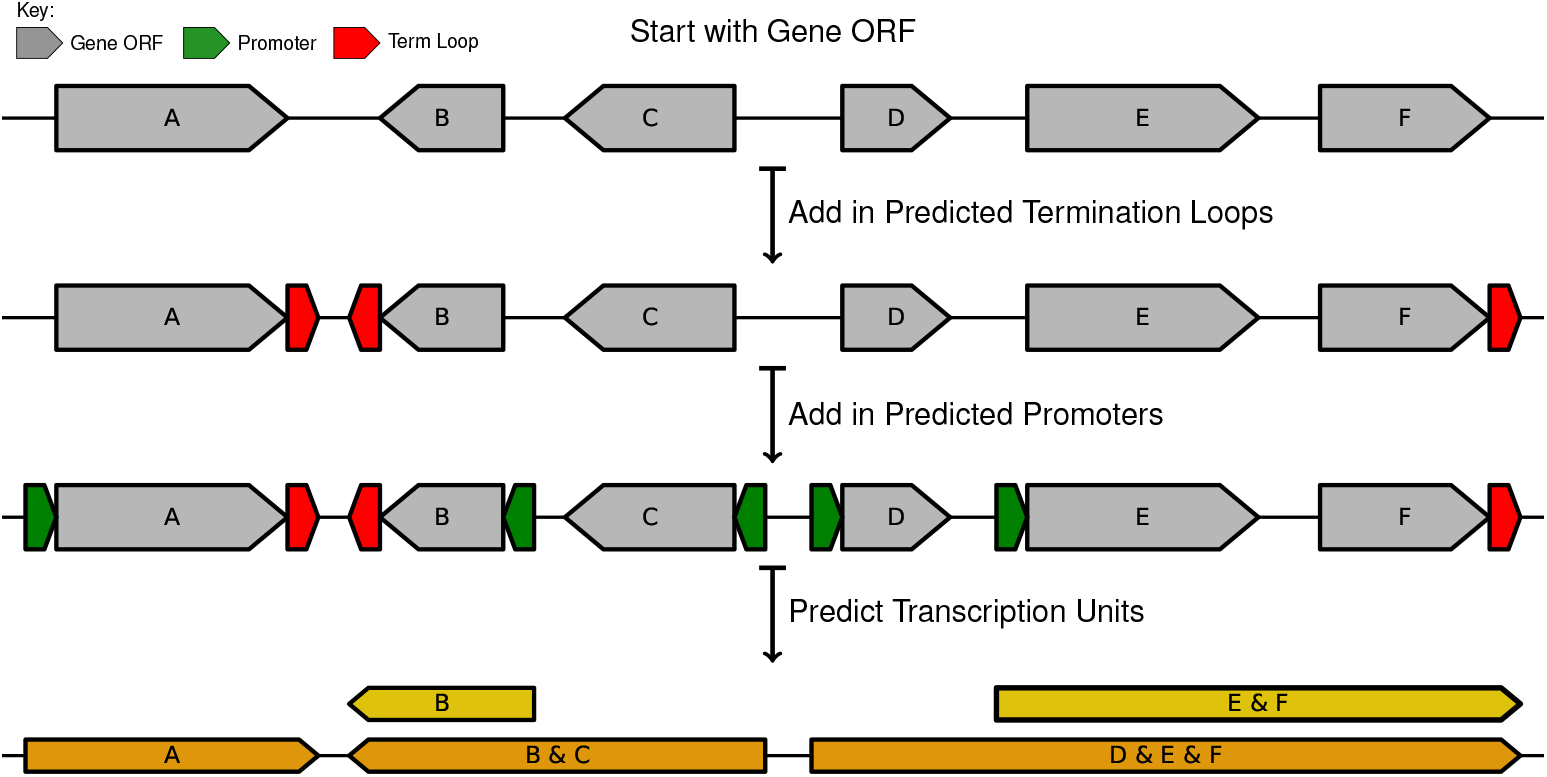
Transcription Unit Prediction Scheme: First, starting with the gene ORFs the predicted promoters (assumed to be the TSSs) are added, then the predicted intrinsic termination loops. (See **Section 4.1** for details about the motifs were predicted.) Finally, assuming the promoters act as the start site of transcription, and the intrinsic termination loops are the end sites of transcription, transcription units are predicted by from each promoter to each intrinsic termination loop. Transcription units can contain a single or many genes, and all genes must be on the same strand.

## 4. Materials and Methods

### 4.1. Motif Identification

#### 4.1.1. Gene Open Reading Frame

The genes open reading frame (ORFs), defined here as the DNA encoding a specific gene, were obtained for each organism in this study from the publicly available NCBI GenBank (see **Table** S3). Additional processing was needed for the GenBank file for Syn3A. The 23S and 16S rRNA within the GenBank are incorrectly designated resulting in incorrect RNA sequences. The 23S rRNA had an additional 13 nucleotides at the 5^*′*^ end, and was missing 6 nucleotides at the 3^*′*^ end. Similarly, the 16S rRNA had an extra 6 nucleotides and was missing 5 nucleotides at the 5^*′*^ and 3^*′*^ ends, respectively. The genomic coordinates were corrected for each rRNA, and the proper sequences are reported in **Fig**. S2.

#### 4.1.2. Shine-Dalgarno

To identify the Shine-Dalgarno within the bacterial genomes of interest, a bioinformatics based method was employed using an organism dependent Shine-Dalgarno template sequence to search for. We derived the template SD sequences using previously reported characteristics of the SD-aSD interaction. The Shine-Dalgarno, the 5^*′*^ end of a mRNA, interacts via RNA hybridization with the anti-Shine-Dalgarno, the 3^*′*^ end of the 16S rRNA (Shine and Dalgarno, 1974). The SD-aSD interaction has been observed structurally in *Thermus thermophilus* (PDB-ID: 2E5L) where eight nucleotides were observed to be involved in the hybridization between the two RNA strands (Kaminishi et al., 2007). In *E. coli* using orthogonal ribosomes, SD are reported to hybridizes at as many as nine nucleotide (Saito et al., 2020). The sequence of the SD has been characterized in many other organisms by the consensus sequence of 5^*′*^-*GGAGG*-3^*′*^ (Hockenberry et al., 2017, 2018; Matteau et al., 2020; Saito et al., 2020). We used the following criteria to define our template sequence: the template must contain the consensus motif (5^*′*^-*GGAGG*-3^*′*^) and be eight or more nucleotide in length. Additionally, we used each organism’s respective 16S rRNA 3^*′*^ end to define the ideal Shine-Dalgarno sequence to account for any variation between the 16S rRNA 3^*′*^ ends. For example, the Syn1.0 and Syn3A 16S rRNA nucleotide sequences ends with 5^*′*^-*CCTCCTTTCT* −3^*′*^, so therefore the template Shine-Dalgarno sequence is its reverse complement, 5^*′*^-*AGAAAGGAGG*-3^*′*^. We included total of 10 nucleotides to ensure the well characterized 5^*′*^-*GGAGG*-3^*′*^ consensus motif was present. All anti-Shine-Dalgarnosequences and corresponding Shine-Dalgarno templates sequences used in this study are reported in **Table** S4.

The relative position of a Shine-Dalgarno to its respective gene is roughly 5-15 nucleotides upstream from the gene ORF, which has been observed in related organisms. The template sequence and the approximate location of the Shine-Dalgarno are all that is needed for the identification algorithm. The algorithm has two steps: (1) detection of potential motifs and (2) assignment of functionality to the motifs. For each gene ORF, a region 25 nucleotides upstream of the gene ORF is selected for local sequence alignment against the template sequence. The local sequence alignment outputs the best possible match to the template indicated by the alignment score. In cases where multiple regions present the same score, the potential Shine-Dalgarno which minimizes the distance of the spacer region between the Shine-Dalgarno and gene ORF. The alignment score is an output of how well the template sequence is matched according to a substitution matrix, and thus is directly related to the affinity between the SD-aSD hybridization. The matrix used for the Shine-Dalgarno assumes a value for a nucleotide match as 1 and mismatch as −3. This match/mismatch ratio was used for scoring because it is for alignments where sequences share 99% identity and is the default matrix used in the NCBI BLAST implementation (Pearson, 2013). The alphabet contains thymine (T) because the DNA sequence is scanned rather than the RNA sequence. A gap and gap extensions in the alignment are penalized by the maximum score of the respective alignment to ensure no gaps are present in the local alignment. The search was performed via a custom Python script available at https://github.com/troyb2/MinimalCell_Motif-Identification_RNAseq, which used Biopython v1.78 (Cock et al., 2009) to parse and perform sequence alignments. Matrix for scoring was defined using a custom matrix building function of Biopython. Within Biopython, local sequence alignments are calculated using the Smith–Waterman Algorithm (Smith and Waterman, 1981).

The detection portion of the search method assumes that each gene has a Shine-Dalgarno motif and returns a potential one with an associated alignment score. The next step is to determine if the motif is truly functioning as a Shine-Dalgarno. Two metrics are used To assign whether a Shine-Dalgarno as is functional or not: (1) the binding energy of the SD-aSD interaction and (2) the distance from the gene ORF. If the binding energy of the SD-aSD interaction is greater than 0, it is assumed that the effect of the Shine-Dalgarno in translation is non-functional. Binding energy is tabulated using RNAcofold (v2.4.18) from the ViennaPackage2 (Lorenz et al., 2011). Calculations were performed using the default settings. Temperature for calculations was set to 37^*°*^C. The distance from the gene ORF is based on experimental evidence in *E. coli*, where Shine-Dalgarnosequences were determined to not impact translation when the distance between it and the gene ORF was less than 5 nucleotides or greater than 10 nucleotides (Saito et al., 2020). The data from the Shine-Dalgarno identification for Syn1.0 is presented in **Fig**. S3 and **Fig**. S8. The results from other organisms are also presented in Supplementary Information.

To validate the search method a second search was performed removing the search location bias. The bacterial genome is randomly searched (N=500,000) using the same template sequence. This is then compared to the previously described method to show the enrichment of the SD sequence near the gene ORF and not throughout the genome (See **Fig**. S8). Sequence logos graphs are made using Logomaker (Tareen and Kinney, 2019). Genome architecture diagrams that show motif positions and strand (like those found in **Fig. 1A** and **Fig. 3**) were made with DNAFeatureVeiwer (Zulkower and Rosser, 2020).

#### 4.1.3. Promoter

The promoter search was performed in a similar manner to the Shine-Dalgarno search with some slight modifications. The nature of the interactions between the promoter and RNA polymerase are protein-RNA unlike the RNA-RNA interaction of the Shine-Dalgarno and anti-Shine-Dalgarno. Specifically, the RNA polymerase sigma factor recognizes the promoter region on the DNA, forming the T-RPc complex between the RNAP and DNA (Chen et al., 2020). Promoters have been well characterized in other bacterial organisms to be composed of two regions: the −10 region or Pribnow box characterized by the sequence 5^*′*^-*TANAAT* −3^*′*^, and the −35 region defined by the consensus sequence 5^*′*^-*TTGACA*-3^*′*^. In the related organisms *M. pneumoniae* (Lloréns-Rico et al., 2015) and *M. florum* (Matteau et al., 2020), promoters were identified from sequence, however the −35 region was not observed to be conserved within the two organisms and is not included in our searches. Additionally the −10 region was found to be extended with the following consensus sequence of 5^*′*^-*TGNTANAAT* −3^*′*^ (Lloréns-Rico et al., 2015). Comparison of the RNAP sigma factor sub-units in related organisms, shows similarity between the Syn1.0/Syn3A RNAP sub-unit (See **Fig**. S2), so for the search in this study the template was defined as 5^*′*^-*TANAAT* −3^*′*^.

The promoter search regions was determined using the distance from the transcription start site (Lloréns-Rico et al., 2015). The transcription start site location was determined using the previously detected SD sequence and using the distance between the transcription start site and the SD. In the instance a Shine-Dalgarnois not found, the transcription start site is assumed to be 3 nucleotides upstream from the gene ORF. The similarity matrix used for the search is a equally weighted combination of the transversion matrix, which accounts for the likelihood of conversion between bases of the same type (purines A <-> G, and pyrimidines C <-> T), and 1/-3 match/mismatch matrix used for the SD:

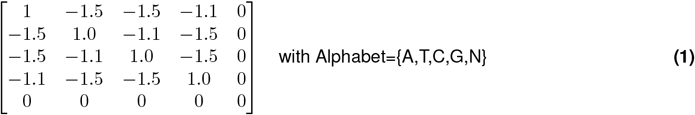

The similarity matrix values a nucleotide match as 1 and mismatch as −1.5 or −1.1. It also accounts for a wildcard with no penalty or gain. A 1/-3 match/mismatch was used for scoring as the searched sequences come from the same organism and would have 99% sequence identity, which is the default matrix used in the NCBI BLAST implementation (Pearson, 2013).

The alphabet contains thymine (T) because the DNA sequence is scanned rather than the RNA sequence. A gap and gap extensions in the alignment are penalized by the maximum score of the respective alignment to ensure no gaps are present in the local alignment. The search was performed via a custom Python script, which used Biopython v1.78 (Cock et al., 2009) to parse and perform sequence alignments. Matrix for scoring was defined using the custom matrix building function of Biopython. Within Biopython local sequence alignments are calculated using the Smith–Waterman Algorithm (Smith and Waterman, 1981).

The Promoter results were validated similar to the SD search. Results for the validation are reported in Figure S14. Results of the search are found in Figure S9A-B and Figures S10-S13. Additional data for the extend region is presented in S15. Functionality within the promoter region is determined via comparison to the random search and applying a threshold for promoter strength that corresponds to a p-value less than 1× 10^*−*5^. This threshold ends up corresponding to roughly a single nucleotide mismatch in the promoter sequence.

#### 4.1.4. Intrinsic Termination Loop

Intrinsic termination loops are characterized by a hairpin loop formation with a stretch of uracil nucleotide downstream of the loop. A decision rule originally parameterized for the related organism, *B. subtilis* (de Hoon et al., 2005) was used to predict termination loops. The decision rule formula is:

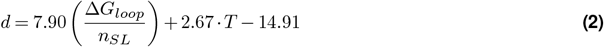

where Δ*G*_*loop*_ is the energy in kcal/mol of the loop formation, *n*_*SL*_ is the number of nucleotides in the stem loop structure, and *T* is the T-stretch score given by:

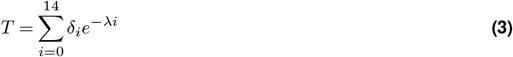

where *δ*_*i*_ is 1 if the nucleotide is a thymine and 0 otherwise. *λ* is a constant fit to experimentally known transcriptional terminators. For each gene ORF, a region downstream the end of the gene ORF is scanned for termination loops until another gene ORF begins or the distance from the gene ORF being searched is larger than 500 nucleotides. de Hoon and coworkers also applied this rule to the parent organism, *Mycoplasma mycoides* subsp. *capri* with a 94.56% selectivity for termination loops. The average values for energy of loop formation, stem length, and T-stretch were compared to those calculated in de Hoon et al. (2005) for the parent organism. The intrinsic termination identification was conducted with a custom Python script obtained from Matteau et al. (2020), which is an updated version of the one used in de Hoon et al. (2005).

### 4.2. Bio-Informatic / Computational Transcription Unit Identification

Transcription units are characterized by the initiation sites (TSSs) and termination sites of (TTSs) of transcription. To predict transcription units, an iterative process of grouping and then separating genes based on strandedness and/or the presence of a TSS / TTS. **Fig. 10** provides a schematic for the transcription unit identification. Scripts to perform the predictions are available at https://github.com/troyb2/MinimalCell_Motif-Identification_RNAseq. Gene ORFs are initially grouped by strand assuming a single RNAP cannot transcribe genes on different strands within a single transcription event (one binding and one unbinding event). Following **Fig. 10**, the gene clusters would be: *A, BC*, and *DEF*. Then TTS or intrinsic termination loops are introduced to separate the genes clustered by strand. At TTS, RNAP diffusion along the DNA is halted and transcription is terminated, so each intrinsic termination loop represents the end of a transcription unit. Finally, TSSs assumed to be identified by the promoter motif because TSS could not be directly identified, are incorporated. Each promoter signifies a potential binding site for RNAP followed by the initiation of transcription. Therefore, each TSS (promoter) starts a TU, which is assumed to continue until a TTS is reached. In cases where the strand separated genes do not have a TSS or TTS detected, a transcription unit is still assigned.

### 4.3. ONT Experimental Methods

#### 4.3.1. Sample Preparation

One mL aliquots of actively growing JCVI-syn1.0 and JCVI-syn3A cells were placed in 25 mL of SP4 + 17% KnockOut serum replacement in 50 mL conical tubes and incubated overnight at 37°C. The cells were harvested in mid-late log phase as indicated by a red-orange color of the phenol red pH indicator in the growth media. The cultures were harvested when the media pH was 7.0. JCVI-syn1.0 cultures achieved this growth stage in 16 hours and JCVI-syn3A cultures in 22 hours. Growth was arrested in the bulk cultures by placing the tubes on wet ice for 10 minutes. Keeping all tube on ice, 1.4 mL of bulk cultures were distributed into fifteen 1.8 mL microfuge tubes per culture. We centrifuged these at 4°C for five minutes at 10,000 × g using a tabletop refrigerated centrifuge to pellet the cells. We aspirated the supernatants from ten of the fifteen tubes. In the remaining five tubes, the supernatant was then used to suspend pellets. The cell suspensions from those five tubes were then used to re-suspend the pellets in five other tubes, and those more concentrated cell suspensions were used to re-suspend the cells in the last five tubes. Now both the Syn1.0 and Syn3A cells were each in five tubes containing concentrated cell suspensions. These tubes were centrifuged for five minutes at 4°C under 10,000 × g with a tabletop refrigerated centrifuge. All supernatants were aspirated completely, and the remaining pellets were snap frozen using liquid nitrogen. Sample tubes were transferred to dry ice and immediately stored at −80°C.

In the next step, cellular RNA was extracted from JCVI-syn1.0 and JCVI-syn3A cell samples using a phenol-chloroform extraction protocol based off procedures adapted from Grünberger et al. (2019b) and the ThermoFisher TRIzolTM Reagent user guide (available online at https://www.thermofisher.com/TFS-Assets/LSG/manuals/trizol_reagent.pdf). Briefly, 1 mL of 4°C TRIzolTM was added to each tube containing frozen cell pellets. We pipetted the liquid up and down multiple times to break up the cell pellet. We incubated the tubes at room temperature for 5 minutes to allow complete disassociation of nucleoprotein complexes. We then added 200 *µ*L of chloroform and mixed the tubes by shaking for 30 seconds. After the tubes incubated at room temperature for 3 minutes, we centrifuged the tubes at 12,000 × g for 15 minutes at 4°C. We pipetted the upper RNA containing aqueous phase without collecting any of the white interphase separating the aqueous from the phenol-chloroform lower phase and placed this solution in fresh 1.8 mL tubes already containing 500 *µ*L isopropanol. The tubes were incubated on ice for 10 minutes and then centrifuged at 12,000 × g for 10 minutes at 4°C. We aspirated and discarded the supernatant off the white gel-like pellets. We inverted the tubes and allowed the pellets to air dry for 15 minutes. The RNA pellets were then dissolved in 20 *µ*L of RNase-free water containing 0.1 mM EDTA. We measured RNA concentrations using a Qubit RNA HS fluorometer assay was used for assessing RNA concentration and either an Agilent Bioanalyzer 2100 (Syn1.0) or an Agilent TapeStation 2200 (Syn3A). The next step was ribosomal RNA (rRNA) depletion.

The original Syn1.0 samples was rRNA depleted using an Invitrogen RiboMinus™ Bacteria 2.0 Transcriptome Isolation Kit. Briefly, 2 *µ*g of total JCVI-syn1.0 RNA was mixed with 100 *µ*l of RiboMinus™ 1X hybridization buffer and 6 *µ*L of RiboMinus™ Pan-Prokaryote Probe Mix. The solution was brought up to 200 *µ*L using RNase-free water. The solution was denatured for 10 minutes at 70°C and then hybridized for 20 minutes at 37°C. The hybridization solution was mixed with RiboMinus™ Magnetic Beads (1 mL of suspension) that had been washed in RNase-free water and re-suspended in 400 *µ*L of RiboMinus™ 1X hybridization buffer. We incubated the mixture at 37°C for 15 minutes. We then placed the sample tube on a RiboMinus™ magnetic stand and transferred the supernatant (600 *µ*L) containing the rRNA-depleted RNA to two new 1.8 mL tubes. To those tubes, each containing 300 *µ*L of RNA, we added premixed Binding Solution Concentrate (400 *µ*L) and Nucleic Acid Binding Beads (10 *µ*L) that came with the kit and pipetted gently to mix. One mL of 100% ethanol was added to each tube, and this homogenous mixture was mixed and then incubated at room temperature for 5 minutes. We placed the tubes back on the magnetic stand for 5 minutes, then aspirated and discarded the supernatant without disturbing the beads. We removed the tubes from the stand, and washed the beads in 300 *µ*L of the kit Wash Solution. We placed the tubes back on the magnetic stand and allowed the solution to clear. Then we aspirated and discard the supernatant without disturbing the bead pellet. After the beads had dried for 5 minutes, we added 25 *µ*L of pre-heated (7°C) nuclease-free water to the beads and incubated the tubes for 1 minute at room temperature to elute the RNA. We returned the tubes to the magnetic stand and carefully collected the supernatant into a new microcentrifuge tube. The next step was addition of a Poly(A) tail to the RNA.

The second Syn1.0 and Syn3A samples were rRNA depleted using the NEBNext® rRNA Depletion Kit for bacteria. One *µ*g of RNA was brought up to 11 *µ*L with RNase-free water. To this we added 2 *µ*L of NEBNext® Bacterial rRNA Depletion Solution and 2 *µ*L of NEBNext® Probe Hybridization Buffer. The reaction was heated to 95°C for 2 minutes and cooled to 22°C at a ramp rate of 0.1°C/sec. This was followed immediately by RNAse H digestion where 2 *µ*L of RNase H Reaction Buffer and 2 *µ*L of NEBNext Thermostable RNase H was added to the hybridized RNA solution to a total volume of 20 *µ*L. This reaction was incubated at 50°C for 30 minutes. Following a bead cleanup Agencourt RNAClean XP beads (Beckman Coulter™, cat No. A63987), the 15 *µ*l of the rRNA solution was Poly(A) tailed at the 3^*′*^ end.

The same Poly(A) tailing protocol was used for Syn1.0 and Syn3A. RNA samples were mixed a premade solution containing 2 *µ*L of 10X E. coli Poly(A) Polymerase Reaction Buffer, 2 *µ*L of ATP and 1 *µ*L of the E.coli Poly(A) Polymerase (New England Biolabs) and incubated at 37°C for 30 minutes. The reaction was stopped by adding EDTA to a final concentration of 10 mM. Following another bead cleanup along with necessary quality control analysis on the samples, RNA was normalized to 9 *µ*L containing 140 ng for Syn1.0 RNA and 50 ng for Syn3A RNA samples.

After normalization, libraries for Oxford Nanopore sequencing were prepared using the ONT Direct RNA Sequencing Kit (SQK-RNA002) with slight modifications. Specifically, 0.5 *µ*L of the RT adapter enzyme (reduced from 1 *µ*L) and 4 *µ*L of the RMX RNA adapter (reduced from 6 *µ*L) were used during library preparation for Syn1.0 and Syn3A. All samples were sequenced using R9.4 flow cells on the ONT GridION platform. Live base-calling was enabled through the recommended MinKNOW (v23.07.5) scripts to generate fast5 files.

Post poly-adenylation of RNA transcripts, the extent of the DNA and buffer contamination in each of the RNA samples were tested by performing standard spectroscopic measurements (NanoDrop One) and using the Qubit 1x dsDNA HS assay kit (ThermoFisher Scientific). Input RNA samples for Oxford Nanopore Direct RNA Sequencing library preparation were finally quantified using the Qubit RNA HS assay kit. RNA concentrations values were used to normalize samples to the desired input volume of 9 *µ*L containing 50-500 ng of input RNA for library preparation. On analyzing ONT run reports for Syn1.0, composite reads generated over 2 runs were 1319.04k with an average N50 of 410. Reads generated for Syn3A over a sing run were 734.08k with N50 430. In all run reports, bases called were 99.99-100%.

#### 4.3.2. Quality Control Analysis

An Agilent Bioanalyzer 2100 assay was used for assessing RNA quality and concentration of the first JCVI-syn1.0 samples both before and after rRNA depletion **Fig**. S34. For the second JCVI-syn1.0 and the JCVI-syn3A samples, both a Qubit RNA HS fluorometer assay and an Agilent TapeStation 2200 were used for assessing RNA concentration post-extraction from minimal cells. Initial Qubit values post RNA extraction were 3000 ^ng^⁄_*µ*L_ for the second Syn1.0 and 1890 ^ng^⁄_*µ*L_ for Syn3A. The Agilent TapeStation 2200 was used throughout the extraction and rRNA depletion steps to assess the quality of the RNA and depletion. TapeStation results gave us an estimate of depletion through the absence of 16S and 23S peaks **Fig**. S35. A Qubit RNA HS fluorometer was used to detect final quantitation of RNA.

Post poly-adenylation of RNA transcripts, the extent of the DNA and buffer contamination in each of the RNA samples were tested by performing standard spectroscopic measurements (NanoDrop One) and using the Qubit 1x dsDNA HS assay kit (ThermoFisher Scientific). Input RNA samples for Oxford Nanopore Direct RNA Sequencing library preparation were finally quantified using the Qubit RNA HS assay kit. RNA concentrations values were used to normalize samples to the desired input volume of 9 *µ*L containing 50-500ng of input RNA for library preparation.

Analysis of Oxford Nanopore Direct RNA sequencing run reports for Syn1.0, composite reads generated over 2 runs were 1319.04k with an average N50 of 410. Reads generated for Syn3A over a single run were 734.08k with N50 430. In all run reports, bases called were 99.99-100% confident.

#### 4.3.3. Analysis of the Oxford Nanopore Long-Read Native Sequencing

Results of the Nanopore reads were collected as individual fast5 files. The individual Nanopore reads are then combined into multi-read files for easier data analysis. Current signals are converted to canonical bases using a technique known as base-calling. For the first JCVI-syn1.0 sequencing, base-calling was performed using ONT standard program Guppy (v6.1.3) Oxford Nanopore Technologies, Ltd. (2022). Base-calling was performed using the RNA trim strategy and with calibration detection and filter enabled. NVIDIA A5000 GPUs were used for the analysis. For the second JCVI-syn1.0 replicate and the JCVI-syn3A experiments, Guppy (v7.0.9,GUI based for the MinKNOW system) was used for base-calling with equivalent parameters. Individual transcripts are then mapped to the respective reference genome (see **Table** S3) with minimap2 (v2.26) (Li, 2018). Standard settings for long read analysis were used for the reference mapping, −x splice, −uf, −k14, −p set to 0.99 for primary and secondary mappings, –MD on to enable MD tag outputting, and −a to output the data as a SAM file. SAM alignment files were further processed with SAMtools (v1.14) (Li et al., 2009) to generate sequencing depth across the genome. Processing statistics for the two Syn1.0 and single Syn3A ONT runs are reported in **Table** S5. Coverage was calculated using the following formula:

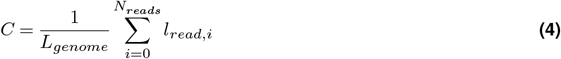

where C is the coverage, *L* is the length of the organism’s genome in nucleotides, *N* is the number of reads in a sample, and *l*_*read,i*_ is the length of read, *i*, in nucleotides. All scripts to run analysis as performed in this study are available at https://github.com/troyb2/MinimalCell_Motif-Identification_RNAseq.

### 4.4. Illumina Experimental Methods

#### 4.4.1. Sample Preparation

Sample preparation was conducted following Grünberger et al. (2019b) as in **Section** 4.3.1. A frozen stock of Syn1.0 strain was provided by the JCVI. The frozen cells stock was scraped and inoculated in 10 mL of SP4+KnockOut media (media components listed in **Table** S8) under BSL2 conditions and incubated in a horizontal rotating shaker at 37°C for about 24 hours until the culture turned to media orange. The SP4 media is made from a two part formulation: part ***I*** –3.5 g Mycoplasma Broth Base, 10 g Bacto Tryptone, 5.3 g Bacto Peptone, and 600 mL distilled water mixed to suspend well, adjusted to pH 7.5, and then autoclaved 15 min at 121°C and part ***II*** –25 mL Glucose 20% w/v stock, 50 mL CMRL 1066 (10X stock w/o phenol red, w/o bicarb, w/o Gln), 14.6 mL sodium bicarbonate 7.5% w/v stock, 5 mL L-glutamine 200 mM stock, 35 mL Yeast extract solution, 100 mL TC Yeastolate 2% w/v stock, 170 mL Serum (heat inactivated FBS, HS) or substitute (KO), 2.5 ml Penicillin G (400,000 U/mL stock), and 1.5 mL Phenol red (1% w/v) filtered and sterilized (0.2 *µ*m) then stored at 4°C. To complete the SP4 medium, combine 1.5 volume of part ***I*** with 1 volume of part ***II***.

RNA purification and depletion steps were conducted using sterile, RNase-free pipette tips (Thomas Scientific 1145N16, VWR 76322-134) and microcentrifuge tubes (Posi-Click 1149K01) on a countertop or biosafety cabinet treated with RNase decontamination solution (Invitrogen AM9780). Solutions apart from those provided with the kits were prepared using DEPC-treated water (Invitrogen 46-2224). RNA was extracted from JCVI-syn1.0 cells using the PureLink RNA Mini Kit (Invitrogen 12183018A) according to the manufacturer’s protocol. Briefly,≤ 1 × 10^9^ cells (2 mL of exponential phase culture) were harvested (4000 × g, 5 min, 4°C) and resuspended in lysozyme solution (100 *µ*L; 10 mM Tris-HCl, pH 8.0, 0.1 mM EDTA, and 20 ^mg^⁄_mL_ lysozyme Egg white (L-040-25, Goldbio). SDS solution (500 *µ*L; 10% w/v) was added and the cells were incubated for 5 min at room temperature (rt). Lysis buffer containing *β*-mercaptoethanol (350 *µ*L) was added and the sample was vortexed and passed six times through an 18-gauge needle. The resulting homogenate was centrifuged (12,000 × g, 2 min, rt) and 100% ethanol (250 *µ*L) was mixed with supernatant separated from the precipitate. The sample was purified through a spin cartridge provided with the kit and eluted with 2 × 100 *µ*L DEPC-treated water. To remove DNA, 1/9 volume of 10X DNase I Buffer (NEB, B0303S) and DNase I (NEB M0303S; 4 U total) were added and the mixture was incubated for 10 min at rt. EDTA (5 mM final concentration) was then added, and the reaction was heat-inactivated for 10 min at 75°C. Quantification with a NanoDrop Lite Spectrophotometer (Thermo Fisher ND-LITE-PR) typically showed yields of 40-60 *µ*g of RNA. Ribosomal RNA was removed from the DNase-treated RNA samples using the RiboMinus Bacteria 2.0 Transcriptome Isolation Kit (Invitrogen A47335) according to the manufacturer’s protocol. Briefly, ≤5 *µ*g of DNase-treated RNA was mixed with 2X Hybridization Buffer (50 *µ*L) and RiboMinus Pan-Prokaryote Probe Mix (3 *µ*L) to a final volume of 100 *µ*L. The RNA/probe mix was denatured for 10 min at 70°C, followed by hybridization of the rRNA and probes for 20 min at 37°C. The RNA/probe mix was added to a microcentrifuge tube containing a 200 *µ*L suspension of RiboMinus Magnetic Beads in 1X Hybridization Buffer and incubated for 15 min at 37°C. The tube was placed on DynaMag-2 Magnet stand (Thermo Fisher 12321D) for 1 min. The supernatant (300 *µ*L) containing the rRNA-depleted RNA was carefully aspirated and mixed with Nucleic Acid Binding Beads (10 *µ*L), Binding Solution Concentrate (400 *µ*L), and 100% ethanol (1 mL). The mixture was incubated for 5 min at rt and placed on the magnetic stand for 5 min. The supernatant was discarded, the beads were washed with Wash Solution (300 *µ*L) and placed on the magnetic stand again. The supernatant was discarded, the beads were air-dried for 5 min, and the RNA was eluted with DEPC-treated water heated to 70°C. The Nucleic Acid Binding Beads were removed using the magnetic stand, and the supernatant was transferred to a fresh microcentrifuge tube.

#### 4.4.2. Quality Control Analysis

Quantification of RNA was conducted using the Qubit RNA High Sensitivity assay (Invitrogen Q32852) and measured on a Qubit fluorometer (Invitrogen). Within-range samples treated with the RiboMinus kit were monitored with an AATI 5200 Fragment Analyzer (Agilent) showing depletion of rRNA-associated peaks. Method name: DNF-472T22 - HS Total RNA 15nt.mthds; Gel prime: No; Full conditioning: Yes; Gel prime to buffer: Yes; Gel selection: Gel 2; Perform prerun: 7.0 kV, 30 sec.; Rinse: No; Marker 1: No; Rinse: Tray: 3, Row: A, Dip count: 2; Sample injection: 6.0 kV, 150 sec.; Separation: 7.0 kV, 31.0 min.; Tray name: Tray-3; Analysis mode: RNA (Eukaryotic). Quality controls plots of RNA libraries have been included in Supplementary Information, **Fig**. S36.

#### 4.4.3. Analysis of the Illumina RNA Sequencing

Processing of the Syn1.0 Illumina data was carried out using a custom NextFlow pipeline. Raw FASTQ reads were assessed for quality using FastQC (v0.11.9) (Andrews, 2010). Adapter trimming and quality filtering were performed using Fastp (v0.20.0) with default parameters (Chen et al., 2018). An additional quality assessment with FastQC was performed on the trimmed reads. The trimmed reads were then aligned to the Syn1.0 reference genome with using BWA-MEM (v0.7.17) (Li, 2013). Post-alignment, the resulting BAM files were sorted and indexed using Samtools (v1.17) (Li et al., 2009). In addition to the NextFlow pipeline, the BAM files were converted to SAM files, and sequencing depth across the genome was extracted with SAMtools (v1.14) (Li et al., 2009). Reads (total and mapped) counts as well as sequencing coverage for the three Syn1.0 experiments are reported in **Table** S5. Coverage was calculated using **Eqn. 4**. All scripts to run analysis as performed in this study are available at https://github.com/troyb2/MinimalCell_Motif-Identification_RNAseq.

### 4.5. PacBio Experimental Methods

#### 4.5.1. Sample Preparation

Sample preparation was conducted in the same manner as in **Section 4.4.1**, except DNase-I treated RNA was purified by spin cartridge (GeneJET RNA Purification Kit, Fisher K0731) instead of heat-inactivated with EDTA. In addition, prior to rRNA depletion, DNase I-treated RNA was polyadenylated using 1-10 *µ*g of the RNA, ATP (1 mM final concentration) (NEB B0756), *E. coli* Poly(A) Polymerase (NEB M0276S), and *E. coli* Poly(A) Polymerase Reaction Buffer (B0276S). The reactions incubated at 37°C for 20 minutes and then purified using the GeneJET RNA spin cartridge.

Ribodepleted and polyadenylated RNAs were purified using an RNA Clean & Concentrator-5 column (Zymo Research, Cat. #R1015) following the protocol designed to remove fragments smaller than 200 nucleotides. Twenty nanograms of purified RNA were converted into complementary DNA (cDNA) using the IsoSeq Express Oligo Kit from Pacific Biosciences. The cDNA molecules were barcoded with the Barcoded Overhang Adaptor Kit 8A and further processed into a library using the SMRTbell Prep Kit 3.0 (Pacific Biosciences). The resulting libraries were pooled at equimolar concentrations, and the pool was sequenced on one SMRT Cell 8M using a PacBio Sequel IIe system in CCS sequencing mode with a 30-hour movie.

#### 4.5.2. Quality Control Analysis

QC was conducted in the same manner as in **Section 4.4.2**. Quality controls plots of RNA libraries have been included in Supplementary Information, **Fig**. S37.

#### 4.5.3. Analysis of the PacBio RNA Sequencing

The built-in PacBio IsoSeq analysis platform was utilized to convert experimental subreads into high fidelity (HiFi) reads for each of the three Syn1.0 replicates. The SMRTLink software (v11.1) (Wenger et al., 2019) was used to classify the subreads, identifying and removing all primers. The results were then clustered based on the CCS algorithm to generate reads. The reads were sorted by quality. HiFi, reads with 99.9% accuracy, were compiled in FASTQ files and then mapped to the Syn1.0 genome using minimap2 (v2.26) (Li, 2018) with the default setting for PacBio HiFi sequencing. SAMtools (v1.14) (Li et al., 2009) was utilized to calculate the sequencing depth across the genome. Reads (total and mapped) counts as well as sequencing coverage for the three Syn1.0 experiments are reported in **Table** S5. Coverage was calculated using **Eqn. 4**. All scripts to run analysis as performed in this study are available at https://github.com/troyb2/MinimalCell_Motif-Identification_RNAseq.

### 4.6. Mass Spectrometry Methods

#### 4.6.1. Sample Preparation – Cell lysis and Protein digestion

Cell pellets of Syn1.0 and Syn3A strains were mixed with 100 *µ*L of SDS-based lysis buffer (4% SDS, 100 mM Tris-HCl, pH 8.0) containing final concentration of 10 mM Tris (2-carboxyethyl) phosphine (TCEP) and 40 mM chloroacetamide (CAA), vortexed for 5 min, and then boiled at 95°C for 10 mins. The lysate samples were then processed for proteomics analysis following the E3technology procedure reported recently (Martin et al., 2024). In brief, the lysates were mixed with 4***x*** volume of 80% acetonitrile (ACN) and then transferred to E3filters (CDS Analytical, Oxford, PA) followed by centrifugation at 3,000 rpm for 2 min. The filters were washed with 500 *µ*L of 80% ACN for three times, and then transferred to clean collection tubes. For protein digestion, around 1.0 *µ*L of trypsin and 200 *µ*L of digestion buffer (50 mM triethylammonium bicarbonate, TEAB) were added to each sample followed by incubation at 37°C for 16-18 hours with gentle shaking (300 rpm/min). After digestion, the filters were first spun at 3,000 rpm for 2 min to collect the flow through, and then eluted sequentially with 200 *µ*L of 0.2% formic acid (FA) in water and 200 *µ*L of 0.2% FA in 50% ACN. The pooled elution was dried in SpeedVac, and stored under −80°C until further analysis.

#### 4.6.2. LC-MS/MS analysis and protein identification

The LC-MS/MS analysis was performed using an Ultimate 3000 RSLCnano system in conjunction with an Orbitrap Eclipse mass spectrometer and FAIMS Pro Interface (Thermo Scientific), similar to the procedures described recently (Sun et al., 2025). In brief, the peptides resuspended in LC buffer A (0.1% FA in water) were first loaded onto a trap column (PepMap100 C18,300 *µ*m × 2 mm, 5 *µ*m particle, Thermo Scientific), and then separated on an analytical column (PepMap100 C18, 50 cm × 75 *µ*m i.d., 3 *µ*m; Thermo Scientific) at a flow of 250 nL/min. A linear LC gradient was applied from 1% to 25% mobile phase B (0.1% formic acid in acetonitrile) over 125 min, followed by an increase to 32% mobile phase B over 10 min. The column was washed with 80% mobile phase B for 5 min, followed by equilibration with mobile phase A for 15 min. For MS analysis, the Detector type was Orbitrap with resolution of 60,000; Precursor MS range (m/z) was 380–980; AGC target was Standard; Maximum injection time mode was Auto. For data-independent acquisition (DIA) MS/MS analysis, the Isolation mode was Quadruple; DIA Window type was Auto and Isolation Window (m/z) was 8 with an overlap of 1; activation type was HCD with fixed collision energy mode (30%); the Detector Type was Orbitrap with a resolution of 15,000; Normalized AGC target (%) was 800, and the Maximum injection time mode was Auto; the Loop Control was 2 sec. For FAIMS compensation voltages (CV) setting, a 3-CV combination (∼40, ∼55, and ∼75) was applied.

For proteome identification, the MS raw data were processed using Spectronaut software (version 19) and a library-free DIA analysis workflow with directDIA+ and the *X. tropicalis* protein database (UniProt 2024 release; 76,225 sequences). Detailed parameters for Pulsar and library generation include: Trypsin/P as specific enzyme with minimum peptide length of 7 amino acids and maximally 2 missed cleavages; toggle N-terminal M turned on; Oxidation on M and Acetyl at protein N-terminus as variable modifications; Carbamidomethyl on C as fixed modification; FDRs at PSM, peptide and protein level all set to 0.01; Quantity MS level set to MS2, and cross-run normalization turned on. The MS raw data associated with this study has been deposited onto the MassIVE Repository with the accession MSV000097054.

#### 4.6.3. Absolute Intracellular Protein Quantification

The Syn1.0 mass spectrometry data provides relative abundances for 735 of the 828 protein coding genes, which can be converted to absolute abundances by scaling the relative amount with the total number of proteins in the cell. The total number of proteins has not been measured directly within Syn1.0, but can be estimated via an approach similar to that used in Breuer et al. (2019). First the dry mass of the cell and protein mass fraction are needed. The protein mass fraction is assumed to be 58.2%, which was measured in the parent organism, *Mycoplasma mycoides* subsp. *capri* (Razin et al., 1963). The dry mass of the cell, 12.8 fg, was also estimated identically to the dry mass of JCVI-syn3A (see Breuer et al. (2019), *Appendix 1*). Accordingly, the average JCVI-syn1.0 cell has approximately 127,000 proteins, which results in a protein density of 2.87 × 10^6^ *proteins/µm*^3^, which is in rough agreement with 3.5 *−* 4.4 × 10^6^ *proteins/µm*^3^ measured in *E. coli* (Milo, 2013). Finally, absolute protein abundances were calculated using the following formula:

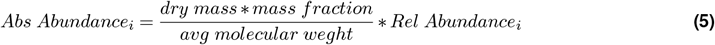

The resulting proteomics are reported in **SI File 1**.

The newly collected Syn3A proteomics were compared to the data set collected in Breuer et al. (2019). There was a slight increase in the total number of detected protein from 428 (Breuer et al., 2019) to 449. When calculating the total number of proteins in the cell, the new data predicts *∼* 100,000 proteins an increase from the 77,000 proteins previously predicted. This change is largely due to an increase in the detection of ribosomal proteins (see **SI File 1**).

### 4.7. Quantification of Cellular Components

#### 4.7.1. Calculation of Dry Mass of a Cell

Absolute abundances were calculated from relative abundances according to the total cellular dry mass, the (dry) mass fraction and the average mass of a cellular specie in question. This approach was used previously for Syn3A (Breuer et al., 2019) and *M. florum* (Matteau et al., 2020). To perform these calculations we first obtained an assumed total cellular dry mass. This approach is modified from the dry mass calculation for JCVI-syn3A used in (Breuer et al., 2019). First the total cellular mass was calculated using the following:

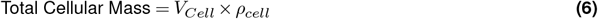

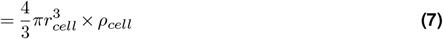

where the *V* is the volume of a sphere with radius, *r*. The density of a cell, *ρ*, is based on measurements in *B. subtilis, E. coli*, and *P. putida* reported by Bratbak and Dundas (1984). The conversion from total cellular mass to the cellular dry mass is achieved by scaling via the cellular dry fraction.

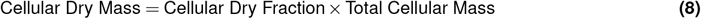

However the cellular dry fraction is unknown value, however it can be defined as:

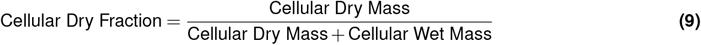

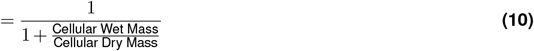

The simplification in **Eqn 10** allows for ratio between the dry and wet mass rather than exact values to calculate the dry fraction. Even without experimental measurements, we used literature values to approximate the ratio between the dry and wet mass via:

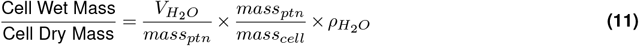

The volume of *H*_2_*O* per mass of protein is taken from a related *Mycoplasma, M. mycoides capri serovar capri PG3* measured in Leblanc and Le Grimellec (1979). The mass of protein is taken from the parent organism of Syn1.0,*M. mycoides* subsp. *capri* (Razin et al., 1963). The true value is likely to deviate however it is the best approximation. The density of water, *ρ*_*H*2_^*O*^, used was 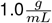. Substituting the results of **Eqn 10** and **Eqn 7** into **Eqn 10** returns a total cellular dry mass, 12.8 fg. All related parameters and calculations are reported in **Table** S9.

#### 4.7.2. Transcriptome Quantification for JCVI-syn1.0

Absolute abundances of the transcriptome were calculated from relative abundances in a similar manner used for the proteome. Starting from the dry mass of the cell (12.8 fg) we determined the dry mass for each type of RNA. The cumulative RNA mass fraction in the parent organism, *Mycoplasma mycoides* subsp. *capri* (Razin et al., 1963), was measured to be 17.3% of the cell. The percent composition of each RNA type within the mass fraction has not been measured in JCVI-syn1.0, but was in the related *Mesoplasma florum*. According to Matteau *et al*., rRNA makes up a majority of the RNA mass in the cell accounting for 80%, while tRNA, mRNA, and sRNA represent 15%, 5%, and *<*1% of the RNA mass respectively (Matteau et al., 2020). These values are in agreement with those from other organisms (Milo et al., 2009; Westermann et al., 2012). We re-scaled these numbers to be out of 100%, holding rRNA fixed at 80%. During the RNAseq library preparation rRNA was depleted, so to account for this a slight modification was performed on the rRNA mass assuming a depletion efficiency of ∼95% according to ribominus kit. Then using the mass and average molecular weight (^g^⁄_mol_) for each RNA type, we calculated the total number of each RNA in a cell. The resulting RNA counts are given in **SI File 5**. The relative abundances are obtained from the RNA sequencing experiments using the transcript per million (TPM) to normalize for each gene. TPM is defined by:

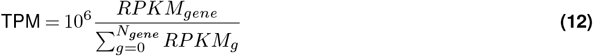

where RPKM or reads per kilobase per million of mapped reads is defined by:

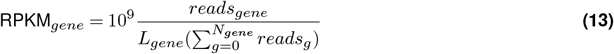

where for a gene of interest *L*_*gene*_ is its length in kbp and *reads*_*gene*_ is the number of reads mapped to the gene. The summation represents the total number of reads mapped to all genes. Reads are assigned to genes using the *htseq-count* software within the HTseq Python analysis package (Anders et al., 2014; Putri et al., 2022). HTseq-count assignment of reads was performed using the *union* mode with reads assigned to multiple feature being split fractionally among all assigned reads (non-unique=fraction). Strandedness is taken to account when applicable (Illumina and ONT). Outputted read counts for each gene are used to solve **Eqn. 13** and **Eqn. 12** resulting absolute RNA abundance data is provided in **SI File 5**.

## 5. Data Availability

Relevant scripts and code are available online in a GitHub repository at https://github.com/troyb2/MinimalCell_Motif-Identification_RNAseq. The repository includes Jupyter notebooks capable of generating a majority of the figures seen in the manuscript, as well as similar figures for other genome regions not shown in this study. JCVI-syn1.0 and JCVI-syn3A proteomics data was deposited on MassIVE Repository, entry MSV000097054 (*insert-link*). RNA sequencing data has been deposited online in the Sequence Read Archive (https://www.ncbi.nlm.nih.gov/sra) database: Syn1.0 and Syn3A ONT – *insert-link*, Syn1.0 Illumina – *insert-link*, and Syn1.0 PacBio – *insert-link*.

(All links will become accessible at the time of publication. Please contact the author for data.)

## 6. Acknowledgments

The authors thank Pratap Venepally (JCVI Rockville) for informative discussion about the ONT experiments and data analysis and Alvaro Hernandez (Roy J Carver Biotechnology Center at UIUC) for carrying out the Illumina and PacBio sequencing. We also thank Kim Wise (JCVI La Jolla) and James Pelletier (Centro Nacional de Biotecnología) for informative discussions. Finally the authors would like to thank Enguang Fu (UIUC) for testing the provided code.

## 6.3. Author Contributions

TAB: conceptualization, methodology, validation, software, formal analysis, writing–original draft, writing–reviewing and editing. JEC: methodology, validation, writing–original draft, and editing. BRG: methodology, validation, software, writing–reviewing and editing. SAG: methodology, validation. YG: methodology, validation. YY: methodology, validation, writing–reviewing and editing. ZRT: methodology, validation. KG: methodology, validation. GJ: methodology, validation. TM: methodology, validation. SS: methodology, validation. CF: validation, software, writing–reviewing and editing. JIG: conceptualization, writing–reviewing and editing, funding acquisition. APM: conceptualization, writing–reviewing and editing, funding acquisition. ZL-S: conceptualization, writing–reviewing and editing, funding acquisition.

## 7. Abbreviations

ONT: Oxford Nanopore Technologies long read RNA sequencing
PacBio: Pacific Biosciences
RNAseq: RNA Sequencing
TU: transcription unit
TSS: transcription start site
TTS: transcription termination site
SD: Shine-Dalgarno
aSD: anti-Shine-Dalgarno
asRNA: anti-sense RNA
ncRNA: non-coding RNA

## References

Anders, S., Pyl, P. T., and Huber, W. (2014). Htseq—a python framework to work with high-throughput sequencing data. Bioinformatics 31, 166–169. doi:10.1093/bioinformatics/btu638

[Dataset] Andrews, S. (2010). Fastqc: A quality control tool for high throughput sequence data

Bailey, T. L., Johnson, J., Grant, C. E., and Noble, W. S. (2015). The MEME suite. Nucleic Acids Research 43, W39–W49. doi:10.1093/nar/gkv416

Balasubramanian, D. and Vanderpool, C. K. (2013). New developments in post-transcriptional regulation of operons by small RNAs. RNA Biology 10, 337–341. doi:10.4161/rna.23696

Benders, G. A., Powell, B. C., and Hutchison, C. A. (2005). Transcriptional analysis of the conserved ftsZ gene cluster in Mycoplasma genitalium and Mycoplasma pneumoniae. Journal of Bacteriology 187, 4542–4551. doi:10.1128/jb.187.13.4542-4551.2005

Bianchi, D. M., Brier, T. A., Poddar, A., Azam, M. S., Vanderpool, C. K., Ha, T., et al. (2020). Stochastic analysis demonstrates the dual role of hfq in chaperoning e. coli sugar shock response. Frontiers in Molecular Biosciences 7. doi:10.3389/fmolb.2020.593826

Bianchi, D. M., Pelletier, J. F., Hutchison, C. A., Glass, J. I., and Luthey-Schulten, Z. (2022). Toward the complete functional characterization of a minimal bacterial proteome. The Journal of Physical Chemistry B 126, 6820–6834. doi:10.1021/acs.jpcb.2c04188

Bittencourt, D. M. d. C., Brown, D. M., Assad-Garcia, N., Romero, M. R., Sun, L., Palhares de Melo, L. A. M., et al. (2024). Minimal bacterial cell jcvi-syn3b as a chassis to investigate interactions between bacteria and mammalian cells. ACS Synthetic Biology 13, 1128–1141. doi:10.1021/acssynbio.3c00513

Björk, G. R., Ericson, J. U., Gustafsson, C. E. D., Hagervall, T. G., Jönsson, Y. H., and Wikström, P. M. (1987). Transfer rna modification. Annual Review of Biochemistry 56, 263–285. doi:10.1146/annurev.bi.56.070187.001403

Boutet, E., Djerroud, S., and Perreault, J. (2022). Small RNAs beyond model organisms: Have we only scratched the surface? International Journal of Molecular Sciences 23, 4448. doi:10.3390/ijms23084448

Bratbak, G. and Dundas, I. (1984). Bacterial dry matter content and biomass estimations. Applied and Environmental Microbiology 48, 755–757. doi:10.1128/aem.48.4.755-757.1984

Breuer, M., Earnest, T. M., Merryman, C., Wise, K. S., Sun, L., Lynott, M. R., et al. (2019). Essential metabolism for a minimal cell. eLife 8. doi:10.7554/eLife.36842

Brewster, R. C., Jones, D. L., and Phillips, R. (2012). Tuning promoter strength through RNA polymerase binding site design in escherichia coli. PLoS Computational Biology 8, e1002811. doi:10.1371/journal.pcbi.1002811

Brown, N. P., Leroy, C., and Sander, C. (1998). MView: a web-compatible database search or multiple alignment viewer. Bioinformatics 14, 380–381. doi:10.1093/bioinformatics/14.4.380

Burgos, R., Weber, M., Gallo, C., Lluch-Senar, M., and Serrano, L. (2021). Widespread ribosome stalling in a genome-reduced bacterium and the need for translational quality control. iScience 24, 102985. doi:10.1016/j.isci.2021.102985

Chen, J., Chiu, C., Gopalkrishnan, S., Chen, A. Y., Olinares, P. D. B., Saecker, R. M., et al. (2020). Stepwise promoter melting by bacterial RNA polymerase. Molecular Cell 78, 275–288.e6. doi:10.1016/j.molcel.2020.02.017

Chen, S., Zhou, Y., Chen, Y., and Gu, J. (2018). fastp: an ultra-fast all-in-one fastq preprocessor. Bioinformatics 34, i884–i890. doi:10.1093/bioinformatics/bty560

Cock, P. J. A., Antao, T., Chang, J. T., Chapman, B. A., Cox, C. J., Dalke, A., et al. (2009). Biopython: freely available python tools for computational molecular biology and bioinformatics. Bioinformatics 25, 1422–1423. doi:10.1093/bioinformatics/btp163

de Crécy-Lagard, V. and Jaroch, M. (2021). Functions of bacterial trna modifications: From ubiquity to diversity. Trends in Microbiology 29, 41–53. doi:10.1016/j.tim.2020.06.010

de Hoon, M. J. L., Makita, Y., Nakai, K., and Miyano, S. (2005). Prediction of transcriptional terminators in bacillus subtilis and related species. PLoS Computational Biology 1, e25. doi:10.1371/journal.pcbi.0010025

Elfmann, C., Dumann, V., van den Berg, T., and Stülke, J. (2024). A new framework for subtiwiki, the database for the model organism bacillus subtilis. Nucleic Acids Research 53, D864–D870. doi:10.1093/nar/gkae957

Feucht, A., Lucet, I., Yudkin, M. D., and Errington, J. (2001). Cytological and biochemical characterization of the ftsa cell division protein of Bacillus subtilis. Molecular Microbiology 40, 115–125. doi:10.1046/j.1365-2958.2001.02356.x

Fischer, S., Dinh, M., Henry, V., Robert, P., Goelzer, A., and Fromion, V. (2021). Bipsim: a flexible and generic stochastic simulator for polymerization processes. Scientific Reports 11. doi:10.1038/s41598-021-92833-5

Fisunov, G., Evsyutina, D., Semashko, T., Arzamasov, A., Manuvera, V., Letarov, A., et al. (2016). Binding site of MraZ transcription factor in mollicutes. Biochimie 125, 59–65. doi:10.1016/j.biochi.2016.02.016

Fondi, M., Pini, F., Riccardi, C., Gemo, P., and Brilli, M. (2022). A selective force driving metabolic genes clustering. BioRxiv doi:10.1101/2022.09.05.506644

Frank, D. N. and Pace, N. R. (1998). Ribonuclease p: Unity and diversity in a trna processing ribozyme. Annual Review of Biochemistry 67, 153–180. doi:10.1146/annurev.biochem.67.1.153

Furlan, M., Delgado-Tejedor, A., Mulroney, L., Pelizzola, M., Novoa, E. M., and Leonardi, T. (2021). Computational methods for rna modification detection from nanopore direct rna sequencing data. RNA Biology 18, 31–40. doi:10.1080/15476286.2021.1978215

Garalde, D. R., Snell, E. A., Jachimowicz, D., Sipos, B., Lloyd, J. H., Bruce, M., et al. (2018). Highly parallel direct RNA sequencing on an array of nanopores. Nature Methods 15, 201–206. doi:10.1038/nmeth.4577

[Dataset] GenBank: Accession No. CP001621.1 (2014). Mycoplasma mycoides subsp. capri str. gm12. NCBI GenBank Entry

[Dataset] GenBank: Accession No. CP002027.1 (2010). synthetic mycoplasma mycoides jcvi-syn1.0. NCBI GenBank Entry

[Dataset] GenBank: Accession No. CP016816.2 (2018). synthetic bacterium jcvi-syn3a. NCBI GenBank Entry

[Dataset] GenBank: Accession No. NC_000912.1 (2001). Mycoplasma pneumoniae m129. NCBI GenBank Entry

[Dataset] GenBank: Accession No. NC_006055.1 (2004). Mesoplasma florum. NCBI GenBank Entry

Gibson, D. G., Glass, J. I., Lartigue, C., Noskov, V. N., Chuang, R.-Y., Algire, M. A., et al. (2010). Creation of a bacterial cell controlled by a chemically synthesized genome. Science 329, 52–56. doi:10.1126/science.1190719

Gilbert, B. R., Thornburg, Z. R., Lam, V., Rashid, F.-Z. M., Glass, J. I., Villa, E., et al. (2021). Generating chromosome geometries in a minimal cell from cryo-electron tomograms and chromosome conformation capture maps. Frontiers in Molecular Biosciences 8. doi:10.3389/fmolb.2021.644133

[Dataset] Glass, J. (2024). doi:10.25345/C5PN89

Grant, C. E., Bailey, T. L., and Noble, W. S. (2011). FIMO: scanning for occurrences of a given motif. Bioinformatics 27, 1017–1018. doi:10.1093/bioinformatics/btr064

Grünberger, F., Knüppel, R., Jüttner, M., Fenk, M., Borst, A., Reichelt, R., et al. (2019a). Exploring prokaryotic transcription, operon structures, rRNA maturation and modifications using nanopore-based native RNA sequencing. bioRxiv doi:10.1101/2019.12.18.880849

Grünberger, F., Reichelt, R., Bunk, B., Spröer, C., Overmann, J., Rachel, R., et al. (2019b). Next generation dna-seq and differential rna-seq allow re-annotation of the pyrococcus furiosus dsm 3638 genome and provide insights into archaeal antisense transcription. Frontiers in Microbiology 10. doi:10.3389/fmicb.2019.01603

Güell, M., van Noort, V., Yus, E., Chen, W.-H., Leigh-Bell, J., Michalodimitrakis, K., et al. (2009). Transcriptome complexity in a genome-reduced bacterium. Science 326, 1268–1271. doi:10.1126/science.1176951

Haas, D., Thamm, A. M., Sun, J., Huang, L., Sun, L., Beaudoin, G. A. W., et al. (2022). Metabolite damage and damage control in a minimal genome. mBio 13. doi:10.1128/mbio.01630-22

Harte, N., Silventoinen, V., Quevillon, E., Robinson, S., Kallio, K., Fustero, X., et al. (2004). Public web-based services from the european bioinformatics institute. Nucleic Acids Research 32, W3–W9. doi:10.1093/nar/gkh405

Herzel, L., Stanley, J. A., Yao, C.-C., and Li, G.-W. (2022). Ubiquitous mRNA decay fragments in E. coli redefine the functional transcriptome. Nucleic Acids Research 50, 5029–5046. doi:10.1093/nar/gkac295

Hockenberry, A. J., Jewett, M. C., Amaral, L. A. N., and Wilke, C. O. (2018). Within-gene shine–dalgarno sequences are not selected for function. Molecular Biology and Evolution 35, 2487–2498. doi:10.1093/molbev/msy150

Hockenberry, A. J., Pah, A. R., Jewett, M. C., and Amaral, L. A. N. (2017). Leveraging genome-wide datasets to quantify the functional role of the anti-shine–dalgarno sequence in regulating translation efficiency. Open Biology 7, 160239. doi:10.1098/rsob.160239

Hutchison, C. A., Chuang, R.-Y., Noskov, V. N., Assad-Garcia, N., Deerinck, T. J., Ellisman, M. H., et al. (2016). Design and synthesis of a minimal bacterial genome. Science 351. doi:10.1126/science.aad6253

Hutchison, C. A., Merryman, C., Sun, L., Assad-Garcia, N., Richter, R. A., Smith, H. O., et al. (2019). Polar effects of transposon insertion into a minimal bacterial genome. Journal of Bacteriology 201. doi:10.1128/jb.00185-19

Illumina, I. (2011). Quality scores for next-generation sequencing. Technical Note: Informatics 31, 39–109

Ireland, W. T., Beeler, S. M., Flores-Bautista, E., McCarty, N. S., Röschinger, T., Belliveau, N. M., et al. (2020). Deciphering the regulatory genome of escherichia coli, one hundred promoters at a time. eLife 9. doi:10.7554/elife.55308

Ju, X., Li, S., Froom, R., Wang, L., Lilic, M., Delbeau, M., et al. (2024). Incomplete transcripts dominate the mycobacterium tuberculosis transcriptome. Nature 627, 424–430. doi:10.1038/s41586-024-07105-9

Justice, I., Kiesel, P., Safronova, N., von Appen, A., and Saenz, J. P. (2024). A tuneable minimal cell membrane reveals that two lipid species suffice for life. Nature Communications 15. doi:10.1038/s41467-024-53975-y

Kaminishi, T., Wilson, D. N., Takemoto, C., Harms, J. M., Kawazoe, M., Schluenzen, F., et al. (2007). A snapshot of the 30s ribosomal subunit capturing mRNA via the shine-dalgarno interaction. Structure 15, 289–297. doi:10.1016/j.str.2006.12.008

Kreuzer, K. D. and Henkin, T. M. (2018). The t-box riboswitch: trna as an effector to modulate gene regulation. Microbiology Spectrum 6. doi:10.1128/microbiolspec.rwr-0028-2018

Kumar, S. and Mohapatra, T. (2021). Deciphering epitranscriptome: Modification of mrna bases provides a new perspective for post-transcriptional regulation of gene expression. Frontiers in Cell and Developmental Biology 9. doi:10.3389/fcell.2021.628415

Lachance, J.-C., Matteau, D., Brodeur, J., Lloyd, C. J., Mih, N., King, Z. A., et al. (2021). Genome-scale metabolic modeling reveals key features of a minimal gene set. Molecular Systems Biology 17. doi:10.15252/msb.202010099

Lachance, J.-C., Rodrigue, S., and Palsson, B. O. (2019). Minimal cells, maximal knowledge. eLife 8. doi:10.7554/elife.45379

Le, T. B. K., Imakaev, M. V., Mirny, L. A., and Laub, M. T. (2013). High-resolution mapping of the spatial organization of a bacterial chromosome. Science 342, 731–734. doi:10.1126/science.1242059

Leblanc, G. and Le Grimellec, C. (1979). Active k+ transport in mycoplasma mycoides var. capri. net and unidirectional k+ movements. Biochimica et Biophysica Acta (BBA) - Biomembranes 554, 156–167. doi:10.1016/0005-2736(79)90015-4

Li, H. (2013). Aligning sequence reads, clone sequences and assembly contigs with bwa-mem doi:10.48550/ARXIV.1303.3997

Li, H. (2018). Minimap2: pairwise alignment for nucleotide sequences. Bioinformatics 34, 3094–3100. doi:10.1093/bioinformatics/bty191

Li, H., Handsaker, B., Wysoker, A., Fennell, T., Ruan, J., Homer, N., et al. (2009). The sequence alignment/map format and SAMtools. Bioinformatics 25, 2078–2079. doi:10.1093/bioinformatics/btp352

Liu, H., Begik, O., Lucas, M. C., Ramirez, J. M., Mason, C. E., Wiener, D., et al. (2019). Accurate detection of m6a RNA modifications in native RNA sequences. Nature Communications 10. doi:10.1038/s41467-019-11713-9

Liu, H., Begik, O., and Novoa, E. M. (2021). EpiNano: Detection of m6a RNA modifications using oxford nanopore direct RNA sequencing. In Methods in Molecular Biology (Springer US). 31–52. doi:10.1007/978-1-0716-1374-0_3

Lloréns-Rico, V., Cano, J., Kamminga, T., Gil, R., Latorre, A., Chen, W.-H., et al. (2016). Bacterial antisense RNAs are mainly the product of transcriptional noise. Science Advances 2. doi:10.1126/sciadv.1501363

Lloréns-Rico, V., Lluch-Senar, M., and Serrano, L. (2015). Distinguishing between productive and abortive promoters using a random forest classifier in mycoplasma pneumoniae. Nucleic Acids Research 43, 3442–3453. doi:10.1093/nar/gkv170

Lorenz, R., Bernhart, S. H., zu Siederdissen, C. H., Tafer, H., Flamm, C., Stadler, P. F., et al. (2011). ViennaRNA package 2.0. Algorithms for Molecular Biology 6. doi:10.1186/1748-7188-6-26

Macklin, D. N., Ahn-Horst, T. A., Choi, H., Ruggero, N. A., Carrera, J., Mason, J. C., et al. (2020). Simultaneous cross-evaluation of heterogeneous e. coli datasets via mechanistic simulation. Science 369. doi:10.1126/science.aav3751

Madeira, F., Pearce, M., Tivey, A. R. N., Basutkar, P., Lee, J., Edbali, O., et al. (2022). Search and sequence analysis tools services from EMBL-EBI in 2022. Nucleic Acids Research 50, W276–W279. doi:10.1093/nar/gkac240

Mandal, M. and Breaker, R. R. (2004). Gene regulation by riboswitches. Nature Reviews Molecular Cell Biology 5, 451–463. doi:10.1038/nrm1403

Maniloff, J., McElhaney, R. M., Baseman, J., and Lloyd, F. (1992). Mycoplasmas: Molecular biology and pathogenesis (American Society for Microbiology)

Mariscal, A. M., Kakizawa, S., Hsu, J. Y., Tanaka, K., González-González, L., Broto, A., et al. (2018). Tuning gene activity by inducible and targeted regulation of gene expression in minimal bacterial cells. ACS Synthetic Biology 7, 1538–1552. doi:10.1021/acssynbio.8b00028

Mars, R. A. T., Nicolas, P., Denham, E. L., and van Dijl, J. M. (2016). Regulatory RNAs in bacillus subtilis: a gram-positive perspective on bacterial RNA-mediated regulation of gene expression. Microbiology and Molecular Biology Reviews 80, 1029–1057. doi:10.1128/mmbr.00026-16

Martin, K. R., Le, H. T., Abdelgawad, A., Yang, C., Lu, G., Keffer, J. L., et al. (2024). Development of an efficient, effective, and economical technology for proteome analysis. Cell Reports Methods 4, 100796. doi:10.1016/j.crmeth.2024.100796

Matteau, D., Lachance, J.-C., Grenier, F., Gauthier, S., Daubenspeck, J. M., Dybvig, K., et al. (2020). Integrative characterization of the near-minimal bacterium Mesoplasma florum. Molecular Systems Biology 16. doi:10.15252/msb.20209844

Milo, R. (2013). What is the total number of protein molecules per cell volume? a call to rethink some published values. BioEssays 35, 1050–1055. doi:10.1002/bies.201300066

Milo, R., Jorgensen, P., Moran, U., Weber, G., and Springer, M. (2009). Bionumbers—the database of key numbers in molecular and cell biology. Nucleic Acids Research 38, D750–D753. doi:10.1093/nar/gkp889

Moger-Reischer, R. Z., Glass, J. I., Wise, K. S., Sun, L., Bittencourt, D. M. C., Lehmkuhl, B. K., et al. (2023). Evolution of a minimal cell. Nature 620, 122–127. doi:10.1038/s41586-023-06288-x

Morowitz, H. J. (1984). The completeness of molecular biology. Israel journal of medical sciences 20, 750–753

Mortazavi, A., Williams, B. A., McCue, K., Schaeffer, L., and Wold, B. (2008). Mapping and quantifying mammalian transcriptomes by rna-seq. Nature Methods 5, 621–628. doi:10.1038/nmeth.1226

Mukherjee, S., Mandal, S. D., Gupta, N., Drory-Retwitzer, M., Barash, D., and Sengupta, S. (2019). RiboD: a comprehensive database for prokaryotic riboswitches. Bioinformatics 35, 3541–3543. doi:10.1093/bioinformatics/btz093

Nagalakshmi, U., Wang, Z., Waern, K., Shou, C., Raha, D., Gerstein, M., et al. (2008). The transcriptional landscape of the yeast genome defined by rna sequencing. Science 320, 1344–1349. doi:10.1126/science.1158441

O’Reilly, F. J., Xue, L., Graziadei, A., Sinn, L., Lenz, S., Tegunov, D., et al. (2020). In-cell architecture of an actively transcribing-translating expressome. Science 369, 554–557. doi:10.1126/science.abb3758

Pearson, W. R. (2013). Selecting the right similarity-scoring matrix. Current Protocols in Bioinformatics 43. doi:10.1002/0471250953.bi0305s43

Pelletier, J. F., Sun, L., Wise, K. S., Assad-Garcia, N., Karas, B. J., Deerinck, T. J., et al. (2021). Genetic requirements for cell division in a genomically minimal cell. Cell 184, 2430–2440.e16. doi:10.1016/j.cell.2021.03.008

Putri, G. H., Anders, S., Pyl, P. T., Pimanda, J. E., and Zanini, F. (2022). Analysing high-throughput sequencing data in python with htseq 2.0. Bioinformatics 38, 2943–2945. doi:10.1093/bioinformatics/btac166

Rae, C. D., Gordiyenko, Y., and Ramakrishnan, V. (2019). How a circularized tmrna moves through the ribosome. Science 363, 740–744. doi:10.1126/science.aav9370

Razin, S., Argaman, M., and Avigan, J. (1963). Chemical composition of mycoplasma cells and membranes. Journal of General Microbiology 33, 477–487. doi:10.1099/00221287-33-3-477

Rhoads, A. and Au, K. F. (2015). Pacbio sequencing and its applications. Genomics, Proteomics & Bioinformatics 13, 278–289. doi:10.1016/j.gpb.2015.08.002

Rueda, S., Vicente, M., and Mingorance, J. (2003). Concentration and assembly of the division ring proteins ftsz, ftsa, and zipa during the Escherichia coli cell cycle. Journal of Bacteriology 185, 3344–3351. doi:10.1128/jb.185.11.3344-3351.2003

Safronova, N., Junghans, L., and Saenz, J. P. (2024). Temperature change elicits lipidome adaptation in the simple organisms mycoplasma mycoides and jcvi-syn3b. Cell Reports 43, 114435. doi:10.1016/j.celrep.2024.114435

Saito, K., Green, R., and Buskirk, A. R. (2020). Translational initiation in e. coli occurs at the correct sites genome-wide in the absence of mRNA-rRNA base-pairing. eLife 9. doi:10.7554/elife.55002

Samuelsson, T., Guindy, Y. S., Lustig, F., Borén, T., and Lagerkvist, U. (1987). Apparent lack of discrimination in the reading of certain codons in mycoplasma mycoides. Proceedings of the National Academy of Sciences 84, 3166–3170. doi:10.1073/pnas.84.10.3166

Sandberg, T. E., Wise, K. S., Dalldorf, C., Szubin, R., Feist, A. M., Glass, J. I., et al. (2023). Adaptive evolution of a minimal organism with a synthetic genome. iScience 26, 107500. doi:10.1016/j.isci.2023.107500

Satam, H., Joshi, K., Mangrolia, U., Waghoo, S., Zaidi, G., Rawool, S., et al. (2023). Next-generation sequencing technology: Current trends and advancements. Biology 12, 997. doi:10.3390/biology12070997

Sayers, E. W., Bolton, E. E., Brister, J. R., Canese, K., Chan, J., Comeau, D. C., et al. (2021). Database resources of the national center for biotechnology information. Nucleic Acids Research 50, D20–D26. doi:10.1093/nar/gkab1112

Shah, S. B., Hill, A. M., Wilke, C. O., and Hockenberry, A. J. (2022). Generating dynamic gene expression patterns without the need for regulatory circuits. PLOS ONE 17, e0268883. doi:10.1371/journal.pone.0268883

Shine, J. and Dalgarno, L. (1974). The 3’-terminal sequence of escherichia coli 16s ribosomal RNA: Complementarity to nonsense triplets and ribosome binding sites. Proceedings of the National Academy of Sciences 71, 1342–1346. doi:10.1073/pnas.71.4.1342

Sievers, F., Wilm, A., Dineen, D., Gibson, T. J., Karplus, K., Li, W., et al. (2011). Fast, scalable generation of high-quality protein multiple sequence alignments using clustal omega. Molecular Systems Biology 7, 539. doi:10.1038/msb.2011.75

Smith, T. and Waterman, M. (1981). Identification of common molecular subsequences. Journal of Molecular Biology 147, 195–197. doi:10.1016/0022-2836(81)90087-5

Sobti, M., Walshe, J. L., Wu, D., Ishmukhametov, R., Zeng, Y. C., Robinson, C. V., et al. (2020). Cryo-EM structures provide insight into how e. coli f1fo ATP synthase accommodates symmetry mismatch. Nature Communications 11. doi:10.1038/s41467-020-16387-2

Soneson, C., Yao, Y., Bratus-Neuenschwander, A., Patrignani, A., Robinson, M. D., and Hussain, S. (2019). A comprehensive examination of nanopore native RNA sequencing for characterization of complex transcriptomes. Nature Communications 10. doi:10.1038/s41467-019-11272-z

Stark, R., Grzelak, M., and Hadfield, J. (2019). Rna sequencing: the teenage years. Nature Reviews Genetics 20, 631–656. doi:10.1038/s41576-019-0150-2

Storz, G., Vogel, J., and Wassarman, K. M. (2011). Regulation by small RNAs in bacteria: Expanding frontiers. Molecular Cell 43, 880–891. doi:10.1016/j.molcel.2011.08.022

Sun, G., DeFelice, M. M., Gillies, T. E., Ahn-Horst, T. A., Andrews, C. J., Krummenacker, M., et al. (2024). Cross-evaluation of e. coli’s operon structures via a whole-cell model suggests alternative cellular benefits for low-versus high-expressing operons. Cell Systems doi:10.1016/j.cels.2024.02.002

Sun, J., Xu, X., Wei, S., and Yu, Y. (2025). In-cell proteomics enables high-resolution spatial and temporal mapping of early xenopus tropicalis embryos doi:10.1101/2025.05.23.655823

Taggart, J. C., Dierksheide, K. J., LeBlanc, H. J., Lalanne, J.-B., Durand, S., Braun, F., et al. (2025). A high-resolution view of rna endonuclease cleavage in bacillus subtilis. Nucleic Acids Research 53. doi:10.1093/nar/gkaf030

Taggart, J. C., Lalanne, J.-B., and Li, G.-W. (2021). Quantitative control for stoichiometric protein synthesis. Annual Review of Microbiology 75, 243–267. doi:10.1146/annurev-micro-041921-012646

Taggart, J. C., Zauber, H., Selbach, M., Li, G.-W., and McShane, E. (2020). Keeping the proportions of protein complex components in check. Cell Systems 10, 125–132. doi:10.1016/j.cels.2020.01.004

Tamames, J. (2001). Evolution of gene order conservation in prokaryotes. Genome Biology 2. doi:10.1186/gb-2001-2-6-research0020

Taniguchi, Y., Choi, P. J., Li, G.-W., Chen, H., Babu, M., Hearn, J., et al. (2010). Quantifying e. coli proteome and transcriptome with single-molecule sensitivity in single cells. Science 329, 533–538. doi:10.1126/science.1188308

Tareen, A. and Kinney, J. B. (2019). Logomaker: beautiful sequence logos in python. Bioinformatics 36, 2272–2274. doi:10.1093/bioinformatics/btz921

Tarnowski, M. J. and Gorochowski, T. E. (2022). Massively parallel characterization of engineered transcript isoforms using direct RNA sequencing. Nature Communications 13. doi:10.1038/s41467-022-28074-5

Tavakoli, S., Nabizadeh, M., Makhamreh, A., Gamper, H., McCormick, C. A., Rezapour, N. K., et al. (2023). Semi-quantitative detection of pseudouridine modifications and type i/ii hypermodifications in human mrnas using direct long-read sequencing. Nature Communications 14. doi:10.1038/s41467-023-35858-w

TheUniprotConsortium (2020). UniProt: the universal protein knowledgebase in 2021. Nucleic Acids Research 49, D480–D489. doi:10.1093/nar/gkaa1100

Thornburg, Z. R., Bianchi, D. M., Brier, T. A., Gilbert, B. R., Earnest, T. M., Melo, M. C., et al. (2022). Fundamental behaviors emerge from simulations of a living minimal cell. Cell 185, 345–360.e28. doi:10.1016/j.cell.2021.12.025

Thornburg, Z. R., Melo, M. C. R., Bianchi, D., Brier, T. A., Crotty, C., Breuer, M., et al. (2019). Kinetic modeling of the genetic information processes in a minimal cell. Frontiers in Molecular Biosciences 6. doi:10.3389/fmolb.2019.00130

Trussart, M., Yus, E., Martinez, S., Baù, D., Tahara, Y. O., Pengo, T., et al. (2017). Defined chromosome structure in the genome-reduced bacterium mycoplasma pneumoniae. Nature Communications 8. doi:10.1038/ncomms14665

Walker, J. E., Saraste, M., and Gay, N. J. (1984). The UNC operon nucleotide sequence, regulation and structure of ATP-synthase. Biochimica et Biophysica Acta (BBA) - Reviews on Bioenergetics 768, 164–200. doi:10.1016/0304-4173(84)90003-x

Webster, M. W., Chauvier, A., Rahil, H., Graziadei, A., Charles, K., Miropolskaya, N., et al. (2024). Molecular basis of mrna delivery to the bacterial ribosome. Science 386. doi:10.1126/science.ado8476

Wenger, A. M., Peluso, P., Rowell, W. J., Chang, P.-C., Hall, R. J., Concepcion, G. T., et al. (2019). Accurate circular consensus long-read sequencing improves variant detection and assembly of a human genome. Nature Biotechnology 37, 1155–1162. doi:10.1038/s41587-019-0217-9

Westermann, A. J., Gorski, S. A., and Vogel, J. (2012). Dual rna-seq of pathogen and host. Nature Reviews Microbiology 10, 618–630. doi:10.1038/nrmicro2852

Wilhelm, B. T., Marguerat, S., Watt, S., Schubert, F., Wood, V., Goodhead, I., et al. (2008). Dynamic repertoire of a eukaryotic transcriptome surveyed at single-nucleotide resolution. Nature 453, 1239–1243. doi:10.1038/nature07002

Wilson, R. C. and Doudna, J. A. (2013). Molecular mechanisms of rna interference. Annual Review of Biophysics 42, 217–239. doi:10.1146/annurev-biophys-083012-130404

Zhao, J., Zhang, H., Qin, B., Nikolay, R., He, Q.-Y., Spahn, C. M. T., et al. (2019). Multifaceted stoichiometry control of bacterial operons revealed by deep proteome quantification. Frontiers in Genetics 10. doi:10.3389/fgene.2019.00473

Zulkower, V. and Rosser, S. (2020). DNA features viewer: a sequence annotation formatting and plotting library for python. Bioinformatics 36, 4350–4352. doi:10.1093/bioinformatics/btaa213

