## Supplemental Information for "Unraveling the Transcriptional Landscape within a Minimized Bacterium via Comparative Analysis"

### S1. Supplementary Information

List of Supplementary Files:

**SI File 1: Syn1_Syn3A_Proteomics_Comparison.xlsx**, a comparison of the relative change in Proteomics for Syn1.0 and Syn3A

**SI File 2: corrected_syn3a_rRNA.fa**, a fasta file with corrected sequences and coordinate for 16S and 23S ribosomal RNA in Syn3A. Coordinates correspond to the JCVI-syn3A GenBank entry(CP016816.2)

**SI File 3: Motifs_Identifications.xlsx**, excel file with motif identifications for all JCVI-syn1.0, JCVI-syn3A, *M. mycoides* subsp. *capri*, *M. florum*, and *M. pneumoniae*

**SI File 4: Predicted_Transcription_Units.xlsx**, transcription unit predictions for JCVI-syn1.0 based on *de novo* and long-read RNAseq TSS and TTS results

**SI File 5: RNA_abundances.xlsx**, excel file with relative and absolute RNA abundances calculated using the JCVI-syn1.0 Illumina, ONT, and PacBio sequencing results

**SI File 6: Illumina_RNASeq_Genome_Coverage.pdf**, full sequencing depth measured across the entire genome for all of the Illumina RNAseq experiments

**SI File 7: PacBio_RNASeq_Genome_Coverage.pdf**, full sequencing depth measured across the entire genome for all of the PacBio RNAseq experiments

**SI File 8: ONT_RNASeq_Genome_Coverage.pdf**, full sequencing depth measured across the entire genome for all of the ONT RNAseq experiments

**SI File 9: ONT_RNASeq_Genome_Coverage_Syn3A.pdf**, full sequencing depth measured across the entire genome for all of the ONT RNAseq experiments for JCVI-syn3A

### S2. Genetic Motif Detection

#### S2.1. Shine-Dalgarno Identification

The Shine-Dalgarno detection was performed using a bio-informatics approach employing sequencing alignment to a set of organisms specific template sequences. Given the physical nature of the SD-aSD interaction, RNA hybridization between the $3'$ end of the 16S rRNA and $5'$ end of the mRNA Shine and Dalgarno (1974), we chose to use the $3'$ end of the 16S rRNA to define the template sequence on an organism-by-organism basis. Previous studies have defined the consensus SD sequence as $5'$-*GGAGG*-$3'$ (Hockenberry et al., 2017, 2018; Matteau et al., 2020). However we chose to use a longer sequence due to structural and orthogonal ribosomes studies which indicate more nucleotides are involved in the SD-aSD interaction (Kaminishi et al., 2007; Saito et al., 2020). All SD sequences used for the search are recorded in **Table** S4. There is significant agreement between the template sequences used, which is expected given the sequence alignment of the 16S rRNA within the organisms of the study (See **Fig.** S1). Using the defined consensus sequence, two searches were performed for each organism.

The first was performed in a region 25 nucleotides upstream of each gene ORF. The distance was chosen based on the location of SDs previously observed (Hockenberry et al., 2017, 2018; Matteau et al., 2020; Saito et al., 2020). The benefit of this approach is each gene has been assigned a potential SD with a SD strength defined by the score of the sequence alignment. The results for the SD detection within Syn1.0 are reported in **Fig.** S3. Data for other organisms is found in **Figs.** S4-S7. In all cases the search returned motifs with relatively good agreement with the consensus template sequence (**Fig.** S3A). Additionally, the strength (alignment score) of the SD sequences correlate reasonably well ($R^2 \approx 0.8$) with binding energies of the SD-aSD hybridization (**Fig.** S3C). Binding energies were calculated with RNAcoFold (Lorenz et al., 2011), and have previously been used to identify SD sequences in Hockenberry et al. (2017). The agreement between alignment scores and binding energies was expected and confirms our search methodology is reasonable. As a result of the method all genes have a potential SD, so additional characterization with respect to the functionality of each SD is needed. Functionality was assigned by two factors: the binding energy of the SD-aSD and the spacer region between the SD and its respective genes start codon. Binding energy greater than 0 kcal/mol are assumed to be non-functional due to the spontaneous nature of the SD-aSD interaction. The spacer region has been observed in *E. coli* to impact the SD's role in increasing translation initiation efficiency (Saito et al., 2020). SDs starting less than 5 or grater than 10 nucleotides away from the start codon of their respective gene do not have improved translation initiation efficiency. We assigned the functionality of our candidate SDs based on the restrictions on the binding energies ($\Delta G \leq 0$) and spacer region distances ($5 \leq d \leq 10$), these are displayed in **Fig.** S3B. The majority of the SD functionality assignments are due to the restrictions on the spacer region.

The second search was performed non-specifically by randomly sampling locations for detection throughout the entire genome to confirm enrichment of the SD near the start of genes. See Materials and Methods–**Section** 4.1 for a more comprehensive description of the protocol for the SD motif detection. Values within the non-specific search are significantly lower than those found in the biased (targeted) SD search. P-values between the two data sets are lower than $1 \times 10^{-4}$ (P value=0.0), which is the threshold used within previously published motif detection software Grant et al. (2011); Bailey et al. (2015). The SD detected have been confirmed to be statistically significant with respect to the rest of the genome and have been assigned functionality using a combination of previously published metrics. The functional SD within the organism are expected to have a enhancing impact on translation output observed via quantitative characterization of translational activity such as protein expression levels (Hockenberry et al., 2017) or ribosome profiling Saito et al. (2020). However within the functional SD, we do not see strong correlation ($R^2 < 0.2$) between SD strength and protein abundances (See **Fig.** S8). There is a general linear relationship implying SD play a role but are not the sole factor in protein abundances. We hypothesize the impact of the SD to be more pronounced due to fewer regulatory genes being present in Syn3A compared to Syn1.0 as a result of the genome reduction.

##### S2.2. Promoter Identification

Promoters were identified in a similar manner to the SD, using a template sequence to perform a targeted sequence alignment in order to identify the motifs. Additional details on the methodology can be found within the Materials and Methods section. The promoter motif is a region on the DNA recognized by the sigma factor sub-unit of RNAP. Promoters are found upstream from the gene start, farther upstream past the SD and transcription start site. The promoter is composed of two well-characterized regions: the -10 region or Pribnow box characterized by the consensus sequence $5'$-*TANAAT*-$3'$, and the -35 region defined by the consensus sequence $5'$-*TTGACA*-$3'$. However in organisms related to Syn1.0, the -35 region is not commonly observed, and the -10 region is found to have an extended region making the -10 consensus sequence $5'$-*TGNTANAAT*-$3'$ (Lloréns-Rico et al., 2015; Matteau et al., 2020). As a result, we have chosen only to search for the -10 region of the promoter motif within this work. Using the approximate location of the promoter motifs and its consensus sequence, potential promoters were identified. Distances between the potential promoters and their respective gene start codon agree well with an analogous study performed in *M. florum* (Matteau et al., 2020). The result for Syn1.0 are reported in **Fig.** S9 and all other **Figs.** S10-S13.

Unlike the SD, the promoter-RNAP binding is DNA-protein interaction, in contrast to the RNA–RNA interaction of the SD-aSD, and as a result it is more difficult to determine the correlation between promoter strength and binding energy. Therefore, we use the non-specific search of the genome for the promoter as a way to assign functionality (see **Fig.** S14 and **Fig.** S15). We have chosen this approach because promoter activity data has not been measured in Syn1.0 or Syn3A, and although data is available in other organisms, promoter activity can varying depending on conditions (Brewster et al., 2012; Ireland et al., 2020). Our approach allows for the detection of all possible promoters rather than those that may be active in a given condition. Functionality was assigned using a threshold p-value less than $1 \times 10^{-5}$, corresponding to a single nucleotide mismatch in the promoter sequence.

##### S2.3. Intrinsic Termination Loop

Intrinsic termination loops, the signal for the end of transcription in Gram-positive bacteria, were detected using a decision rule with parameters developed for the parent organism, *M. mycoides* subsp. *capri* in de Hoon et al. (2005). The decision rule takes into account the $\Delta$G of the loop structure as well as the stretch of T nucleotides after the stem loop (see Materials and Methods section for more detail). The resulting terminators found within the organism agree with average values for $\Delta$G of the stem loop, stem loop size, and T-stretch length reported in de Hoon et al. (2005) (Table S1). For each organism examined, approximately 40% of genes are found with a terminator downstream. *M. pneumoniae* showed the lowest percentage (~21%) of genes with intrinsic termination loops. The relatively low fraction of genes was previously observed in de Hoon et al. (2005) (DataSet4). We hypothesis a reason for the lower fraction of identified terminators is the functional differences in transcription such as the observed transcription-translation coupling (O'Reilly et al., 2020) in *M. pneumoniae* compared to the other organisms. Distances between the gene and intrinsic termination loop are reported for Syn1.0 in **Fig.** S9C and data for other organisms are in **Figs.** S10-S13C.

#### S3. RNA Modification Detection

Bacterial RNA can undergo biochemical modifications, reversible enzyme-catalyzed processes where canonical bases (A,C,G, and U) are converted into modified forms (Kumar and Mohapatra, 2021). The set of all RNA species and their modified forms is known as the epitranscriptome. Modifications have been observed in most types of RNA, however the exact function and cause of the modifications has yet to be fully explained (Kumar and Mohapatra, 2021; Furlan et al., 2021). Those found in rRNA and tRNA have been the most characterized (impacting the RNA structure and function) (Björk et al., 1987), but ongoing has begun to highlight the importance of modifications in mRNA and their relationship with gene regulation (Kumar and Mohapatra, 2021). The frequent occurrence of RNA modifications indicate a critical role for cellular life and thus a need to understand the epitranscriptome landscape.

In attempt to characterize the epitranscriptome of the minimal bacteria, we primarily focused on identifying any potential changes between Syn1.0 and Syn3A which would be result from the genome reduction. **Table** S10 lists the identified enzymes within Syn1.0 (Hutchison et al., 2016) and Syn3A (Breuer et al., 2019; Thornburg et al., 2022) suspected to have methyltransferase ability. A majority (14/26) of the methyltransferase genes were removed during the genome reduction, so we hypothesize there should be some deviation between the epitranscriptome of Syn1.0 and Syn3A. Given the large variety of possible modifications such as $N^6$-methyladenosine ($m^6$A), 5-methylcytosine($m^5$C), and psuedouridine($\psi$), we focus on a single type of modification, $m^6$A. The $m^6$A modification and its location among the RNA nucleotide sequence has been confirmed to occur in the tRNA of *Mycoplasma* (Samuelsson et al., 1987; de Crécy-Lagard and Jaroch, 2021). Alanine, Serine, and Valine tRNA are known to undergo this modification at the $37^{th}$ base (canonical numbering) (Samuelsson et al., 1987). Additionally, a hypothetical enzyme, gene MMSYN1_0043, responsible for this function was proposed in de Crécy-Lagard and Jaroch (2021),Table S2. Computational methods coupled with experimental data are commonly utilized to detect RNA modications (Furlan et al., 2021). Therefore, we leveraged the ONT RNA sequencing data generated in this study coupled with a computational RNA modification detection algorithm to characterize the $m^6$A modifications within the minimal bacteria.

ONT direct RNA sequencing enables RNA modification detection because there is no cDNA conversion, which would remove the modifications. The RNA measured in ONT is the direct transcripts generated in the cell. A pivotal step in the ONT method is the threading of transcripts through the nanopore and then the conversion of an ionic current into a nucleotide sequence, base-calling. Base-calling only assigns canonical bases irrespective whether a base is modified or not. Computational methods have been developed which take advantage of the systematic error in the base-calling process effectively improving the its resolution, thus enabling the detection of modified bases (Furlan et al., 2021; Liu et al., 2019, 2021). The detection algorithm chosen for this work was the EpiNano reported in Liu et al. (2019, 2021). EpiNano algorithm is a supervised machine learning algorithm which can build models for $m^6$A detection. The original models were built by training with two datasets from yeast: RNA with known modifications and RNA with no modifications. The final results of the EpiNano algorithm for each base in the genome are a probability modification and a binary modified or unmodified. Using the original data sets reported in Liu et al. (2019) and the EpiNano pipeline, we re-built the models accounting for an updated base-caller (Guppy v6.1.3). The resulting detection model was then applied to the ONT data from JCVI-syn1.0 and JCVI-syn3A to corroborate the expected modifications sites in the tRNA genes. Additionally, we applied the newly trained model to some mRNA to see if difference between the organisms was detected. The results of the tRNA and mRNA are reported in **Fig.** S32 and **Fig.** S33, respectively. **Fig.** S32. The tRNA data (**Fig.** S32) generally shows no modifications detected for the known sites reported by Samuelsson et al. (1987); de Crécy-Lagard and Jaroch (2021). Only Syn1.0 rep1 data shows some corroborating detection in Valine (MMSYN1_0679) and Alanine (MMSYN1_0719), however there are a significant amounts of additional detected sites (orange markers). The mRNA data (**Fig.** S33) to compare organisms is not informative for identifying differences in the epitranscriptome. Syn1.0 rep1 and the Syn3Adata varies, however the Syn1.0 rep2 and Syn3Adata is essentially identical. The large disagreement between the Syn1.0 rep1 and rep2 data is possibly because of the base-caller methods used (see Materials and Methods). These comparison trends are also observed in tRNA data. In general the results are inconclusive in determining differences in the $m^6$A modifications between the two organisms. Most importantly the lack of detection at known literature sites is alarming and motivates a more comprehensive study in which the detection model is trained using *Mycoplasma* modified and unmodified data sets similar to the original usage in Liu et al. (2019). In doing so this could also be expanded to other modification types with the right complementary data sets.

### S4. Additional Supplementary Tables

**Table S1. Transcription factors of JCVI-syn1.0 and JCVI-syn3A**: Identified genes encoding transcription factors as determined by gene functional classifications according to the organisms GenBank files and Breuer et al. (2019). If the gene is found within Syn3A, then the gene classification is taken from Breuer et al. (2019). See footnotes for additional information about the transcription factors and their potential function in JCVI-syn1.0 and JCVI-syn3A assumed from the observed function in the model Gram-positive *B. subtilis*.

| Locus | Gene Name | NCBI Description | In Syn3A |
|---|---|---|---|
| 0042 | | Uncharacterized transcriptional regulator | Yes |
| 0020 | | transcriptional regulator, RpiR family | No |
| 0107 | nusB[a] | Transcription antitermination factor | Yes |
| 0187 | | transcriptional regulator, GntR family | No |
| 0211 | | transcription regulator GntR family | No |
| 0300 | nusA[b] | Transcription termination/antitermination protein NusA | Yes |
| 0350*** | hupA[c] | DNA-binding protein | Yes |
| 0407 | rpoD | RNA polymerase sigma factor | Yes |
| 0428 | phoU | phosphate transport system regulatory protein PhoU | Yes |
| 0430 | ylxM[d] | Putative (ylxM-like) effector of signal recognition particle | Yes |
| 0525 | mraZ[e] | Cell division/cell wall cluster transcriptional repressor | Yes |
| 0544 | hrcA[e] | Heat-inducible transcription repressor | Yes |
| 0620 | perR[e] | Uncharacterized transcriptional regulator | Yes |
| 0676 | | transcription regulator, LacI family | No |
| 0745 | | transcriptional regulator of the fructose operon, DeoR family | No |
| 0817*** | whiA[f] | Uncharacterized DNA-binding protein | Yes |
| 0840 | nusG[b] | Antitermination protein | Yes |
| 0858 | | Predicted transcriptional regulator, contains HTH domain | No |

[a] Associated to transcription of the ribosomal operon TheUniprotConsortium (2020)
[b] Involved in expressome formation O'Reilly et al. (2020)
[c] Related to supercoiling stabilization Gilbert et al. (2021)
[d] Predicted annotation from Bianchi et al. (2022)
[e] Regulated target identified and found in JCVI-Syn3A
[f] Involved in cell division and chromosome segregation and no transcription factor functionality in organism *B. subtilus* TheUniprotConsortium (2020)

**Table S2. Non-Coding RNA found within Syn1.0 and Syn3A**: Locus tags are consistent between both organisms. Table does not included any asRNA.

| Locus | Gene Name | NCBI Description | ncRNA / Riboswitch |
|---|---|---|---|
| 0049 | ffs | Signal recognition particle sRNA small type | ncRNA |
| 0158 | ssrA | Transfer-messenger RNA | ncRNA |
| 0356 | rnpB | RNase P RNA component class B | ncRNA |
| intergenic[a] | n/a | SAM riboswitch class I | riboswitch |
| intergenic[b] | n/a | PP riboswitch (THI element) | riboswitch |
| intergenic[c] | n/a | T-Box Riboswitch I | riboswitch |
| intergenic[d] | n/a | T-Box Riboswitch II | riboswitch |

[a] Between gene 0431 and 0432 on forward strand, Assigned locus tag 0933 within JCVI-syn3A
[b] Between gene 0708 and 0710 on reverse strand, Assigned locus tag 0934 within JCVI-syn3A
[c] Between gene 0221 and 0222 on forward strand, identified using RiboD prediction scanner Mukherjee et al. (2019)
[d] Between gene 0519 and 0520 on reverse strand, identified using RiboD prediction scanner Mukherjee et al. (2019)

**Table S3. NCBI GenBank Entries:** For each organism in the study, the accession number and reference link is provided. All gene features and sequence information was obtained from the GenBank files. Specific version have been listed as GenBank files are constantly updated. GenBanks are available at `https://www.ncbi.nlm.nih.gov/nuccore/` Sayers et al. (2021)

| Organism | Accession # | Version | Reference |
|---|---|---|---|
| JCVI-syn3A | CP016816.2 | 12-APR-2018 | GenBank: Accession No. CP016816.2 (2018) |
| JCVI-syn1.0 | CP002027.1 | 29-SEP-2010 | GenBank: Accession No. CP002027.1 (2010) |
| *Mycoplasma mycoides* subsp. *capri* | CP001621.1 | 31-JAN-2014 | GenBank: Accession No. CP001621.1 (2014) |
| *Mesoplasma florum* | NC_006055.1 | 29-MAR-2022 | GenBank: Accession No. NC_006055.1 (2004) |
| *Mycoplasma pneumoniae* | NC_000912.1 | 15-FEB-2022 | GenBank: Accession No. NC_000912.1 (2001) |

**Table S4. Sequences for Motif Identification**: List of all sequences used in the bioinformatic identification of motifs within the genomes of Syn1.0 and related organisms. Protocol for motif identification is outlined in Materials and Methods–**Section** 4.1. Results are discussed in **Section** S2.

| Organism | anti-Shine-Dalgarno | Shine-Dalgarno | Promoter[1] |
|---|---|---|---|
| JCVI-syn1.0 | $5'$-*CCTCCTTTCT*-$3'$ | $5'$-*AGAAAGGAGG*-$3'$ | $5'$-*TANAAT*-$3'$ |
| JCVI-syn3A | $5'$-*CCTCCTTTCT*-$3'$ | $5'$-*AGAAAGGAGG*-$3'$ | $5'$-*TANAAT*-$3'$ |
| *Mycoplasma mycoides* subsp. *capri* | $5'$-*CCTCCTTTCT*-$3'$ | $5'$-*AGAAAGGAGG*-$3'$ | $5'$-*TANAAT*-$3'$ |
| *Mesoplasma florum* | $5'$-*ACCTCCTTT*-$3'$ | $5'$-*AAAGGAGGT*-$3'$ | $5'$-*TANAAT*-$3'$ |
| *Mycoplasma pneumoniae* | $5'$-*ACCTCCTTT*-$3'$ | $5'$-*AAAGGAGGT*-$3'$ | $5'$-*TANAAT*-$3'$ |

[1] An additional sequence was also used as template including the extended region of the promoter $5'$-*TGNTANAAT*-$3'$

**Table S5. RNA Sequencing Read Summary**: Summary of the RNA sequencing performed in the experiments. PacBio coverage values are likely inflated (factor of 2 greater) as the experiments in this study are not strand specific (see **Section** 4.5).

| Method | Total Reads | Mapped Reads | Avg Read Length (kbp) | Coverage | Organism |
|---|---|---|---|---|---|
| ONT | 234,477 | 211,631 | 0.580 | 58x | Syn1.0 |
| ONT | 584,958 | 472,818 | 0.348 | 64x | Syn1.0 |
| ONT | 734,075 | 586,484 | 0.395 | 158x | Syn3A |
| Illumina | 2,171,606 | 2,168,938 | 0.201 | 201x | Syn1.0 |
| Illumina | 1,021,820 | 1,019,507 | 0.192 | 91x | Syn1.0 |
| Illumina | 1,009,782 | 1,006,549 | 0.195 | 90x | Syn1.0 |
| PacBio | 930,830 | 930,814 | 1.82 | 755x | Syn1.0 |
| PacBio | 962,582 | 962,519 | 1.83 | 784x | Syn1.0 |
| PacBio | 1,056,477 | 1,055,803 | 1.79 | 840x | Syn1.0 |

**Table S6. Cellular RNA Concentrations**: JCVI-syn1.0 RNA concentrations categorized by RNA types for each RNAseq method. Concentrations were calculated according to absolute counts assuming spherical cells with a diameter of 493 nm Moger-Reischer et al. (2023). See **Section** 4.7 for explanation of absolute abundances determination. Additional values obtained from *Mesoplasma florum* Matteau et al. (2020) (Table EV1) are given for comparison. (ncRNA were not reported for *M. florum*). Values given in units of RNA/$\mu m^3$.

| Sample | mRNA | ncRNA | rRNA | tRNA |
|---|---|---|---|---|
| ONT Rep 1 | $6.3 \times 10^3$ | $2.3 \times 10^3$ | $3.1 \times 10^4$ | $1.7 \times 10^5$ |
| ONT Rep 2 | $4.7 \times 10^3$ | $2.3 \times 10^3$ | $1.5 \times 10^3$ | $1.7 \times 10^5$ |
| Illumina Rep 1 | $6.5 \times 10^3$ | $3.8 \times 10^3$ | $4.3 \times 10^3$ | $1.7 \times 10^5$ |
| Illumina Rep 2 | $8.6 \times 10^3$ | $3.5 \times 10^3$ | $1.1 \times 10^4$ | $1.7 \times 10^5$ |
| Illumina Rep 3 | $9.5 \times 10^3$ | $4.2 \times 10^3$ | $1.3 \times 10^3$ | $1.7 \times 10^5$ |
| PacBio Rep 1 | $5.3 \times 10^3$ | $4.8 \times 10^3$ | $6.6 \times 10^3$ | $1.7 \times 10^5$ |
| PacBio Rep 2 | $5.4 \times 10^3$ | $4.5 \times 10^3$ | $1.8 \times 10^3$ | $1.7 \times 10^5$ |
| PacBio Rep 3 | $5.3 \times 10^3$ | $3.7 \times 10^3$ | $2.5 \times 10^3$ | $1.7 \times 10^5$ |
| Avg Syn1.0 | $6.5 \times 10^3$ | $3.6 \times 10^3$ | $7.5 \times 10^3$ | $1.7 \times 10^5$ |
| Avg Syn3A | $5.8 \times 10^3$ | n.r. | n.r. | n.r. |
| *M. florum* | $4.7 \times 10^3$ | n.r. | $5.4 \times 10^4$ | $2.0 \times 10^5$ |

**Table S7. Cellular RNA Abundances**: JCVI-syn1.0 RNA abundances categorized by RNA types for each RNAseq method. Concentrations were calculated according to absolute counts assuming spherical cells with a diameter of 493 nm Moger-Reischer et al. (2023). See **Section** 4.7 for explanation of absolute abundance determination. Additional values obtained from *Mesoplasma florum* Matteau et al. (2020) (Table EV1, Dataset EV8) are given for comparison. (ncRNA were not reported for *M. florum*).Syn3A data is obtained from Sandberg et al. (2023), and then converted to absolute abundances following **Section** 4.7 with Syn3A relevant parameters.

| Sample | mRNA | | ncRNA | | rRNA | | tRNA | |
|---|---|---|---|---|---|---|---|---|
| | Total | Avg | Total | Avg | Total | Avg | Total | Avg |
| ONT Rep 1 | 280 | 0.3±1.1 | 104 | 20.7±25.9 | 1388 | 231.4±463.6 | 7666 | 255.5±590.8 |
| ONT Rep 2 | 210 | 0.2±0.8 | 101 | 20.2±31.9 | 65 | 10.8±10.4 | 7629 | 254.3±478.0 |
| Illumina Rep 1 | 290 | 0.3±3.1 | 168 | 33.7±22.7 | 192 | 32.0±42.8 | 7723 | 257.4±423.1 |
| Illumina Rep 2 | 383 | 0.4±6.1 | 154 | 30.8±20.4 | 506 | 84.4±166.5 | 7651 | 255.0±370.3 |
| Illumina Rep 3 | 420 | 0.5±7.4 | 187 | 37.3±37.4 | 57 | 9.5±21.3 | 7645 | 254.8±486.3 |
| PacBio Rep 1 | 236 | 0.3±1.1 | 215 | 42.9±22.3 | 291 | 48.6±79.9 | 7694 | 256.5±660.4 |
| PacBio Rep 2 | 238 | 0.3±0.9 | 197 | 39.5±12.9 | 82 | 13.6±10.4 | 7695 | 256.5±680.7 |
| PacBio Rep 3 | 236 | 0.3±0.8 | 165 | 33.0±11.9 | 109 | 18.2±16.8 | 7700 | 256.7±690.0 |
| Avg Syn1.0 | 236 | 0.3±0.8 | 165 | 33.0±11.9 | 109 | 18.2±16.8 | 7700 | 256.7±690.0 |
| Avg Syn3A | 196 | 0.4±1.0 | n.r | n.r | n.r | n.r | n.r | n.r |
| *M. florum* | 420 | 0.6±0.9 | n.r | n.r | 4900 | 809.6±0.0 | 18000 | 625.7±0.0 |

**Table S8. Syn1.0 Experimental Media Components**: Detailed list of the SP4+KnockOut media components used to grow the Syn1.0 cells. Preparation instructions are provided in **Section** 4.4.1.

| Media Component | Vendor | Serial # |
|---|---|---|
| Mycoplasma Broth Base | BD | DF0554-17-1 |
| Bacto Tryptone | BD | 211705 |
| Bacto Peptone | BD | 211677 |
| Glucose 20% w/v stock | Thermo Fisher Scientific | 34273900 |
| CMRL 1066 (10X stock w/o phenol red; w/o bicarb; w/o Gln) | Thermo Fisher Scientific | 21-540-026 |
| Sodium bicarbonate 7.5% w/v stock | Thermo Fisher Scientific | 32100113 |
| L-glutamine 200 mM stock | Thermo Fisher Scientific | 25030081 |
| Yeast extract solution | Thermo Fisher Scientific | 18180059 |
| TC Yeastolate 2% w/v stock, autoclaved | Gibco | 255772 |
| Serum (heat inactivated FBS, HS) OR substitute (KO) | Thermo Fisher Scientific | 10828028 |
| Penicillin G (400,000 U/mL stock) | Sigma-Aldrich | P3032-1MU |
| Phenol red (0.5% w/v filter sterilized) | Sigma-Aldrich | P0290-100ML |

**Table S9. Parameters for Quantification of Cellular Components**: Values were used for the calculation of the dry weight calculation of Syn1.0, see **Section** 4.7

| Parameter | Value | Unit | Reference |
|---|---|---|---|
| Water Volume per Protein Mass | 4.8 | $\mu L/mg$ | Leblanc and Le Grimellec (1979)[a] |
| Cellular Density ($\rho_{cell}$) | 1.1 | $g/mL$ | Bratbak and Dundas (1984)[c] |
| Syn1.0 Protein Dry Mass Fraction | 58.2 | % | Razin et al. (1963)[b] |
| Syn1.0 RNA Dry Mass Fraction | 17.3 | % | Razin et al. (1963)[b] |
| Syn1.0 Diameter | 439 | nm | Moger-Reischer et al. (2023) |
| Syn3A Protein Dry Mass Fraction | 54.727 | % | Breuer et al. (2019) |
| Syn3A RNA Dry Mass Fraction | 16.274 | % | Breuer et al. (2019) |
| Syn3A Diameter | 400 | nm | Breuer et al. (2019) |
| Syn1.0 Total Cellular Dry Mass | 12.8 | fg | *This Study* |
| Syn3A Total Cellular Dry Mass | 10.2 | fg | Breuer et al. (2019) |
| *M. florum* Total Cellular Dry Mass | 22.1 | fg | Matteau et al. (2020) |

[a] from *M. mycoides capri* serovar capri PG3
[b] from *M. mycoides* var. *capri*
[c] from *B. subtilis*, *E. coli*, and *P. putida*

**Table S10. JCVI-syn1.0 Methyltransferases:** Genes within Syn1.0 suspected to have methyltransferase activity. NCBI descriptions of the genes are pulled from the Syn1.0 GenBank EntryGenBank: Accession No. CP002027.1 (2010). Genes retained in Syn3A as a result of the genome reduction are designated. Gene MMSYN1_0043 has been identified to encode for the protein responsible for $m^6A$ modifications de Crécy-Lagard and Jaroch (2021)(SI Table 2).

| Locus Tag | NCBI Gene Product | In Syn3A |
|---|---|---|
| 0004 | dimethyladenosine transferase | Yes |
| 0043 | methyltransferase | Yes |
| 0056 | GATC–recognizing Type II restriction modification system (MmyCV) adenine DNA methyltransferase subunit | No |
| 0098 | CCATC–recognizing Type II restriction modification system (MmyCVI) adenine DNA methyltransferase subunit 1 | No |
| 0099 | CCATC–recognizing Type II restriction modification system (MmyCVI) adenine DNA methyltransferase subunit 2 | No |
| 0104 | ribosomal RNA large subunit methyltransferase J | No |
| 0142 | protein-(glutamine-N5) methyltransferase, release factor-specific | Yes |
| 0156 | tRNA (guanine-N(7)-)-methyltransferase | No |
| 0202 | methyltransferase | Yes |
| 0204 | ribosomal RNA small subunit methyltransferase B | No |
| 0313 | methylenetetrahydrofolate–tRNA-(uracil-5-)-methyltransferase trmFO 2 | No |
| 0364 | tRNA (guanine-N1)-methyltransferase | Yes |
| 0387 | tRNA (5-methylaminomethyl-2-thiouridylate)-methyltransferase | Yes |
| 0434 | tRNA:M(5)U-54 methyltransferase | Yes |
| 0448 | rRNA methylase | Yes |
| 0524 | S-adenosyl-methyltransferase MraW | Yes |
| 0548 | SAM-dependent tRNA methylase related to TrmL | Yes |
| 0554 | C-5 cytosine-specific DNA methylase | No |
| 0591 | Type III restriction-modification system (MmyCI) adenine DNA methyltransferase subunit[a] | No |
| 0705 | C-5 cytosine-specific DNA methylase | No |
| 0742 | CCTTC-recognizing Type II restriction modification system (MmyCII) adenine/cytosine DNA methyltransferase subunit | No |
| 0754 | GCATC–recognizing Type II restriction modification system (MmyCIII) adenine DNA methyltransferase subunit 1[a] | No |
| 0755 | GCATC–recognizing Type II restriction modification system (MmyCIII) adenine DNA methyltransferase subunit 2 | No |
| 0769 | GANTC–recognizing Type II restriction modification system (MmyCIV) adenine DNA methylase subunit | No |
| 0799 | glycine hydroxymethyltransferase | Yes |
| 0838 | RNA methyltransferase, TrmH family, group 3 | Yes |
| 0874 | 16S rRNA methyltransferase GidB | Yes |

[a] Identified as a pseudogene in JCVI-syn1.0 GenBank GenBank: Accession No. CP002027.1 (2010)

### S5. Additional Supplementary Figures

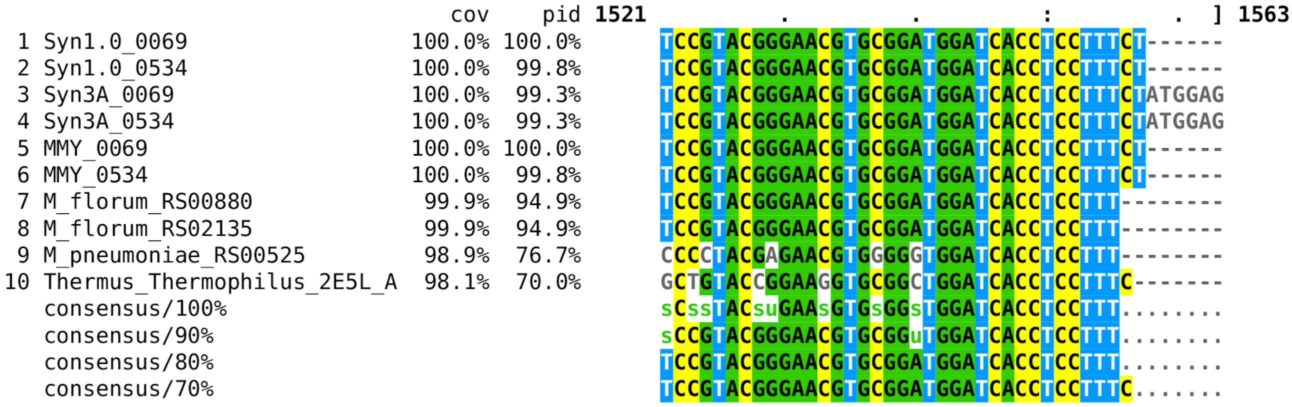


**Figure S1. Sequence Alignment of 16S rRNA**: Sequence alignment of the $3'$end of 16S rRNAs for the organisms used in this study. Some organisms have multiple entries because they contain duplicates of 16S rRNA encoding genes. Locus tag information has been included for each entry. MMY refers to *M. mycoides* subsp. *capri*. The 16S rRNA of Thermus Thermophilus was included as the SD-aSD structure observed motivated the identification approach used Kaminishi et al. (2007). Its sequence was obtained from the PDB entry (2E5L) published in Kaminishi et al. (2007), so no locus tag is information is included. Alignments were performed using CLUSTAL OMEGA Sievers et al. (2011) via EMBL-EBI Services Harte et al. (2004); Madeira et al. (2022). Results are visualized using MView Brown et al. (1998).

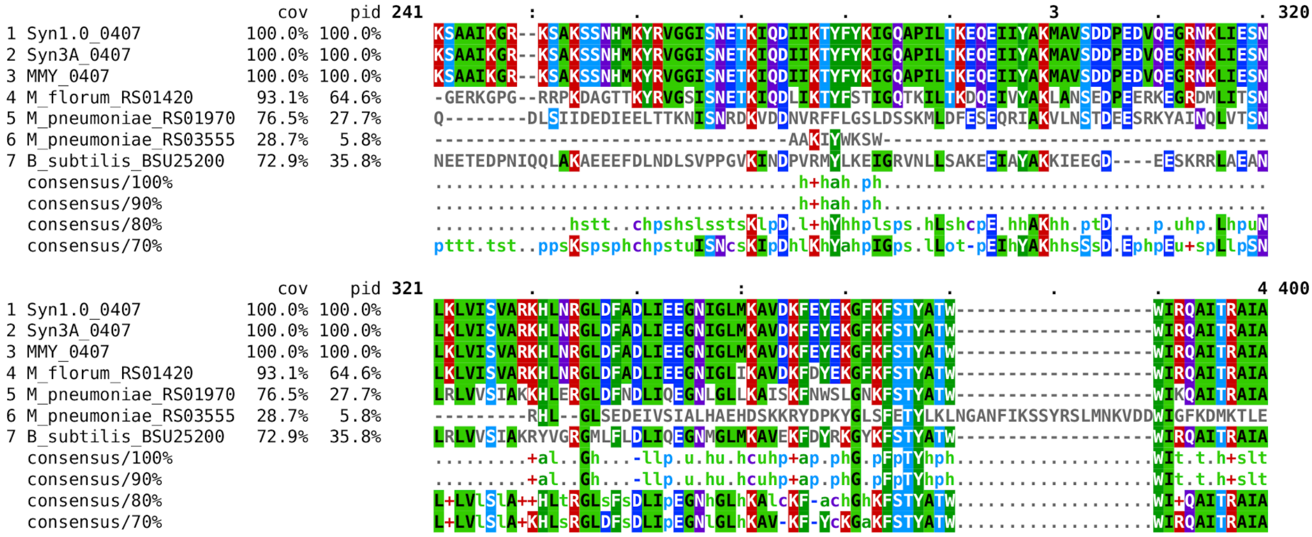


**Figure S2. Sequence Alignment of RNAP sigma70 factor**: Sequence alignment of domain 2 and domain 3 regions of the RNAP sigma70 factor are shown for Syn1.0 and related organisms. These domains are observed to make the physical contacts with the promoter motif on the DNA Chen et al. (2020). General agreement in the sequence alignment allows for the usage of the well characterized promoter consensus sequence, $5'$-*TANAAT*-$3'$. Alignments were performed using CLUSTAL OMEGA Sievers et al. (2011) via te EMBL-EBI Services Harte et al. (2004); Madeira et al. (2022). Results are visualized using MView Brown et al. (1998).

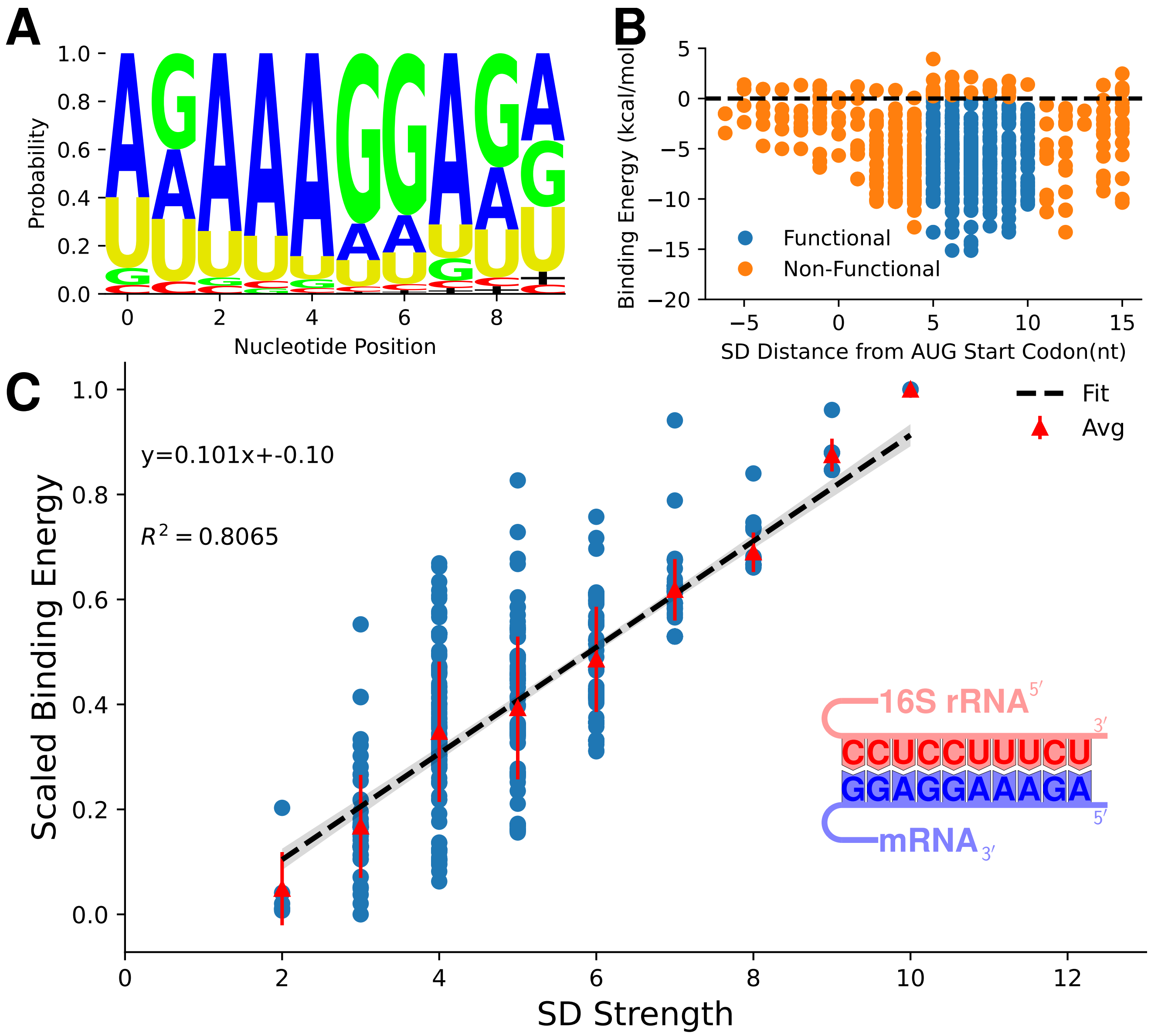


**Figure S3. Shine-Dalgarno (SD) results for JCVI-syn1.0**: **(A)** Nucleotide probability for the SD sequences for each gene. As expected the observed sequences matches closely with the SD template sequence, $5'$-*AGAAAGGAGG*-$3'$. Black plus signs indicate where no nucleotides is assigned due to motif sequence being found near the boundary of the search region. **(B)** Breakdown of functionality within the identified SD for all genes. Functionality is assigned by the SD distance from the AUG start codon (between 5 and 10 nucleotides) and whether the calculated binding energy between the SD-aSD is non-positive. (the black dotted line shows the boundary line, $\Delta G \leq 0$). **(C)** Relationship of the SD strength (sequence alignment score) to the binding energy between the SD and aSD. Data is shown for the genes with functional SD only. Related to **Fig.** S8A.

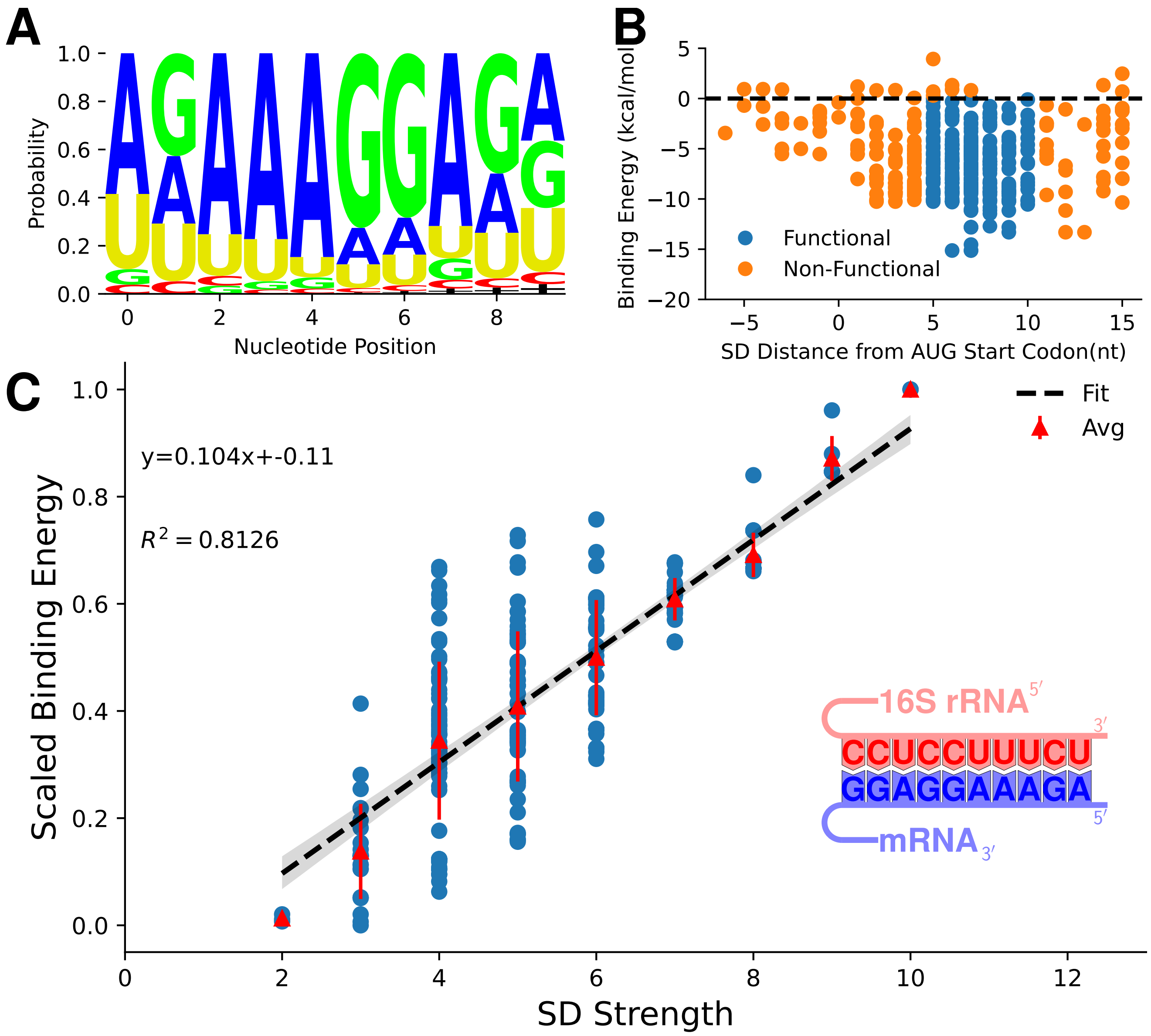


**Figure S4. Shine-Dalgarno results for Syn3A**: (A) Nucleotide probability for the SD sequences for each gene. As expected the observed sequences matches closely with the SD template sequence, $5'$-*AGAAAGGAGG*-$3'$. Black plus signs indicate where no nucleotides is assigned due to motif sequence being found near the boundary of the search region. (B) Breakdown of functionality within the identified SD for all genes. Functionality is assigned by the SD distance from the AUG start codon (between 5 and 10 nucleotides) and whether the calculated binding energy between the SD-aSD is non-positive. (the black dotted line shows the boundary line, $\Delta G <= 0$). (C) Relationship of the SD strength (sequence alignment score) to the binding energy between the SD and aSD. Data is shown for the genes with functional SD only. Validation and relationship to proteomics shown in **Fig.** S8B.

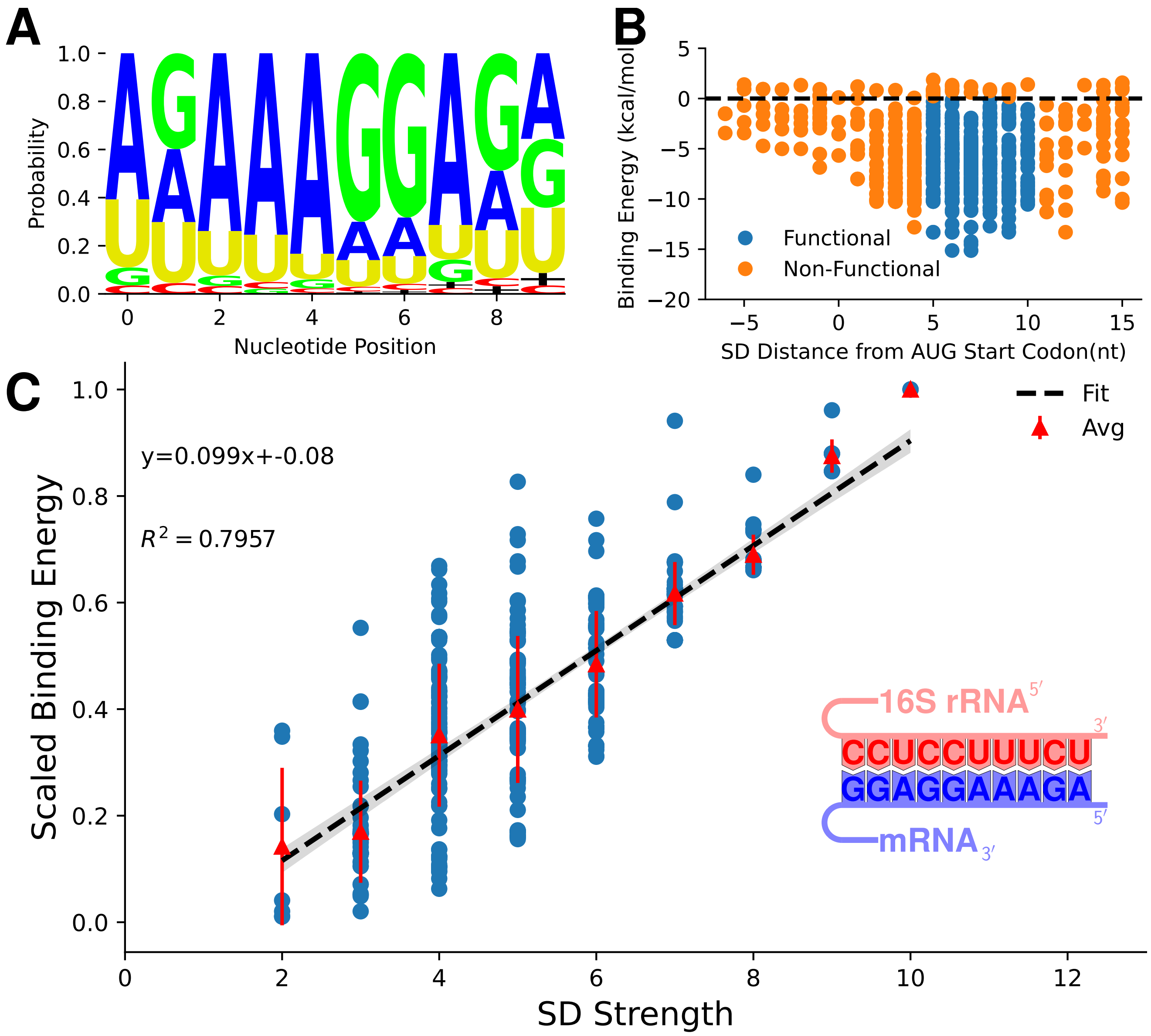


**Figure S5. Shine-Dalgarno results for** *M. florum*: (A) Nucleotide probability for the SD sequences for each gene. As expected the observed sequences matches closely with the SD template sequence, $5'$-*AGAAAGGAGG*-$3'$. Black plus signs indicate where no nucleotides is assigned due to motif sequence being found near the boundary of the search region. (B) Breakdown of functionality within the identified SD for all genes. Functionality is assigned by the SD distance from the AUG start codon (between 5 and 10 nucleotides) and whether the calculated binding energy between the SD-aSD is non-positive. (the black dotted line shows the boundary line, $\Delta G <= 0$). (C) Relationship of the SD strength (sequence alignment score) to the binding energy between the SD and aSD. Data is shown for the genes with functional SD only. Validation and relationship to proteomics shown in **Fig.** S8C.

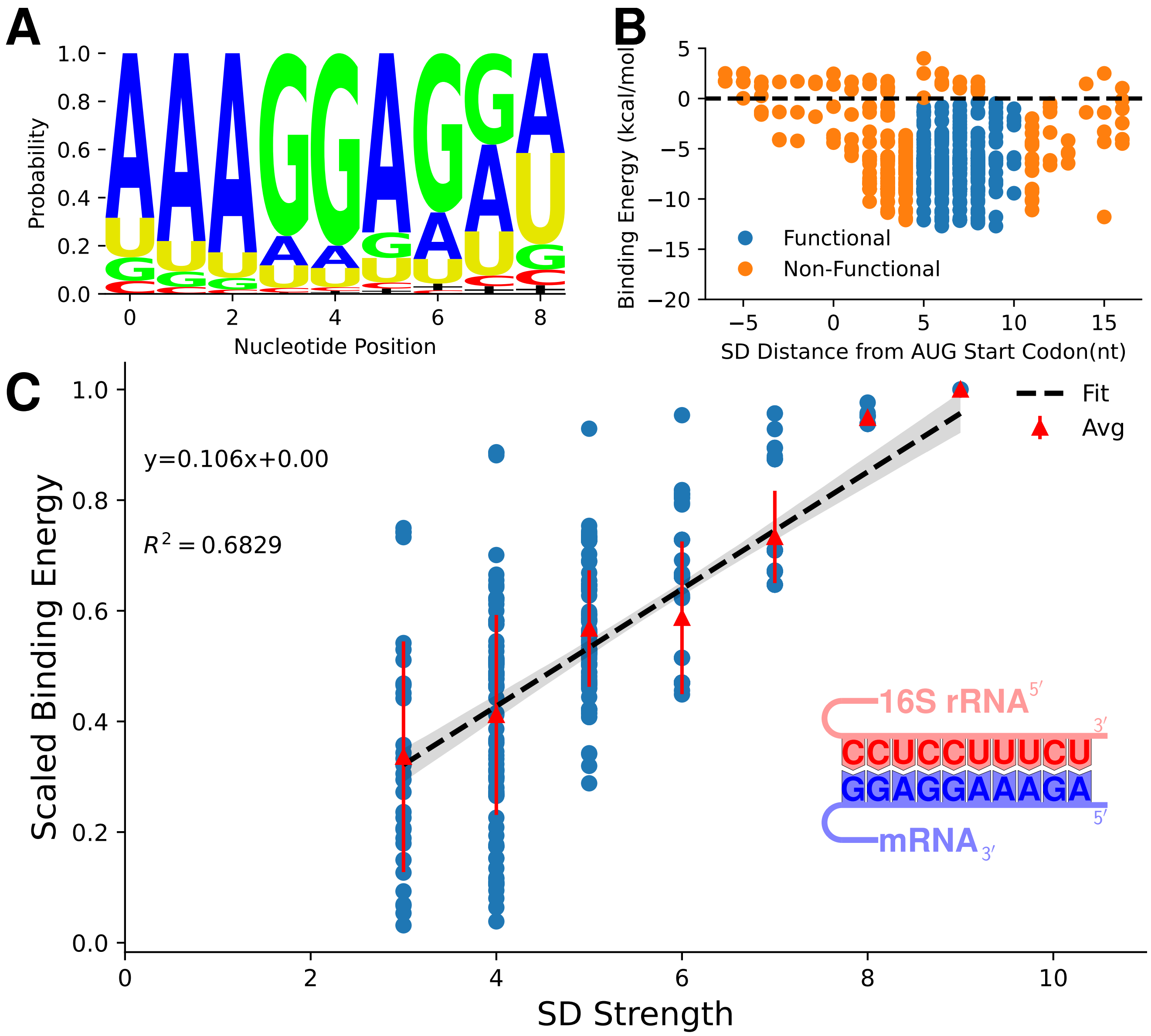


**Figure S6. Shine-Dalgarno results for** *M. mycoides* **subsp.** *capri*. (A) Nucleotide probability for the SD sequences for each gene. As expected the observed sequences matches closely with the SD template sequence, $5'$-*AGAAAGGAGG*-$3'$. Black plus signs indicate where no nucleotides is assigned due to motif sequence being found near the boundary of the search region. (B) Breakdown of functionality within the identified SD for all genes. Functionality is assigned by the SD distance from the AUG start codon (between 5 and 10 nucleotides) and whether the calculated binding energy between the SD-aSD is non-positive. (the black dotted line shows the boundary line, $\Delta G <= 0$). (C) Relationship of the SD strength (sequence alignment score) to the binding energy between the SD and aSD. Data is shown for the genes with functional SD only. Validation shown in **Fig.** S8D

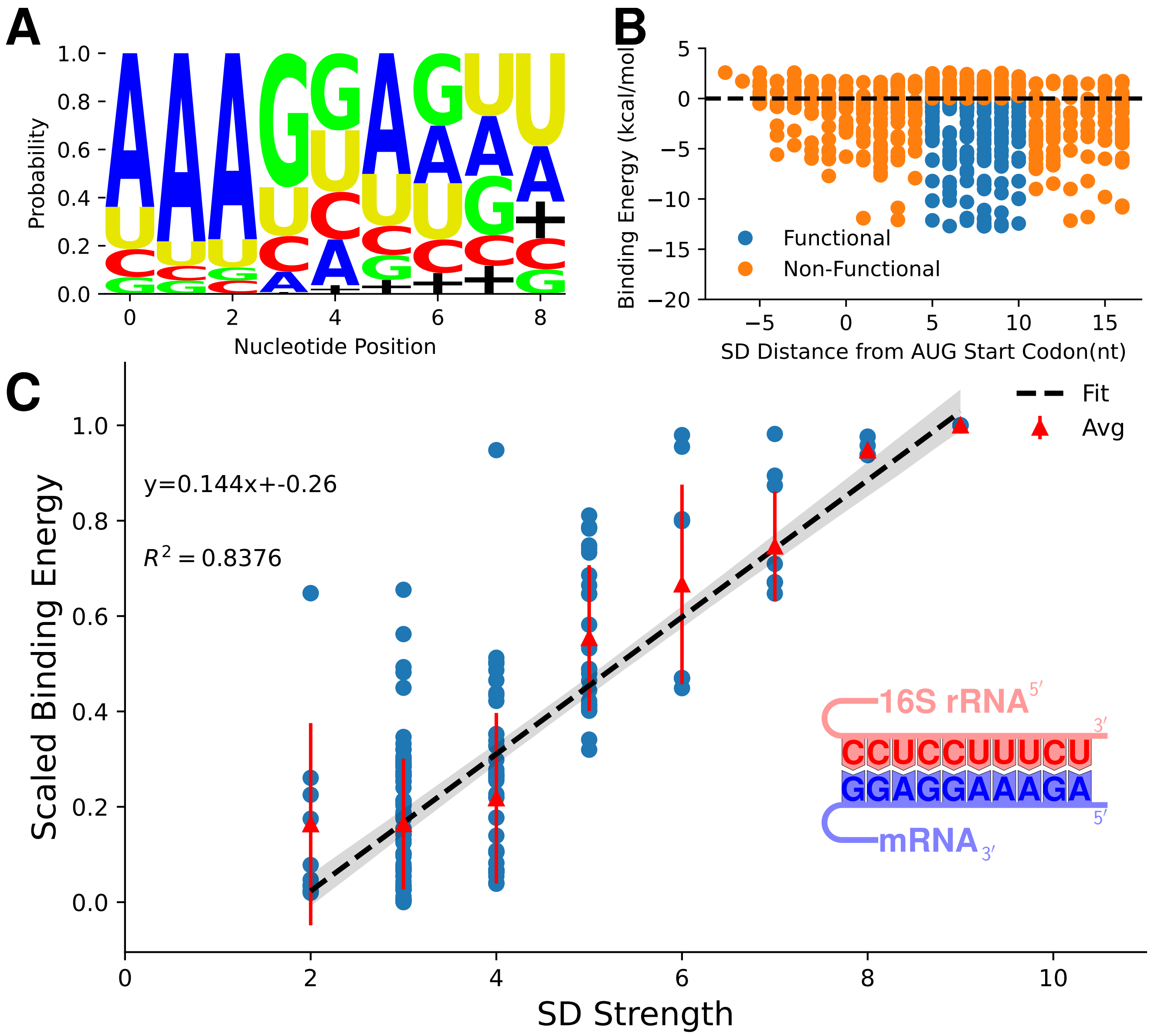


**Figure S7. Shine-Dalgarno results for** ***M. pneumoniae***: (A) Nucleotide probability for the SD sequences for each gene. As expected the observed sequences matches closely with the SD template sequence, $5'$-*AGAAAGGAGG*-$3'$. Black plus signs indicate where no nucleotides is assigned due to motif sequence being found near the boundary of the search region. (B) Breakdown of functionality within the identified SD for all genes. Functionality is assigned by the SD distance from the AUG start codon (between 5 and 10 nucleotides) and whether the calculated binding energy between the SD-aSD is non-positive. (the black dotted line shows the boundary line, $\Delta G <= 0$). (C) Relationship of the SD strength (sequence alignment score) to the binding energy between the SD and aSD. Data is shown for the genes with functional SD only. Validation shown in **Fig.** S8E.

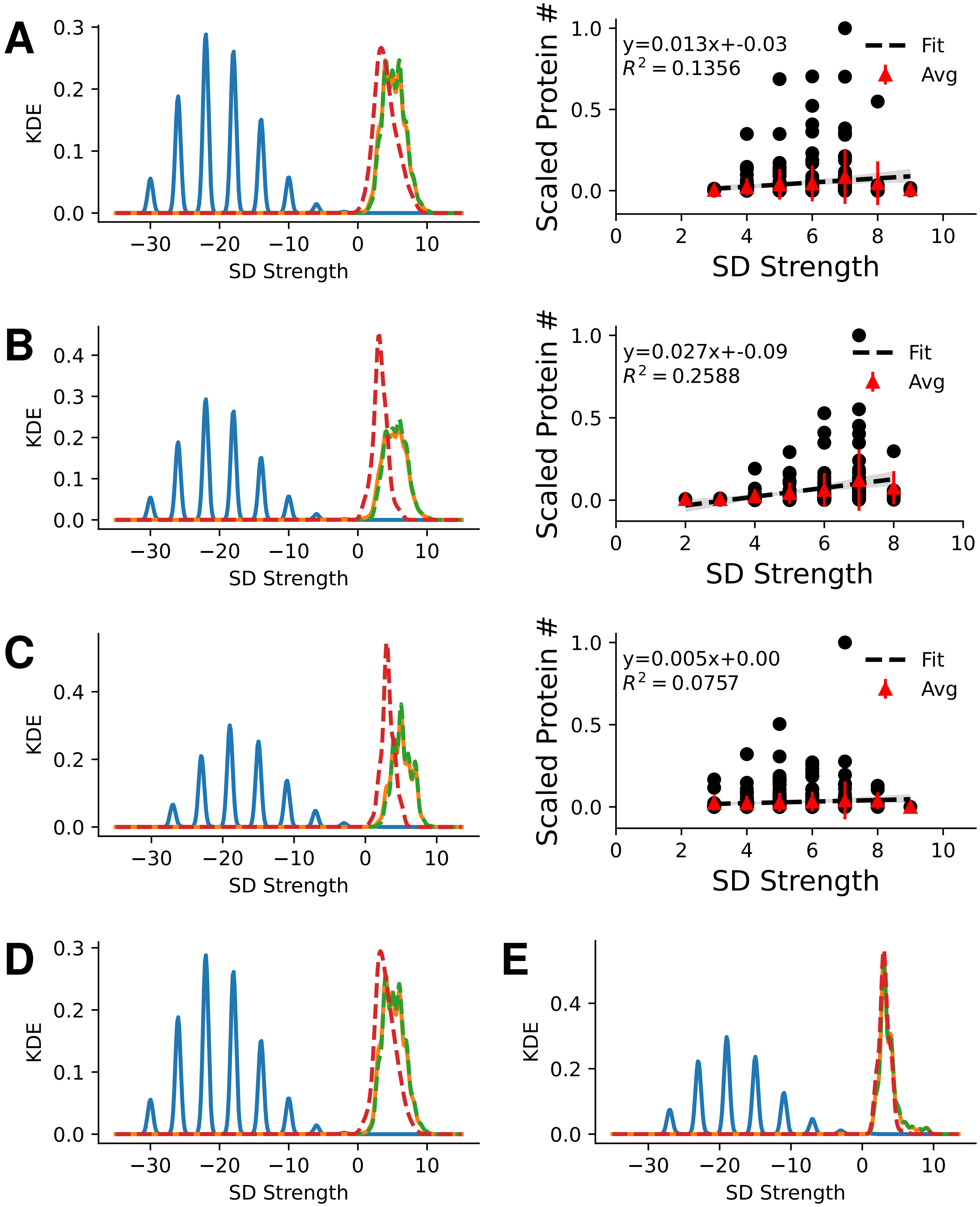


**Figure S8. Validation of Shine-Dalgarnosequences**: Results of the non-specific SD search within Syn1.0 **(A)** (left), Syn3A **(B)** (left), *M. florum* **(C)** (left), *M. mycoides* subsp. *capri* **(D)**, and *M. pneumoniae* **(E)**. Non-specific SD strengths (blue) show much lower values than the targeted search for SD upstream of genes (orange). Protein coding (green) and non-coding genes (red) are separated to highlight slightly higher average SD strengths found within protein coding (translated) genes. When proteomics data was available the linear relationship between SD strength and protein abundances was measured for functional SD only. Results show weak correlation ($R^2 < 0.2$) for Syn1.0, Syn3A, and *M. florum*,(**A, B, and C**, right) respectively. Proteomic data was obtained from this study (Syn1.0), Breuer et al. (2019) (Syn3A), and Matteau et al. (2020) (*M. florum*). Related to **Figs.** S3-S7

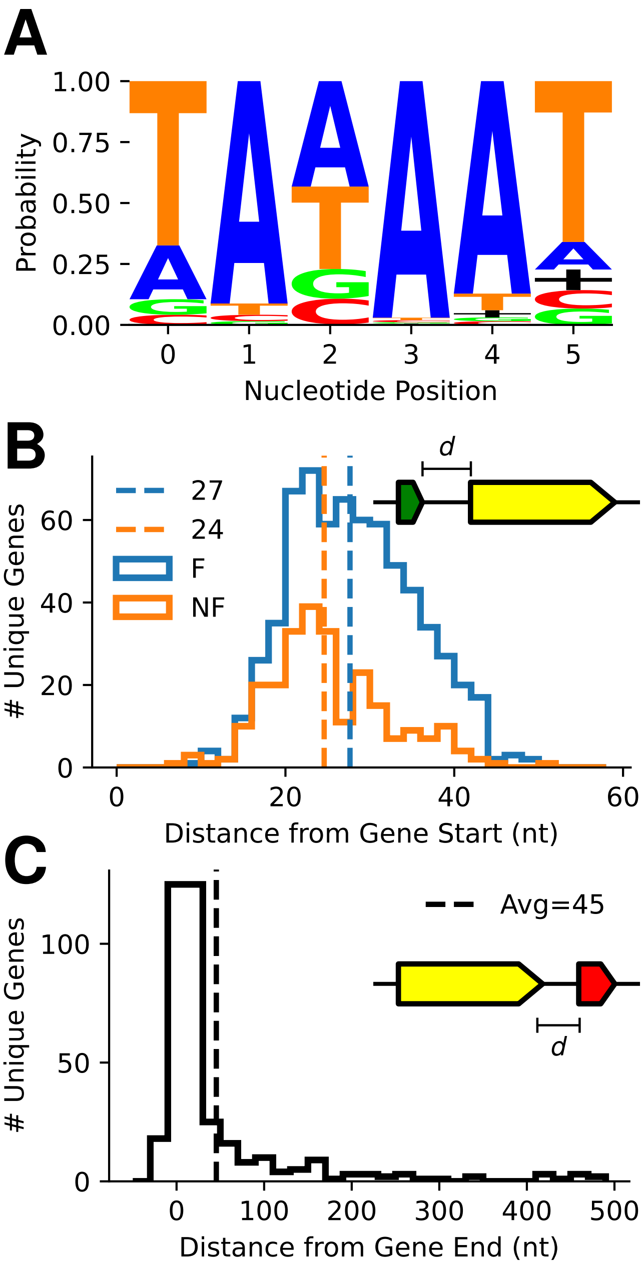


**Figure S9. Promoter and Termination results for JCVI-syn1.0**: **(A)** Nucleotide probability for the SD sequences for each gene. As expected the observed sequences matches closely with the promoter template sequence, $5'$-*TANAAT*-$3'$. Black plus signs indicate where no nucleotides were assigned due to motif sequences being found near the boundary of the search region. **(B)** Distance from gene start to the identified promoter sequences for functional (blue) and non-functional (orange) motifs. Results are in agreement with data from *M. florum* Matteau et al. (2020). Functionality is determined using the scores and a comparison to non-target search of the genome. (See **Section** 4.1.3). **(C)** Distance in nucleotide from the gene end to the start of the intrinsic termination loop. Values agree with reported numbers for *M. mycoides* subsp. *capri* in de Hoon et al. (2005). Related to **Fig.** S14A.

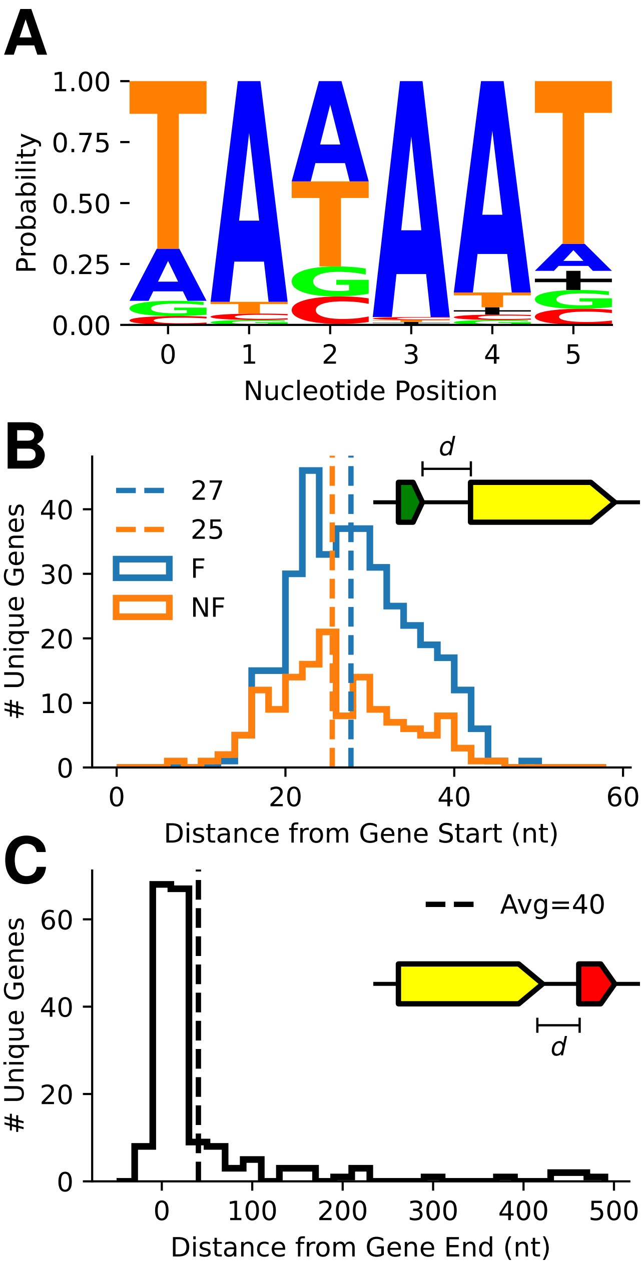


**Figure S10. Promoter and Termination results for JCVI-syn3A (A)** Nucleotide probability for the SD sequences for each gene. As expected the observed sequences matches closely with the promoter template sequence, $5'$-*TANAAT*-$3'$. Black plus signs indicate where no nucleotides is assigned due to motif sequence being found near the boundary of the search region. **(B)** Distance from gene start to the identified promoter sequences for functional (blue) and non-functional (orange) motifs. Results are in agreement with data from *M. florum* Matteau et al. (2020). Functionality is determined using the scores and a comparison to non-target search of the genome. (See **Section** 4.1.3). **(C)** Distance in nucleotide from the gene end to the start of the intrinsic termination loop. Values agree with reported numbers for *M. mycoides* subsp. *capri* in de Hoon et al. (2005). Validation and relationship to proteomics shown in Figure **Fig.** S14B.

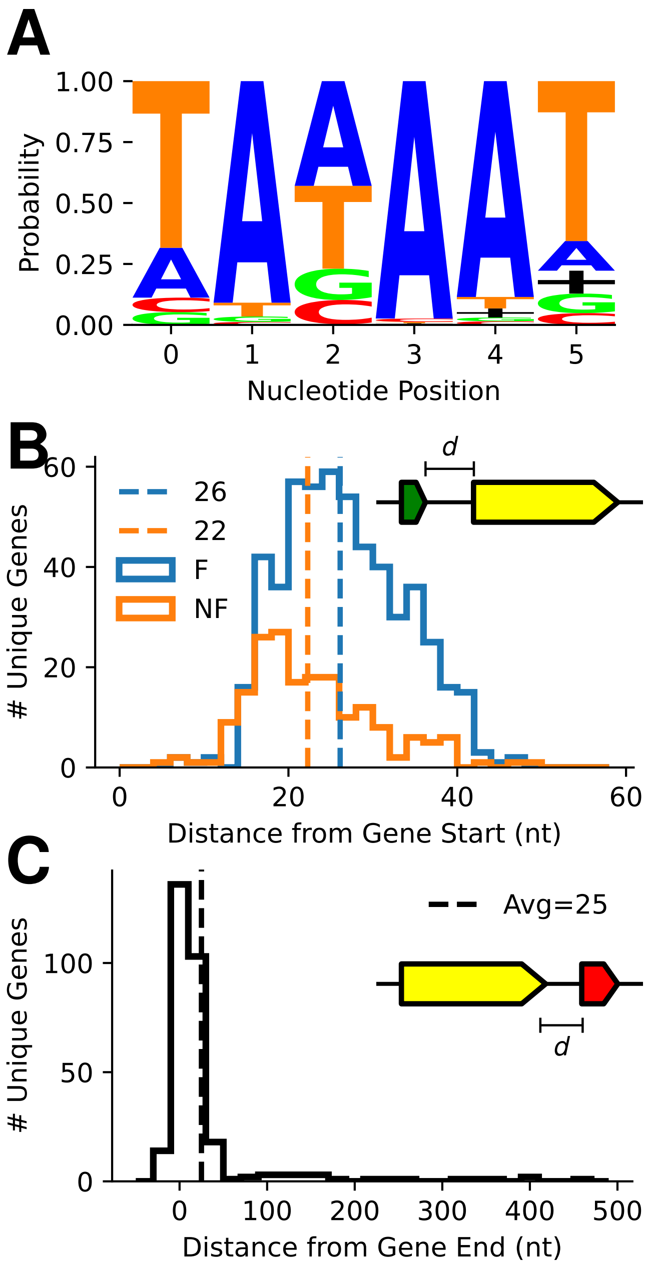


**Figure S11. Promoter and Termination results for** *M. florum*: **(A)** Nucleotide probability for the SD sequences for each gene. As expected the observed sequences matches closely with the promoter template sequence, $5'$-*TANAAT*-$3'$. Black plus signs indicate where no nucleotides is assigned due to motif sequence being found near the boundary of the search region. **(B)** Distance from gene start to the identified promoter sequences for functional (blue) and non-functional (orange) motifs. Results are in agreement with data from *M. florum* Matteau et al. (2020). Functionality is determined using the scores and a comparison to non-target search of the genome. (See **Section** 4.1.3). **(C)** Distance in nucleotide from the gene end to the start of the intrinsic termination loop. Values agree with reported numbers for *M. florum* in de Hoon et al. (2005). Validation and relationship to proteomics shown in **Fig.** S14C.

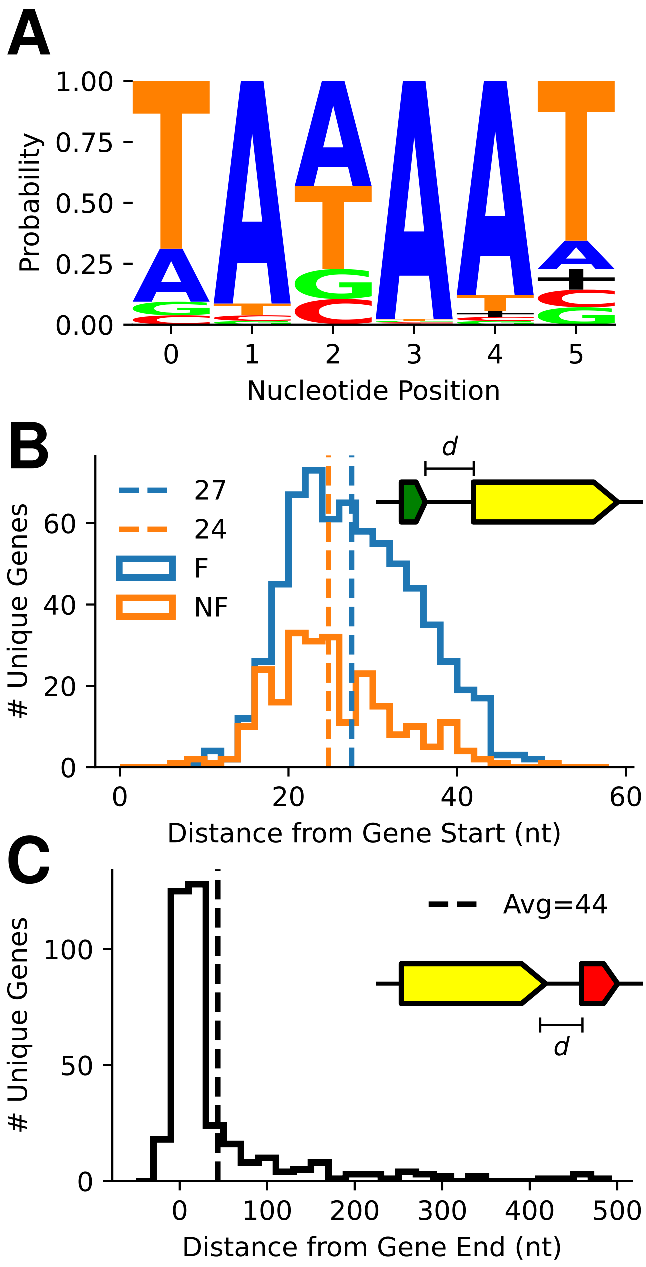


**Figure S12. Promoter and Termination results for** *M. mycoides* **subsp.** *capri*: **(A)** Nucleotide probability for the SD sequences for each gene. As expected the observed sequences matches closely with the promoter template sequence, $5'$-*TANAAT*-$3'$. Black plus signs indicate where no nucleotides is assigned due to motif sequence being found near the boundary of the search region. **(B)** Distance from gene start to the identified promoter sequences for functional (blue) and non-functional (orange) motifs. Results are in agreement with data from *M. florum* Matteau et al. (2020). Functionality is determined using the scores and a comparison to non-target search of the genome. (See **Section** 4.1.3). **(C)** Distance in nucleotide from the gene end to the start of the intrinsic termination loop. Values agree with reported numbers for *M. mycoides* subsp. *capri* in de Hoon et al. (2005). Validation shown in **Fig.** S14D.

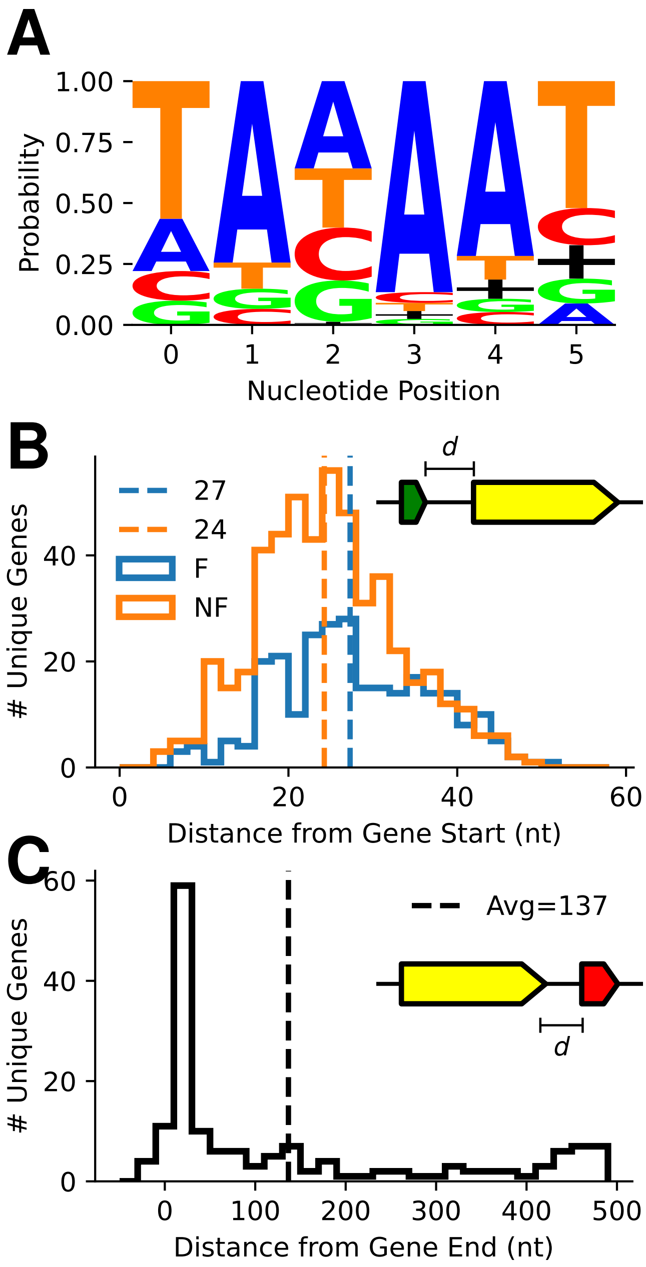


**Figure S13. Promoter and Termination results for** *M. pneumoniae*: **(A)** Nucleotide probability for the SD sequences for each gene. As expected the observed sequences matches closely with the promoter template sequence, $5'$-*TANAAT*-$3'$. Black plus signs indicate where no nucleotides is assigned due to motif sequence being found near the boundary of the search region. **(B)** Distance from gene start to the identified promoter sequences for functional (blue) and non-functional (orange) motifs. Results are in agreement with data from *M. florum* Matteau et al. (2020). Functionality is determined using the scores and a comparison to non-target search of the genome. (See **Section** 4.1.3). **(C)** Distance in nucleotide from the gene end to the start of the intrinsic termination loop. Values agree with reported numbers for *M. pneumoniae* in de Hoon et al. (2005). Validation shown in **Fig.** S14D.

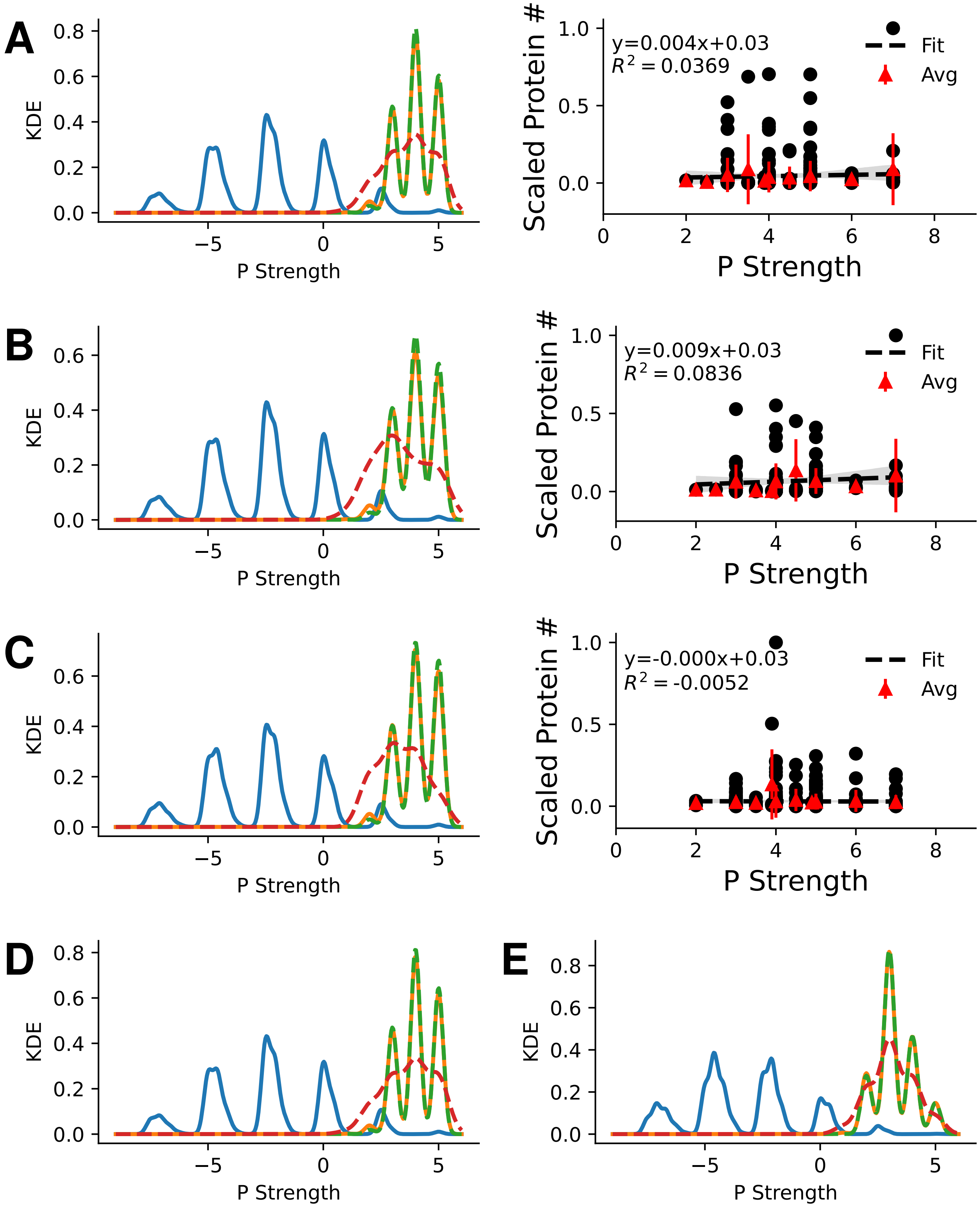


**Figure S14. Validation of Promoters**: Results of the non-specific promoter search within Syn1.0 **(A)** (left), Syn3A **(B)** (left), *M. florum* **(C)** (left), *M. mycoides* subsp. *capri* **(D)**, and *M. pneumoniae* **(E)**. Non-specific promoter strengths (blue) show lower values on average than the targeted search for promoters upstream of genes (orange). Protein coding (green) and non-coding genes (red) are separated, but unlike the SD show similar results to each other, which is expected because all are transcribed. When proteomics data was available the linear relationship between promoter strength and protein abundances was measured for functional promoters only. Results show no correlation for Syn1.0, Syn3A, and *M. florum* (**A, B, and C**, right). Proteomic data was obtained from this study (Syn1.0), Breuer et al. (2019) (Syn3A), and Matteau et al. (2020) (*M. florum*). Related to **Figs.** S9-S13

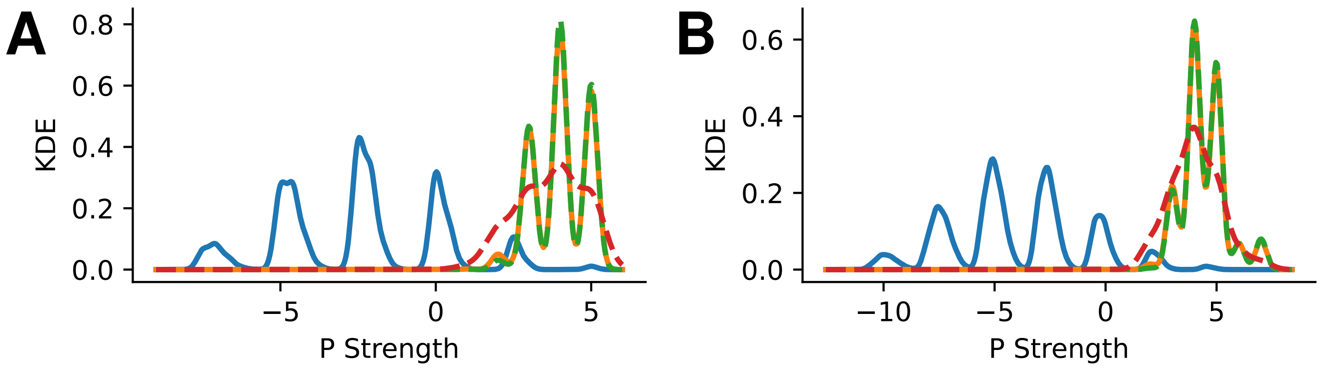


**Figure S15. Consensus Promoter vs Extended Promoter**: Comparison of the promoter strengths for the two different promoters: (A) the consensus sequence ($5'$-*TANAAT*-$3'$) and (B) the extended promoter sequence ($5'$-*TGNTANAAT*-$3'$). Data is shown for Syn1.0.

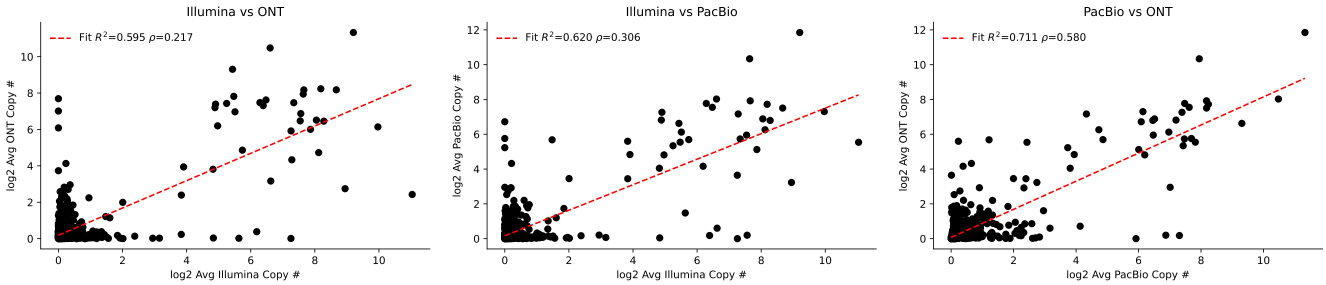


**Figure S16. RNA Copy # Correlation**: Comparison between RNA copy numbers observed in the RNAseq methods. Values are shown with black markers and a linear fit with a red dotted line. Pearson's ($R^2$) and Spearman's rank ($\rho$) correlation coefficient values have been reported in graph legend. RNA copy calculation is outlined in **Section** 4.7. Absolute copy number values have been averaged among replicates and then log transformed to account for sample size differences.

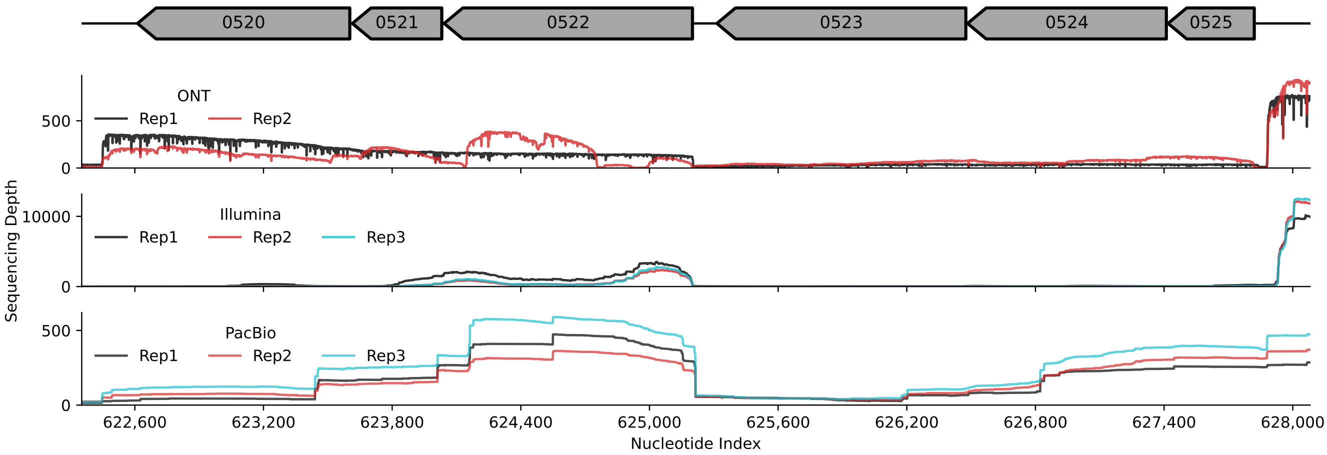


**Figure S17. DCW Operon Sequencing Depth:** Sequencing depth measured in ONT (top), Illumina (mid), and PacBio (bot) RNA sequencing experiments in Syn1.0. Data shows inconsisent transcriptional activity across the operon. PacBio results may be exaggerated due to the lack of strand specificity within the PacBio methods, however data still shows inconsitent gene expression. Transcriptional activity is observed by measuring a non-zero sequencing depth indicating a transcript was sequenced and mapped back to the genomic region.

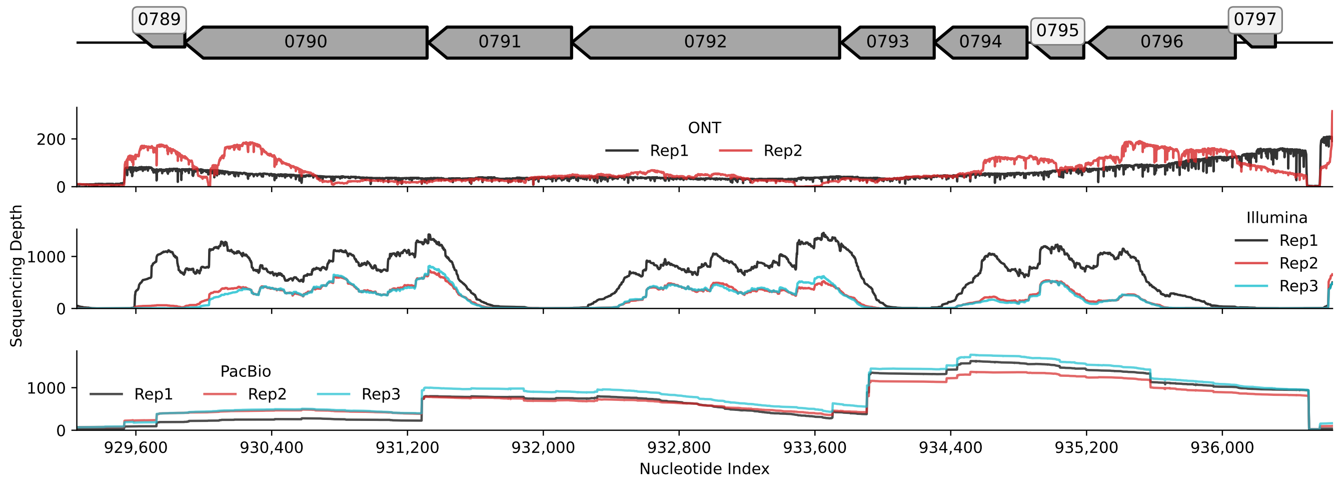


**Figure S18. ATP Synthase Operon Sequencing Depth:** Sequencing depth measured in ONT (top), Illumina (mid), and PacBio (bot) RNA sequencing experiments in Syn1.0. Data shows inconsisent transcriptional activity across the operon. PacBio results may be exaggerated due to the lack of strand specificity within the PacBio methods, however data still shows inconsitent gene expression. Transcriptional activity is observed by measuring a non-zero sequencing depth indicating a transcript was sequenced and mapped back to the genomic region.

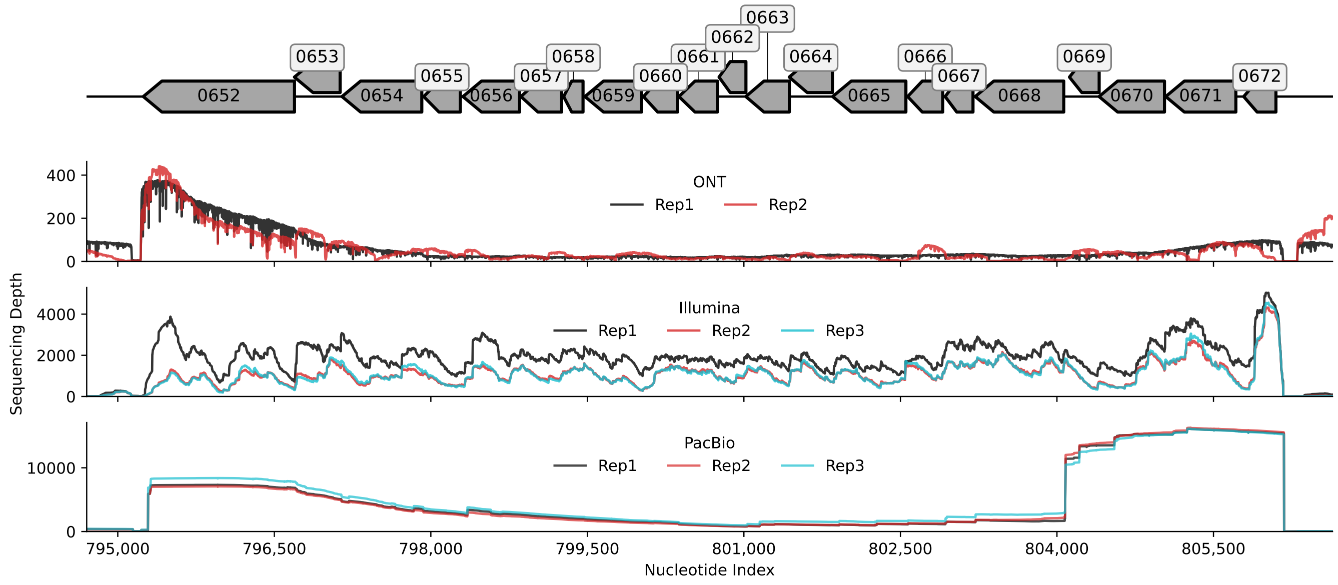


**Figure S19. Ribosomal Protein Operon Sequencing Depth:** Sequencing depth measured in ONT (top), Illumina (mid), and PacBio (bot) RNA sequencing experiments in Syn1.0. Data shows inconsisent transcriptional activity across the operon. PacBio results may be exaggerated due to the lack of strand specificity within the PacBio methods, however data still shows inconsitent gene expression. Transcriptional activity is observed by measuring a non-zero sequencing depth indicating a transcript was sequenced and mapped back to the genomic region.

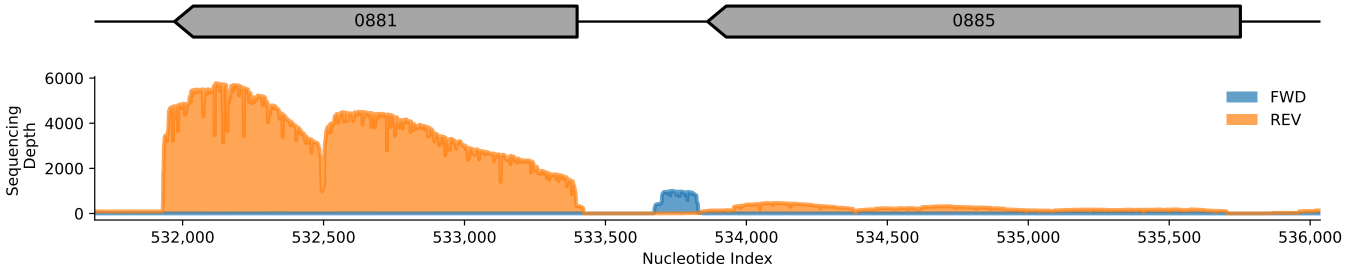


**Figure S20. Intergenic Transcription Activity in JCVI-syn3A**: Showing the detection of intergenic expression within a region unassigned to a gene ORF in Syn3A. Sequencing depth measure in single replicate in Syn3A, where non-zero values indicate transcription activity observed via the ONT RNAseq experiment. Forward strand is shown in blue and reverse strand in orange. Related to **Fig.** 3.

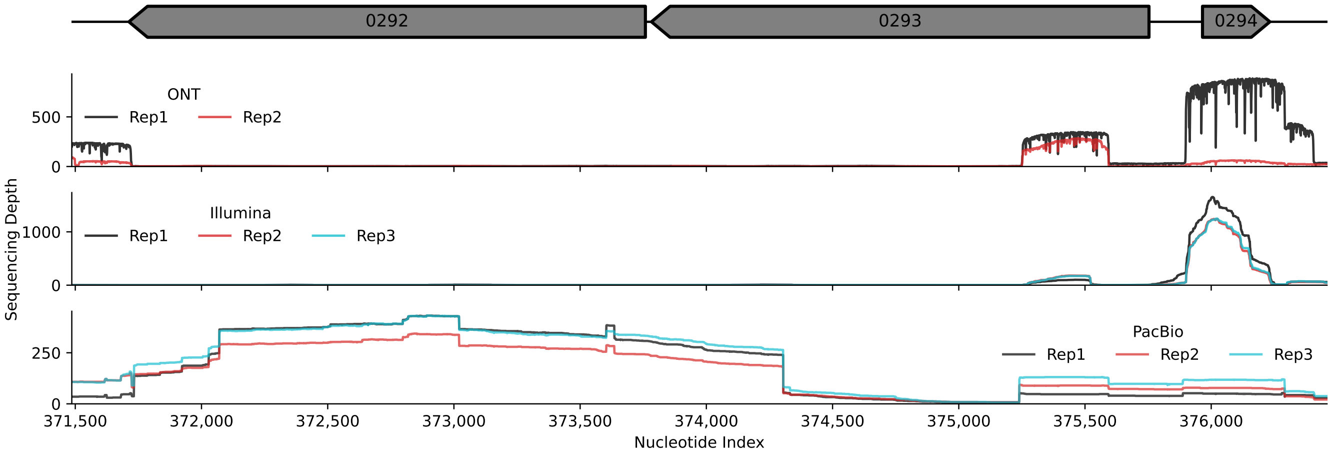


**Figure S21. Anti-sense Transcriptional Activity**: Sequencing depth measured in ONT (top), Illumina (mid), and PacBio (bot) RNA sequencing experiments in Syn1.0. Illumina and ONT data is shown for the forward strand (strand specific), and PacBio is strand agnostic. Each replicate within each data set reveals transcriptional activity near the $5'$ end of MMSYN1_0293 contrary to the fact the gene is on the reverse strand. Transcriptional activity is observed by measuring a non-zero sequencing depth indicating a transcript was sequenced and mapped back to genomic region. The observed activity is result of anti-sense transcription (asRNA formation), which was observed previously in Lloréns-Rico et al. (2016).

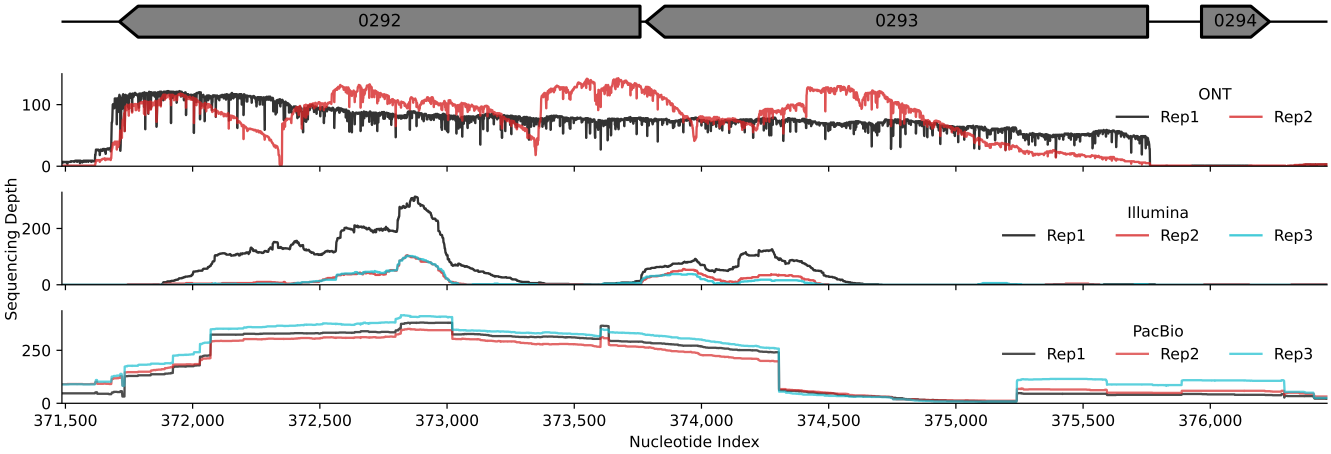


**Figure S22. Validation of Anti-sense Transcriptional Activity**: Sequencing depth measured in ONT (top), Illumina (mid), and PacBio (bot) RNA sequencing experiments in Syn1.0. Illumina and ONT data is shown for the reverse strand (strand specific), and PacBio is strand agnostic. The reverse strand data validates expression within MMSYN1_0293 which differs from the forward (anti-sense) strand expression within that gene. Related to **Fig.** S21

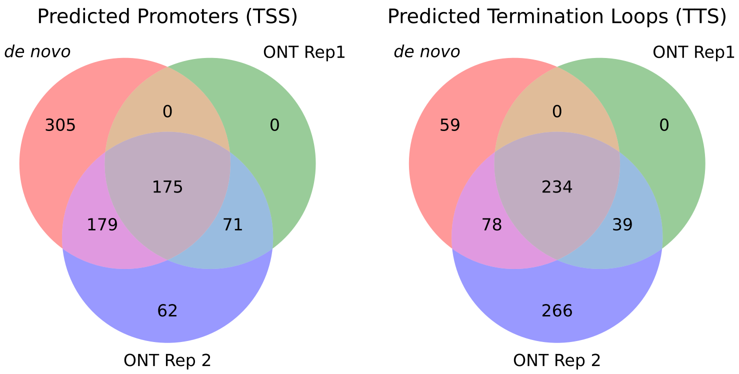


**Figure S23. *De novo* vs RNAseq-based TSS and TTS Prediction**: Venn diagrams showing the comparison of TSS and TTS predictions based on the bioinformatic motif identification (**Section** 2.1) and ONT RNAseq experiments (**Section** 2.3). Numbers indicates how many genes' TSS/TTSs were found in one of the datasets (outer ring values), two of the datasets (inner ring values), or all of the datasets (middle value). All TSSs and TTSs found in ONT Rep1 were found in either the bioinformatic predictions or the other ONT replicate. The resulting TUs generated by these results are shown in **SI File 4**.

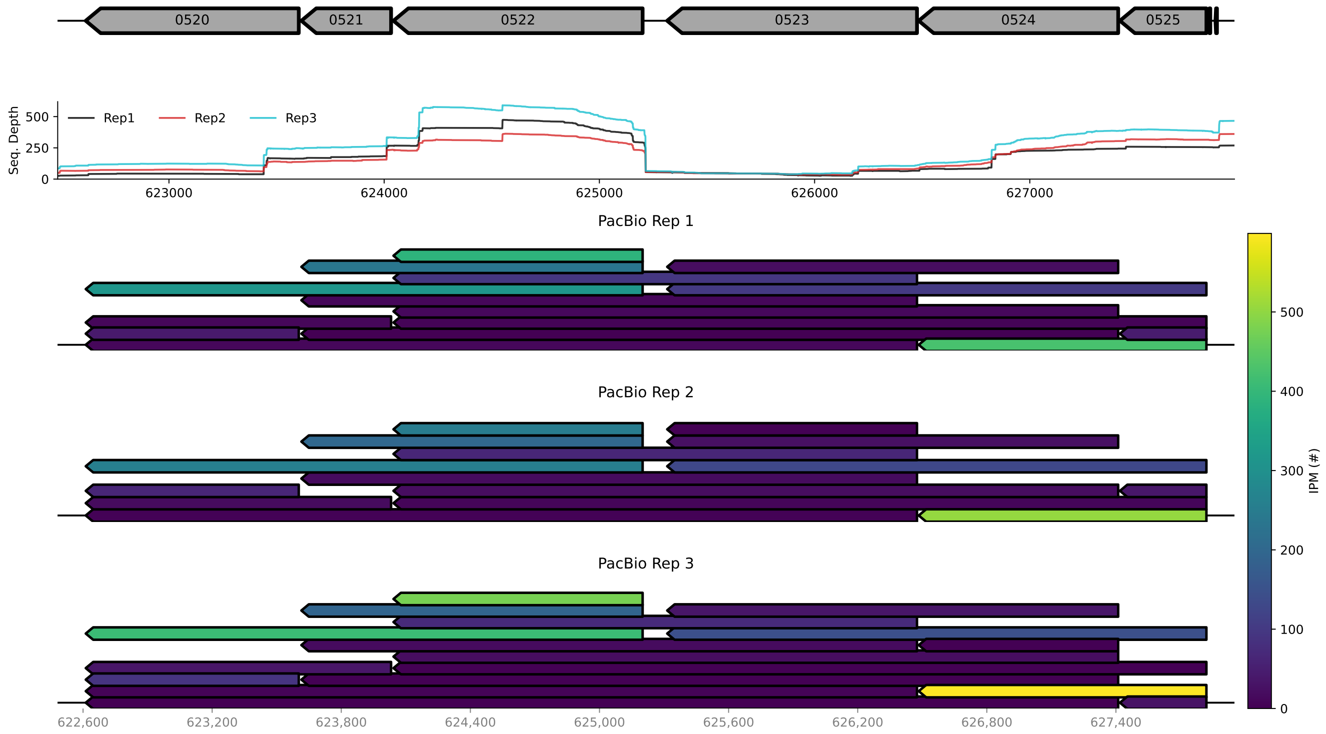


**Figure S24. DCW "Operon" RNA Isoforms**: (top) PacBio RNAseq derived isoforms from the Division and Cell Wall (DCW) genomic region. (mid) Traces of the PacBio sequencing depth across the region. Large changes in sequencing depth reflect differences in transcription activity between the two sub-regions: (i) genes 0520, 0521, and 0522 and (ii) genes 0523, 0524, and 0525. The first sub-region show relatively higher transcriptional activity. (bot) RNA isoforms colored by absolute abundances according to observations in the RNAseq data. RNA isoforms are drawn with lengths overlapping their associated genes. Related to **Fig.** 5.

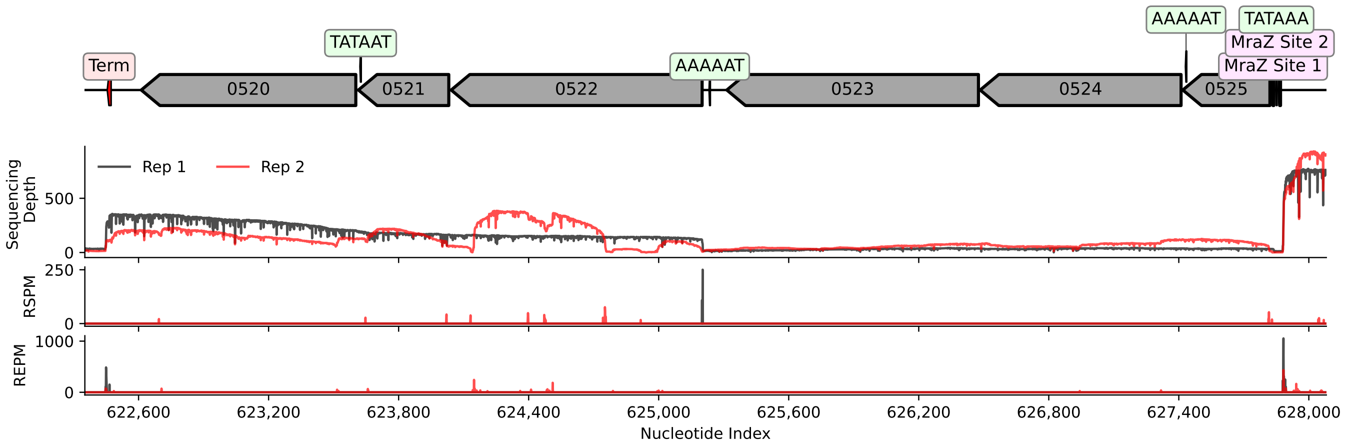


**Figure S25. DCW "Operon" TSS and TTS**: (top) Shows the genome sequence architecture for the genomic region. Bioinformatic predictions for the intrinsic terminators (red) and promoters with sequence (green) are shown along with a MraZ transcription factor binding sites upstream of *mraZ*/0525. The MraZ sites have consensus sequence $5'$-*AAAGTG[G/T]NNN*-$3'$ and are observed to repress transcription of the region Fisunov et al. (2016).(bot) Prediction of the TSS and TTS from the ONT RNAseq data determined by RSPM and REPM, respectively. Sites are identified by peaks within the REPM and RSPM traces, where the larger the value the greater the frequency of observed initiation/termination events. There is relative agreement between the bioinformatic and RNAseq predictions. Related to **Fig.** 5.

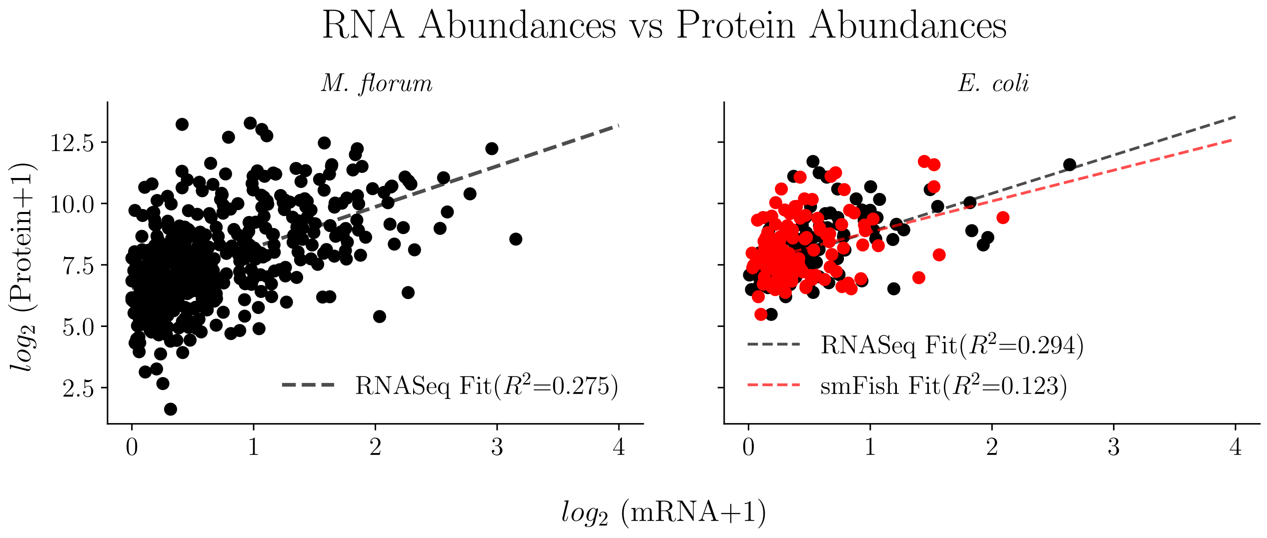


**Figure S26. mRNA vs Protein Abundances Related Organisms:** Relationship between standard-scaled abundances measurements of mRNA and its encoded protein for two related organisms: *Escherichia coli*, and *Mesoplasma florum*. Dashed lines reflect a linear fit to the data. Coefficient of determination, $R^2$, are provided of each fit. mRNA and proteomic data is obtained from *E. coli* Taniguchi et al. (2010) (RNAseq results) and *M. florum* Matteau et al. (2020).

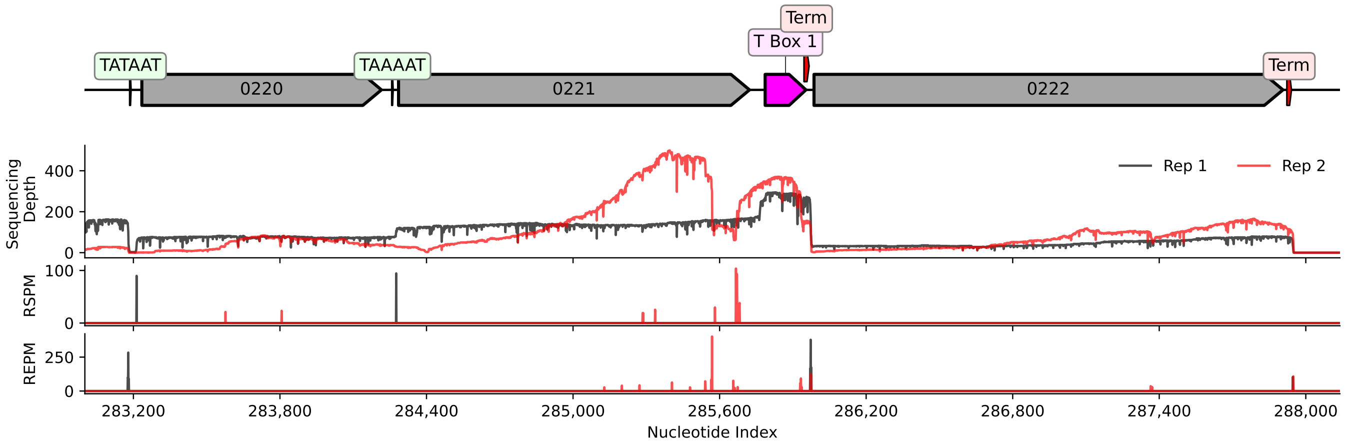


**Figure S27. T Box Riboswitch I Region TSS and TTS Predictions**: (top) Shows the genome sequence architecture for the T Box Riboswitch I genomic region. Bioinformatic predictions for the intrinsic terminators (red) and promoters with sequence (green) are shown as well as the riboswitch (magenta). (bot) Prediction of the TSS and TTS from the ONT RNAseq data determined by RSPM and REPM, respectively. Sites are identified by peaks within the RSPM and REPM traces, where the larger the value the greater the frequency of observed initiation/termination events. The TTS sites show strong termination signal at the predicted intrinsic termination loop, thus the riboswitch is primarily in an inactive. The TSS sites agree well with the bioinformatic promoter predictions. Related to **Fig.** 6.

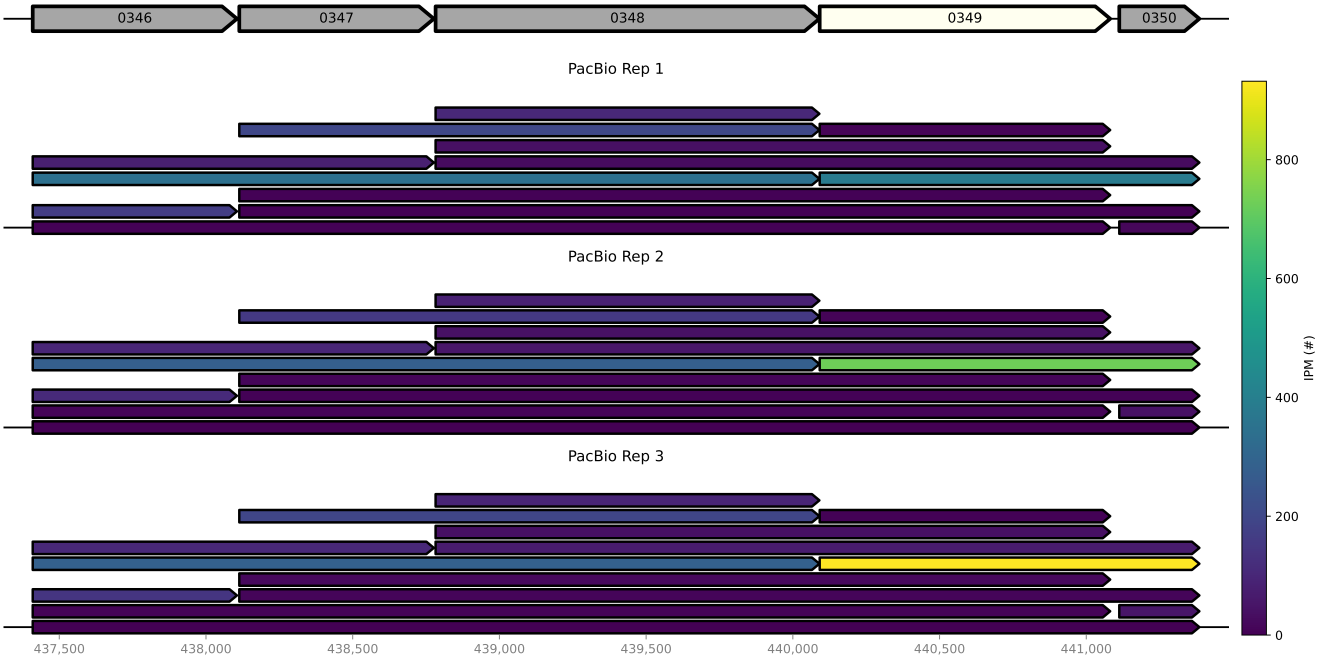


**Figure S28. HupA RNA Isoforms**: PacBio perspective of the RNA isoforms for the *hupA*/0350 region. Related to **Fig.** 8C. RNA isoforms are drawn with lengths overlapping the genes they are associated with. RNA isoforms are colored by absolute abundances according to observations in the RNAseq data. *hupA*/0350 and *gpsA*/0349 are commonly found encoded within a single RNA isoform. Therefore the perturbation of the genome sequence architecture via the removal of *gpsA*/0349 is likely to have resulted in the significant (~$10^2$) drop in the protein expression of *gpsA*/0350 from Syn1.0 toSyn3A.

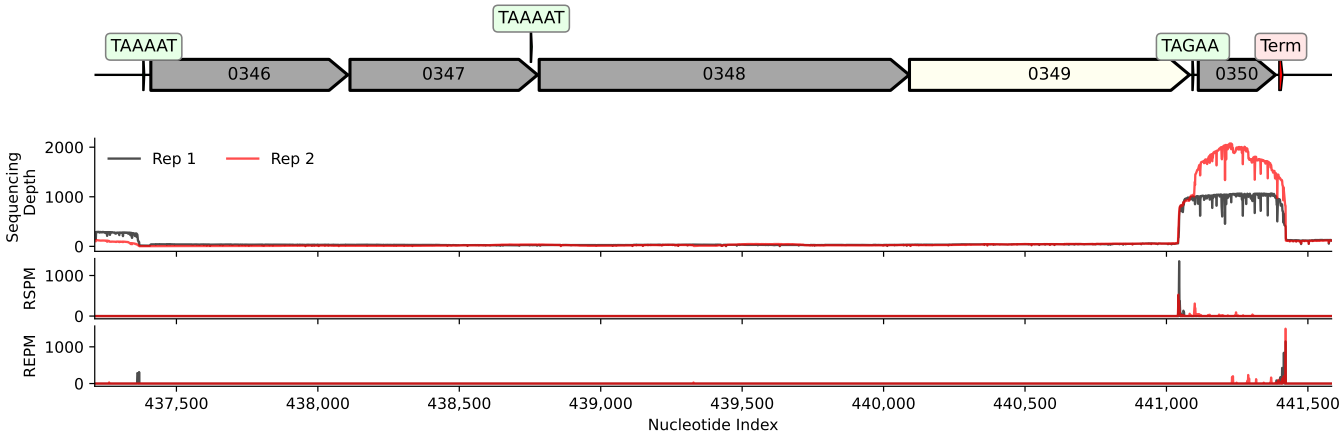


**Figure S29. HupA TSS and TTS Predictions**: (top) Shows the genome sequence architecture for the HupA genomic region with the bioinformatic predictions for the intrinsic terminators (red) and promoters with sequence (green) shown. (bot) Prediction of the TSS and TTS from the ONT RNAseq data determined by RSPM and REPM, respectively. Sites are identified by peaks within the RSPM and REPM traces, where the larger the value the greater the frequency of observed initiation/termination events. A strong TSS site is observed within the $5'$ end of *gpsA*/0349, which is likely related to the predicted promoter upstream of *hupA*/0350. Additionally, the strong TTS site in the RNAseq agrees with the termination loop prediction. Related to **Fig.** 8.

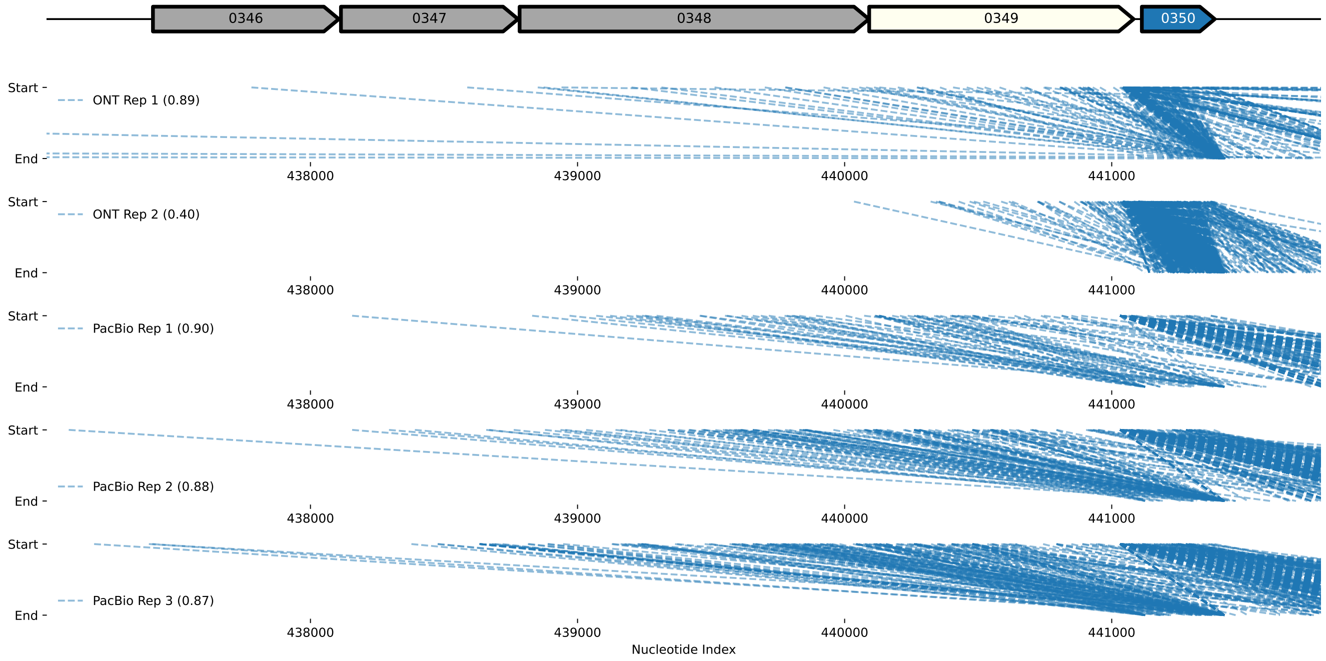


**Figure S30. HupA Isoform Genomic Positions**: (top) Shows the genome sequence architecture for the HupA (blue) genomic region. *gpsA*/0349 (white), which was removed during genome reduction to Syn3A, is shown in by a white gene ORF. (bot) Start and end positions for the RNA isoforms assigned to *hupA*/0350 observed in each of the long-read RNAseq experiments. Data consistently shows most transcripts associated with messengers encoding HupA start within the *gpsA*/0349 gene ORF (nearly 90% for all replicates). Related to **Fig.** 8.

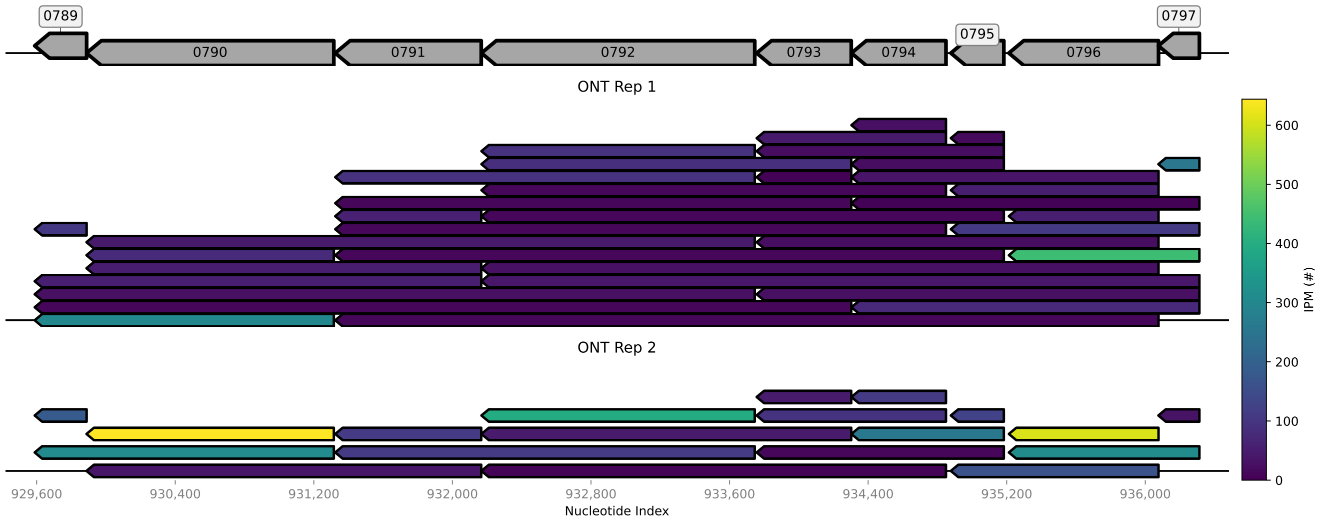


**Figure S31. ATP Synthase Motifs and RNA Isoforms**: (top) Predicted Shine-Dalgarno (cyan), promoters (green), intrinsic termination loops (red) for the ATP synthase operon. Shine-Dalgarnosequences are denoted with strength values from predictions (max value of 10), while promoters are shown with identified sequence. (bot) ONT derived RNA isoforms drawn overlapping their associated genes. RNA isoforms are colored by absolute abundances according to observations in the RNAseq data. The ATP synthase complex has sub-unit stoichiometries of 1:3:1:3:1:2:10:1 respectively from *atpC*/0789 to *atpB*/0796 Sobti et al. (2020). Functional SDs, which are commonly found within the RNA isoforms, may explain give insight into how differential translation resulting from varied translation initiation efficiencies gives rise to the non-trivial stoichiometry of the ATP synthase complex.

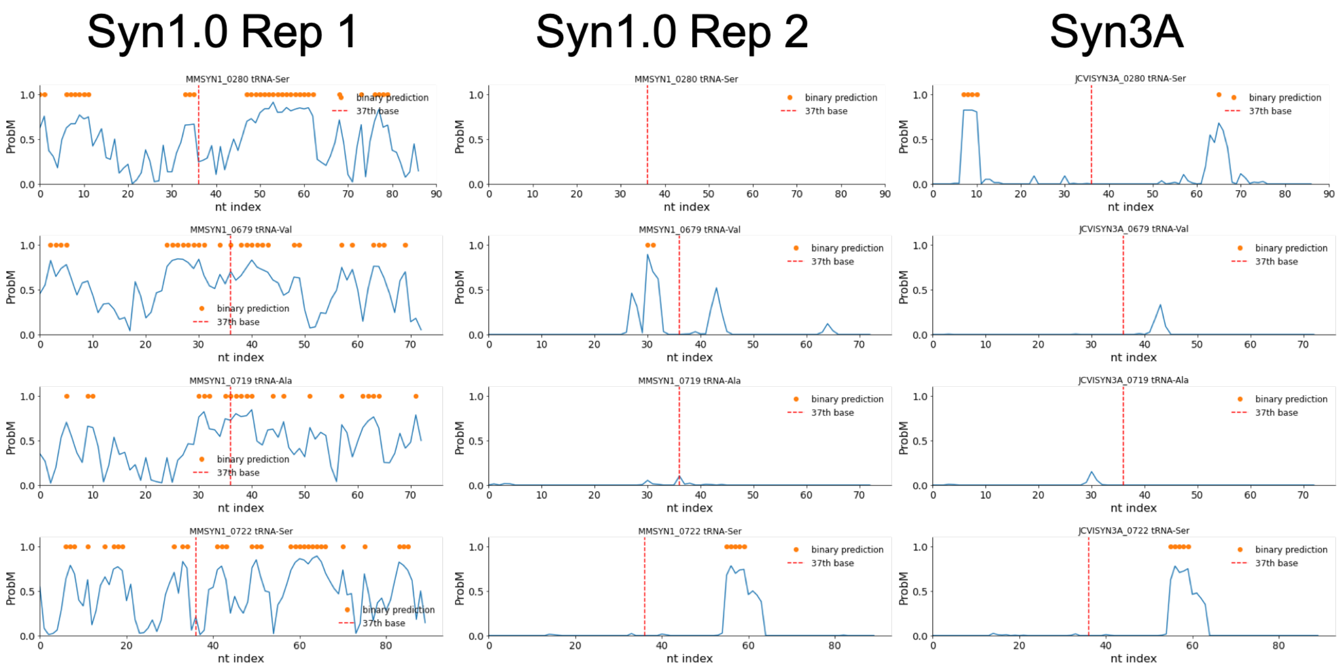


**Figure S32. tRNA m$^6$A Prediction:** Probability (blue) of a m$^6$A modification detected using the EpiNano Liu et al. (2019, 2021). Orange markers denote a modified base according to the EpiNano algorithm. Red dotted lines highlight the 37$^{th}$ nucleotide in the tRNA sequence, the typical location of the m$^6$A modification (Val and Ala is at 37, VAL at 40 Samuelsson et al. (1987)). Serine (MMSYN1_0280) Syn1.0 rep2 shows no blue trace as the coverage in that region of the genome for that replicate did not allow for proper investigation with the EpiNano alogithm.

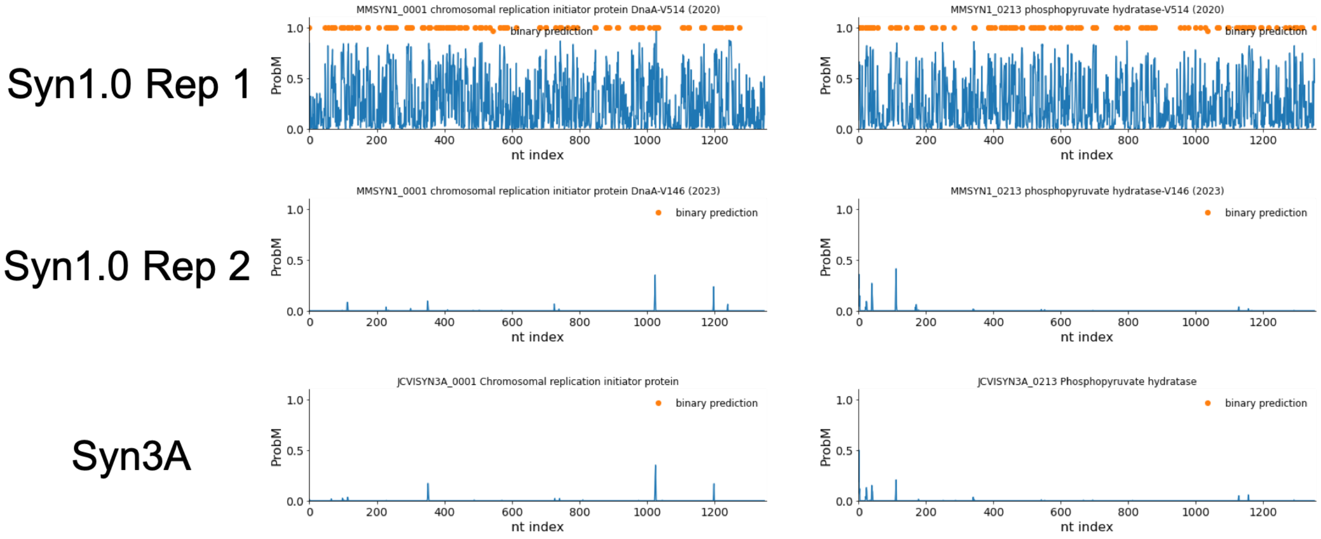


**Figure S33. mRNA m$^6$A Prediction:** Probability (blue) of a m$^6$A modification detected using the EpiNano Liu et al. (2019, 2021). Orange markers denote a modified base according to the EpiNano algorithm. Two mRNA were chosen: *0001*/dnaA – chromosome replication initiation protein and *0213*/eno – enolase.

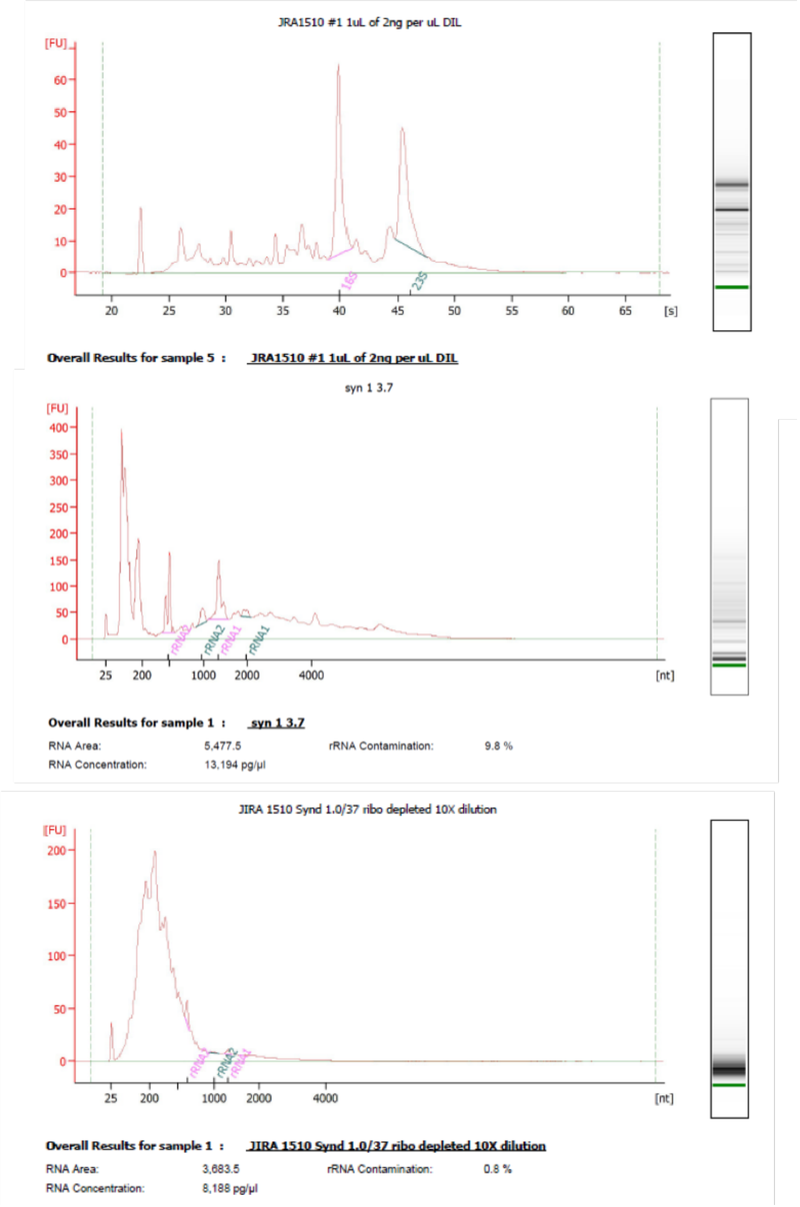


**Figure S34. ONT RNA Quality Control**: RNA traces for quality assessment of initial JCVI-syn1.0 sample using an Agilent Bioanalyzer 2100 for Oxford Nanopore Direct RNA Sequencing. (top) Total RNA before ribosomal RNA (rRNA) depletion depletion using an Invitrogen RiboMinus$^{TM}$ Bacteria 2.0 Transcriptome Isolation Kit. (mid) After one round of rRNA depletion the sample contained 9.8% rRNA. (bot) After a second cycle rRNA was reduced to 0.8%. Our threshold for maximum rRNA content was $\leq$3%. Related to **Section** 4.3.2.

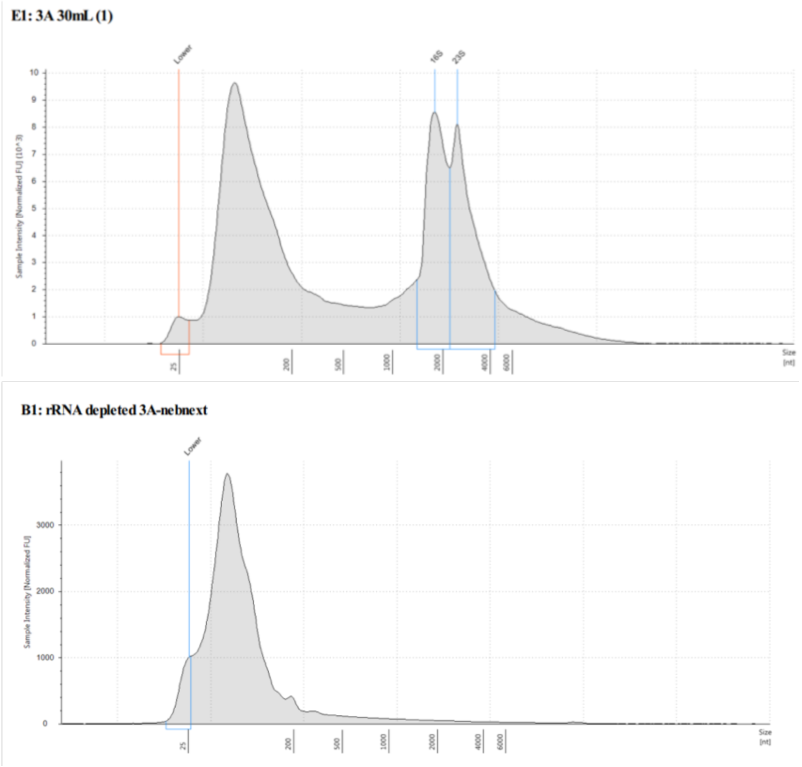


**Figure S35. ONT RNA Quality Control**: RNA traces for quality assessment of JCVI-syn3A samples using an Agilent TapeStation 2200 for Oxford Nanopore Direct RNA Sequencing. (top) Total RNA before ribosomal RNA (rRNA) depletion using a NEBNext® rRNA Depletion Kit for Bacteria. (bot) After rRNA depletion using Our threshold for maximum rRNA content was $\leq$3%. Related to **Section** 4.3.2.

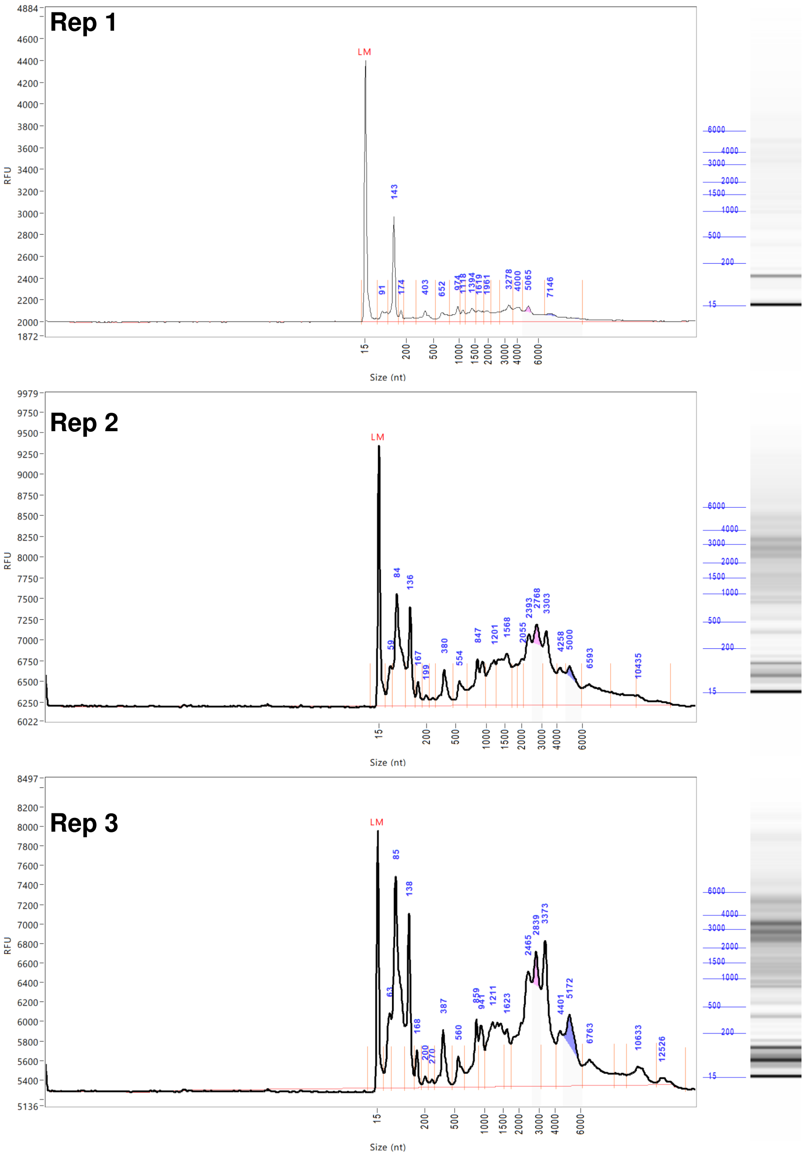


**Figure S36. Illumina RNA Quality Control**: RNA traces for quality assessment of library sample preparation related to **Section** 4.4.2.

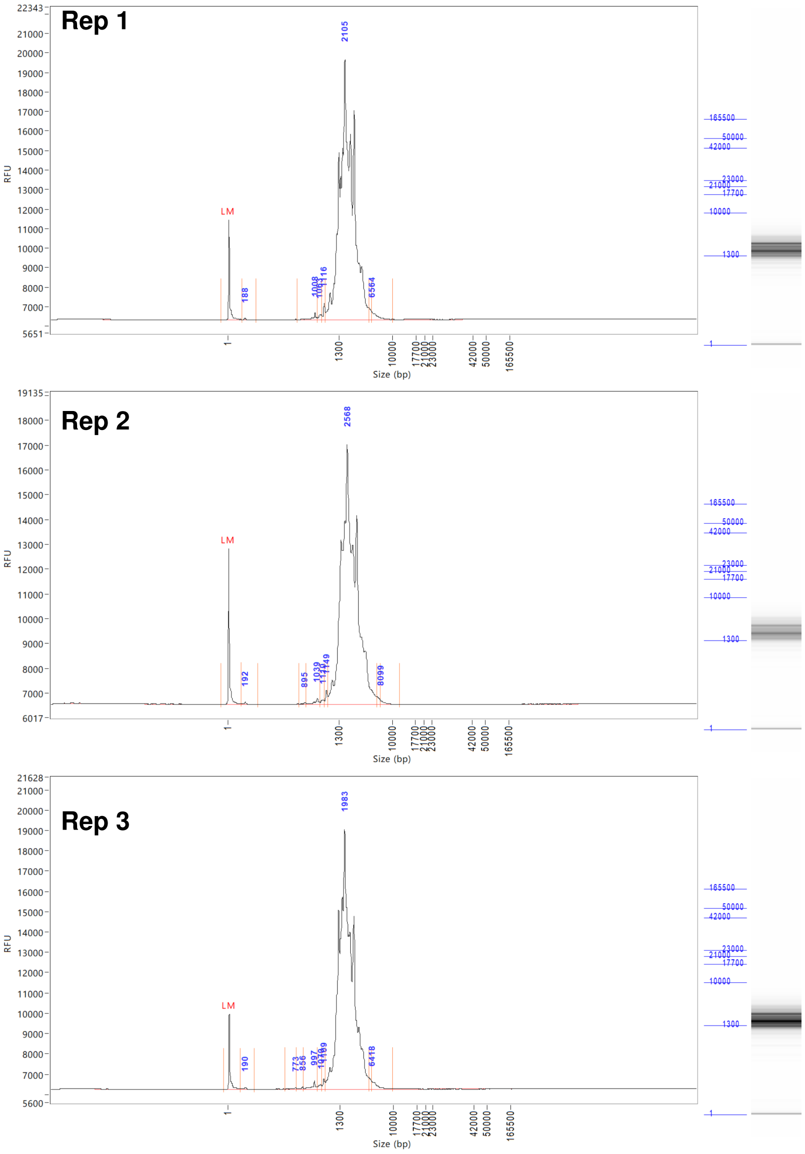


**Figure S37. PacBio RNA Quality Control**: RNA traces for quality assessment of library sample preparation related to **Section** 4.5.2.
